# Ultrasound-Driven Programmable Artificial Muscles

**DOI:** 10.1101/2024.01.08.574699

**Authors:** Zhan Shi, Zhiyuan Zhang, Justus Schnermann, Stephan C. F. Neuhauss, Nitesh Nama, Raphael Wittkowski, Daniel Ahmed

## Abstract

Muscular systems^1^, the fundamental components of mobility in animals, have sparked innovations across technological and medical fields^2,3^. Yet artificial muscles suffer from dynamic programmability, scalability, and responsiveness due to complex actuation mechanisms and demanding material requirements. Here, we introduce a design paradigm for artificial muscles, utilizing >10,000 microbubbles with targeted ultrasound activation. These microbubbles are engineered with precise dimensions that correspond to distinct resonance frequencies. When stimulated by a sweeping-frequency ultrasound, microbubble arrays in the artificial muscle undergo selective oscillations and generate distributed point thrusts, enabling the muscle to achieve programmable deformation with remarkable attributes: a high compactness of ∼3,000 microbubbles/mm², a low weight of 0.047 mg/mm^2^, a substantial force intensity of ∼1.21 μN/mm^2^, and fast response (sub-100 ms during gripping). Moreover, they offer good scalability (from micro- to -centimeter scale), exceptional compliance, and many degrees of freedom. We support our approach with a theoretical model and demonstrate applications spanning flexible organism manipulation, conformable robotic skins for adding mobility to static objects and conformally attach to ex vivo porcine organs, and biomimetic stingraybots for propulsion within ex vivo biological environments. The customizable artificial muscles could offer both immediate and long-term impact on soft robotics, wearable technologies, haptics, and biomedical instrumentation.

## Main

Flexible, compact, and adaptive artificial muscles are set to be transformative across multiple fields, including soft robotics^4,5^, wearables for human-machine interactions and healthcare, like prosthetics^6^, orthotics^7^, and embodied sensing^8,9^, and assistance in sophisticated manufacturing through dexterous manipulation^10,11^. In biomedicine, they could revolutionize soft surgical tools^12^, implantable electrodes^13^, and artificial organs like the heart^14^. Despite their potential, current artificial muscles like tendon-based^15^ and pneumatic types^16^ encounter substantial challenges in wireless control, integration, and miniaturization due to dependencies on tethering, complex operational mechanisms, and large input requirements. While external stimuli such as chemicals^17^, light^18,19^, temperature^20–22^, electric fields^23–25^, and magnetic fields^26,27^ have been deployed for wireless actuation, they face challenges in biocompatibility, spatial resolution, and dynamic programmability. Chemical methods often require fuels that could be toxic^28^, light-based systems suffer from limited tissue penetration and potential thermal damage^29^, and magnetic systems necessitate bulky hardware while risking Joule heating^30^. In contrast, acoustic actuation emerges as a promising biocompatible alternative. It offers a material-independent and simplified design, enabling wireless control, remote deployment, millisecond-scale responsiveness, multimodal programmability, high spatial selectivity, and deep tissue penetration—all without invasive hardware. Moreover, its compatibility with existing clinical ultrasound devices and imaging systems makes it particularly uniquely suited for in vivo use and broader biomedical applications^31–38^.

Central to this approach are resonant microbubbles, which concentrate acoustic energy and enable weak ultrasound sources to generate amplified responses. While prior ultrasound-actuated microrobots and actuators have employed single or sparse microbubbles embedded in polymers to achieve basic propulsion^39–41^, their functionality remained limited. Directional steering has been demonstrated through strategies such as tuning microbubble sizes^42^, applying magnetic navigation^43^, or hybrid methods that combine magnetic fields with asymmetric appendages in encapsulated shells^44^. An actuator composed of a microbubble attached to a flexible beam was developed to analyze the kinematic behavior of some simple microstructures through the excitation of different pairs of bubble actuator modules^45^. Another study utilized arrays of microbubbles integrated onto centimeter-scale rigid substrates to induce bi-rotational motion^46^, demonstrating potential applications in endoscope design^47^. However, these systems lack the programmability, scalability, and dynamic adaptability required to emulate natural muscle behavior. Critically, to the best of our knowledge, no prior work has achieved ultrasound-actuated soft artificial muscles, marking a significant gap in biologically inspired actuation technologies.

Why did ultrasound-based artificial muscles remain undeveloped? Soft materials typically exhibit low acoustic contrast factors compared to water, leading to inadequate force generation for efficient functionality when activated by ultrasound. This inadequacy becomes more pronounced at larger scales, which requires lower frequencies and thus lower damping to match their increased dimensions. This predicament is exacerbated by a lack of understanding of the interactions between sound and complex soft materials, impeding the progress of effective sound-driven muscle systems. However, we found that integrating ultrasound-activated microbubble arrays into soft artificial muscles presents a clever approach that could potentially address these limitations.

Here, we introduce an artificial muscle built on acoustically activated microbubble arrays. This synthetic muscle comprises a thin, transparent, and flexible membrane that houses over 10,000 microcavities arranged in arrays, designed to confine microbubbles of various sizes. When these microbubbles are acoustically stimulated, they generate thrust, causing the membrane to deform. Tailored activation of differently sized microbubble arrays through programmable sweeping-frequency ultrasound excitation results in localized point forces, allowing dynamic multi-modal deformation of the artificial muscle. The tunable nature and scalability of these microbubble arrays herald a new era of possibilities, positioning these acoustic artificial muscles at the forefront of innovation in robotics, wearable technology, prosthetic development, and soft surgical devices.

## Results

### Design and fabrication of microbubble-array artificial muscles

In the initial design of the ultrasound-driven artificial muscle (**Fig. 1a**), we incorporated uniform-size microcavities on the muscle’s bottom surface. When the muscle was submerged in an acoustic chamber filled with water, it resulted in the simultaneous trapping of tens of thousands of gas-filled microbubbles within these cavities, a phenomenon driven by surface tension. To test the muscle’s actuation, we anchored one end and left the other free, forming a cantilever configuration. Subsequently, we activated a piezoelectric transducer to generate ultrasound. The incident sound waves propagated through the liquid, triggering oscillations in the microbubbles. Since all microbubbles in the muscle were of identical dimensions, they were simultaneously excited. This harmonic bubble oscillation generated collective acoustic streaming and radiation forces, applying a uniform opposing force to the muscle’s bottom surface and resulting in upward flexion. By modulating the ultrasound excitation voltage, we controlled the deformation amplitude of the artificial muscle.

**Fig. 1.**
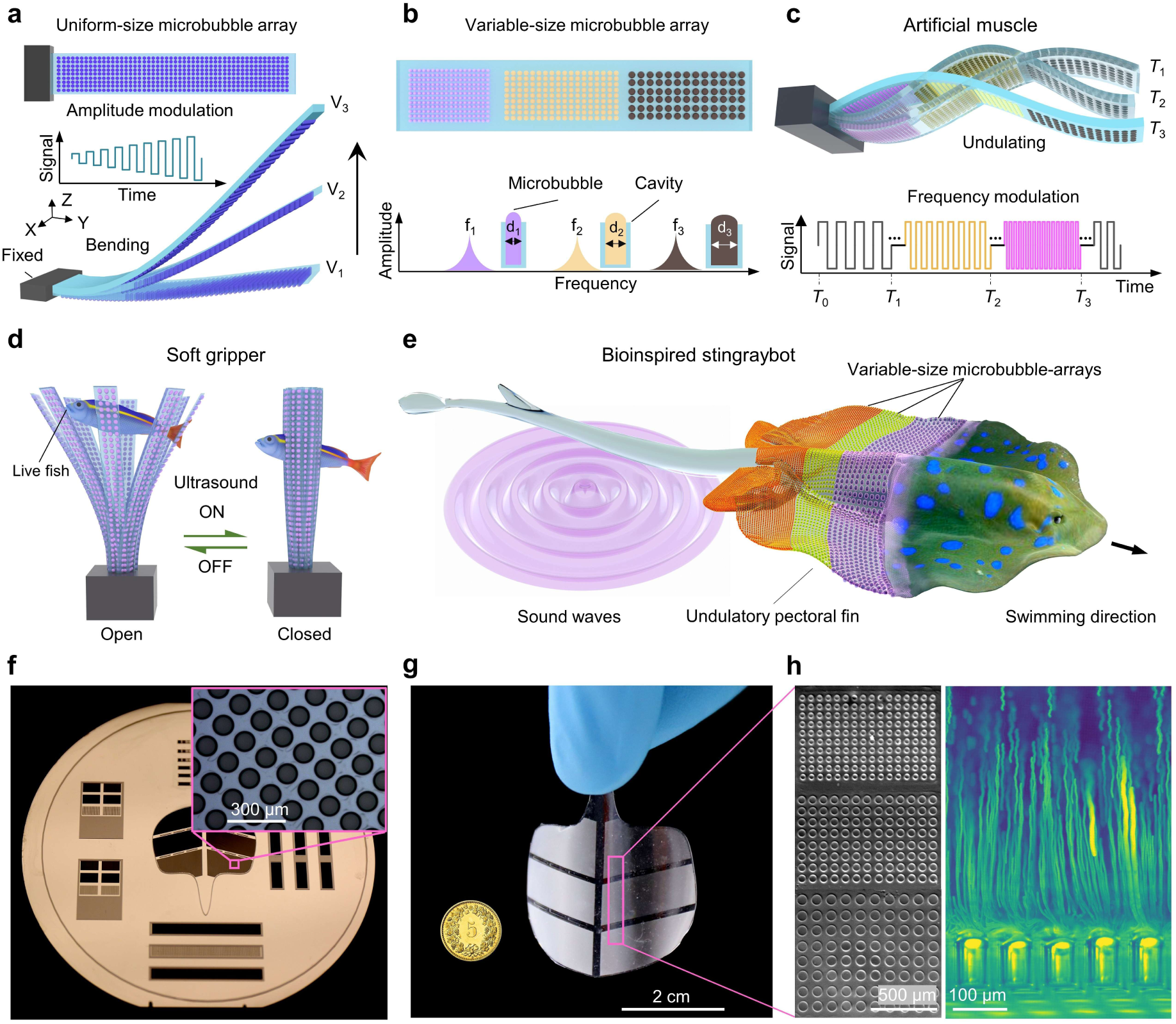
Ultrasound-actuated microbubble-array artificial muscles. **a**, A uniform-size microbubble-array artificial muscle consists of thousands of microbubbles on its bottom surface. Under continuous ultrasound excitation, the artificial muscle bends upwards with different excitation voltages, labeled as *V*_1_, *V*_2_, and *V*_3_. The inset shows the input ultrasound signal with modulated amplitude versus time. **b**, A variable-size microbubble-array artificial muscle consists of three microbubble arrays with different microbubble diameters featuring different natural frequencies represented by distinct colors. **c**, Under sweeping-frequency ultrasound excitation, the artificial muscle exhibits multi-mode deformation in the time domain, shown at time points *T*_1_, *T*_2_, and *T*_3_. **d**, Schematic of a soft gripper constructed with an array of artificial muscles patterned with uniform-size microbubble arrays. Upon ultrasound excitation, these muscles close simultaneously in milliseconds. **e**, Schematic of a bioinspired stingraybot incorporates variable-size microbubble array artificial muscles. Under sweeping-frequency ultrasound excitation, the stingraybot enacts undulating propulsion. **f**, A silicon wafer with micropillar arrays serves as the negative mold of microbubble cavities in standard soft-lithography fabrication. Inset shows the micropillar array. **g,** A prototype of the stingraybot. Inset shows a 5-cent Swiss franc coin. **h**, Left: Trapped microbubbles on the stingraybot. Right: Upward microstreaming jetting generated from a microbubble array oscillating under ultrasound excitation demonstrated by 6 μm diameter tracer microparticles.

We then designed an artificial muscle featuring microbubble arrays of varying bubble sizes, illustrated in **Fig. 1b**. Since microbubbles of different sizes exhibit distinct resonance frequencies, they can be independently activated to produce localized opposing forces and selective muscle deformation. By applying a sweeping-frequency ultrasound signal that encompasses the natural frequencies of all microbubbles, we sequentially activated distinct arrays along the muscle’s longitudinal axis. This orchestrated activation generated complex undulatory motion across multiple excitation cycles (**Fig. 1c**). Thus, in implementing this arrangement of microbubbles of varied sizes and their frequency-selective excitation through ultrasound modulation, we have unlocked a capability to control multi-modal deformations. The versatility of these artificial muscles facilitates a wide array of applications. For example, we implemented these artificial muscles in the development of a soft gripper, crafted to delicately handle live fish (**Fig. 1d**), and in the design of a soft swimmer, inspired by the form and function of stingrays (**Fig. 1e**), among other functional systems.

Prototypes of these artificial muscles were fabricated using a high-resolution mold replica method. First, micropillar arrays were patterned on a silicon wafer using soft lithography to serve as negative molds for cylindrical microcavities (**Fig. 1f**). All pillars were designed of identical heights and spacings, corresponding to dimensions of desired microbubbles (**Supplementary Fig. 1**). A thin layer of polydimethylsiloxane (PDMS) was then spin-coated onto the wafer, yielding uniform thicknesses ranging from 80 μm to 250 μm (**Supplementary Fig. 2**). After curing, these artificial muscles including the artificial stingray (**Fig. 1g)** were demolded, sectioned, and prepared for testing. Full fabrication details are provided in the Materials and Methods section. **Figure 1h** demonstrates microbubbles trapped on the “stingraybot” and the upward microstreaming jets produced during ultrasound excitation.

### Characterization of microbubble arrays

To advance our understanding and control of microbubble arrays in artificial muscles, we observed the transient dynamics of the microbubbles while applying acoustic fields using a high-speed camera with excitation frequencies ranging from 10−100 kHz and voltage amplitudes of 10−60 V_PP_ in square waveforms. Further details of the acoustic setup are provided in the **Methods** section.

We began by identifying the resonance frequency of microbubbles confined within cavities of different diameters (40−140 μm, in 10 μm increments) and depths (50 μm, 150 μm, and 175 μm) while maintaining a constant excitation voltage of 15 V_PP_. Resonance frequencies were identified by locating peak oscillation amplitudes during frequency sweeps (**Extended Data Fig. 1a)**. As shown in **Extended Data Fig. 1b**, resonance frequencies decreased from 95.5 kHz to 8.9 kHz with increasing microbubble diameter, consistent with the inverse scaling relationship between natural frequency and the bubble diameter^48^. Additionally, microbubbles with depths of 50 μm (purple line), 150 μm (green line), and 175 μm (blue line) exhibited a reduction in resonance frequency, indicating that bubble depth also affects oscillation. We further investigated the selective actuation of variable-size microbubble arrays of 40 μm, 60 μm, and 80 μm diameter, each 150 μm in depth, integrated within a single miniaturized artificial muscle (500 μm × 500 μm × 200 μm) with corresponding frequencies (27.6 kHz, 57.4 kHz, and 76.3 kHz, respectively), as demonstrated in **Extended Data Fig. 2** and **Supplementary Video 1**. The distinct resonance profiles of microbubbles across sizes enable selective ultrasound excitation, forming the basis for programmable microbubble arrays. Detailed microstreaming characterization is provided in the **Methods** section.

### Programmable actuation of microbubble-array artificial muscles

The versatility of microbubble arrays in terms of programmability and selectivity enables an innovative approach for designing soft actuators with enhanced flexibility and control. To verify that microbubble oscillation is the dominant driver of this muscle bending, we systematically varied the transducer’s position relative to the microbubble-embedded side of the artificial muscle (3 cm × 0.5 cm × 80 μm), which contains over 10,000 uniform microbubbles within cavities (40 μm diameter, 50 μm depth). The transducer was positioned with four distinct orientations: (1) directly facing the microbubble-embedded side, (2) opposite to it, and (3-4) perpendicular to the array’s left and right sides of the artificial muscle (**Supplementary Fig. 3**, and **Supplementary Video 2**). When activated at 80.5 kHz and 60 V_PP_, the muscle consistently bent away in the direction opposite to the microbubble-array side, across all configurations, despite variations in bending amplitudes. This directional uniformity confirms that microbubble-generated reverse thrust is the primary force driving the deformation. More control experiments and characterization of artificial muscle deformation are provided in the **Methods** section.

To demonstrate the selective excitation capability of the artificial muscle, we further investigated the deformation of an artificial muscle equipped with variable-size microbubble arrays. The muscle, measuring 3 cm × 0.5 cm × 80 μm, contains three arrays of microbubbles with diameters of 12, 16, and 66 μm, each with a depth of 50 μm. Upon stimulation at its resonance frequency (96.5 kHz), the 12 μm × 50 μm microbubble array, covering an area of 0.5 cm², induced a leftward deformation in the corresponding muscle region, as depicted in **Fig. 2a** and **Supplementary Video 3**. Similarly, when the frequency was respectively increased to match the resonance frequencies of the 16 μm (82.3 kHz, **Fig. 2b**) and 66 μm (33.2 kHz, **Fig. 2c**) bubble arrays, the muscle showed a localized leftward deformation in the middle region and bottom region, respectively. We further demonstrated an undulatory sinusoidal-like deformation by actuating the artificial muscle with a sweeping-frequency ultrasound excitation (20 to 90 kHz). This continuous, time-dependent motion, as shown in **Fig. 2d** and **Supplementary Video 4**, resulted from the periodic reverse thrust generated across different regions of the muscle.

**Fig. 2.**
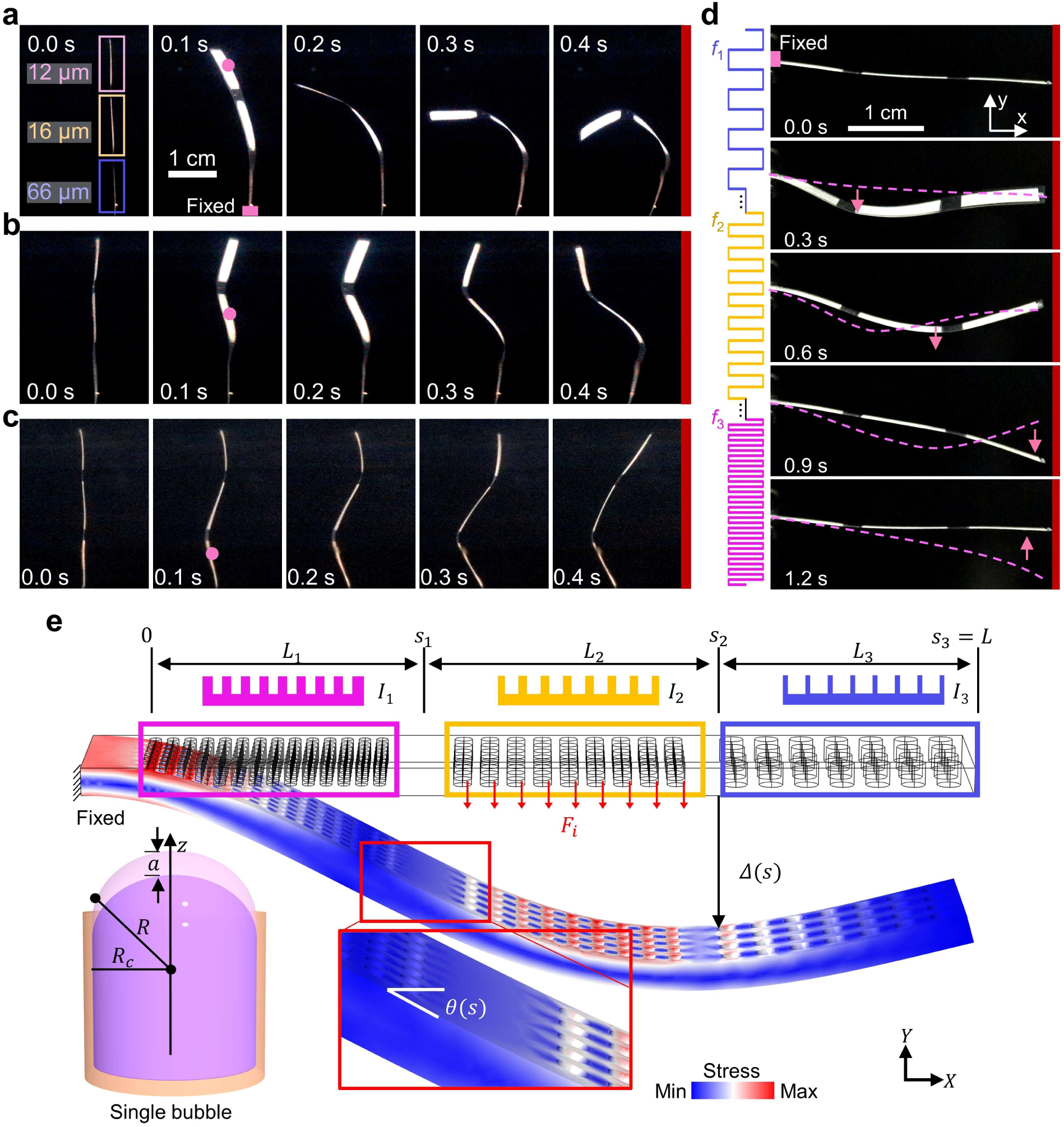
Actuation and modeling of microbubble-array artificial muscles. a to c, Time-lapse images of the individual deformation shapes of a variable-size microbubble array artificial muscle (3 cm × 0.5 cm × 80 μm) with microbubble diameters of 12 μm, 16 μm, and 66 μm, excited at 96.5 kHz, 82.3 kHz and 33.2 kHz, respectively. After 0.4 s the deformation will not change significantly. The microbubbles have a constant depth of 50 μm. The pink dots indicate the region of the bubble array that is activated. **d**, Time-lapse images of the variable-size microbubble array artificial muscle under sweeping-frequency ultrasound excitation (20-90 kHz, 1.2 s). The ultrasound excitation voltage for panels **a** to **d** was consistently at 60 V_PP_. The pink dashed lines mark the shape of the muscle at the previous time step, and the pink arrows mark the bending direction of the excited part. **e**, Modeling of the activation mechanism of microbubble-array artificial muscles. The pink, yellow, and blue boxes represent differently sized microbubble-array segments. The upper portion illustrates schematics of the cross-section of the artificial muscle, each corresponding to a specific second moment of area (*I*). *F_i_* denotes the thrust force generated by the microstreaming (here the yellow segment of the muscle generates thrust), *Δ* and *θ* denote the deflection and rotation angle along the long axis (*Y* axis). The lower left inset shows the modeling of a microbubble, where *R*_c_ is the radius of the cavity, *R* is the curvature radius of the trapped microbubble, and *a* is the amplitude of the center displacement during oscillation.

### Modeling of microbubble-array artificial muscles

We have developed a theoretical model to improve our understanding of the response of soft artificial muscles to sound waves. This model divides the entire artificial muscle into discrete segments that correspond to the patterned microbubble arrays, as illustrated in **Fig. 2e**. We began by modeling the acoustofluidic thrust force from a single trapped microbubble and analyzing the resulting artificial muscle deformation. To formulate the model, we assumed that (i) the ultrasound produces a homogeneous oscillating pressure field at the microbubble, generating the thrust force; (ii) the beam’s oscillation amplitude is negligible compared to that of the microbubbles, such that its motion does not significantly affect the surrounding flow field; (iii) hydrodynamic coupling between oscillating microbubbles can be neglected; (iv) the acoustic streaming force is dominated; (v) beam stretching is negligible; and (vi) gravity of the muscle does not influence the beam deformation.

To calculate the thrust force arising from acoustic streaming generated by a single oscillating microbubble, we adopted a model developed by Spelman and Lauga^49,50^. With additional approximations (**Supplementary Note 1**), we derived an expression for the thrust force

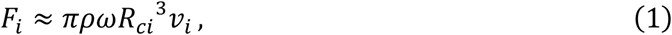

where *ρ* is the fluid density, *ω* the ultrasound frequency, *R*_c*i*_ the cavity radius in segment *i*, and *v_i_* the mean tangential velocity along the microbubble surface perpendicular to the beam, measured experimentally (**Extended Data Fig. 3**). For example, a microbubble with a 40 μm radius in water (*ρ* = 1000 kg/m³), excited at 27.6 kHz with 60 V_PP_, produced a measured velocity of *v_i_* = 2.5 mm/s, yielding a thrust force of *F_i_* = 13.9 nN according to Eq. (1). Scaling this to an array of approximately 18,000 uniformly sized microbubbles on a 30 mm × 5 mm artificial muscle yields a total force reaching up to 182 μN, corresponding to a force intensity of 1.21 μN/mm².

To describe the beam deformation, we parameterized the slender beam length by a variable *s*. Owing to planar symmetry, the deformation is fully described by the local slope angle *θ*(*s*). Using linear elasticity and the known orthogonal thrust force density, we derived the governing equation for *θ*(*s*) (see **Supplementary Note 2**). Assuming small variations in *θ* within each segment, we obtained an analytical expression for *θ*(*s*) in terms of the beam’s Young’s modulus *E*, second moment of area *I*, and the segmental thrust force densities. The resulting *Y*-direction deformation as a function of *s* in *i*-th segment is then given by

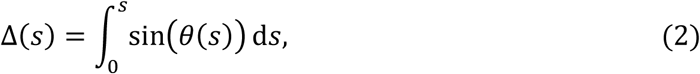

Our model is applicable to artificial muscles featuring both uniform-size and variable-size microbubble arrays. The model demonstrates that the deformation amplitude of an artificial muscle can be amplified quadratically by increasing the ultrasound excitation voltage (Δ ∝ V^2^), as shown in **Supplementary Fig. 4**. Besides, the model shows that the deformation can be increased by increasing the number of microbubbles (**Supplementary Fig. 5**). Additionally, larger deformation can be achieved by either reducing the material’s Young’s modulus or decreasing the muscle’s thickness (**Supplementary Fig. 6**). Furthermore, we envision that expanding the range of microbubble sizes enhances the manipulation freedom.

### Applications of microbubble-array artificial muscles

The development of programmable microbubble-array artificial muscles presents an exciting alternative for wireless actuation, enabling innovative designs in the field of soft robotics. Trapping and manipulating small, fragile model animals (e.g., zebrafish embryos) could become an appealing area of research in soft robotics. Conventional micro-tweezers often lack sufficient gripping force, while bulkier grippers risk damaging delicate targets. To address this, we designed a miniaturized soft gripper comprised of six to ten uniform-size microbubble array artificial muscles (**Supplementary Video 5**). Each tentacle houses over 60,000 microbubbles when submerged in water. As illustrated in **Fig. 3a**, upon subjected to an ultrasound stimulus (95.5 kHz, 60 V_PP_), the tentacles gripped a zebrafish larva within 100 milliseconds. When the ultrasound stimulus was deactivated, the larva easily swam away (**Supplementary Video 6**). Repeated actuation showed no notable heating or adverse effects on the larva, confirming the biocompatibility of the mechanism.

**Fig. 3.**
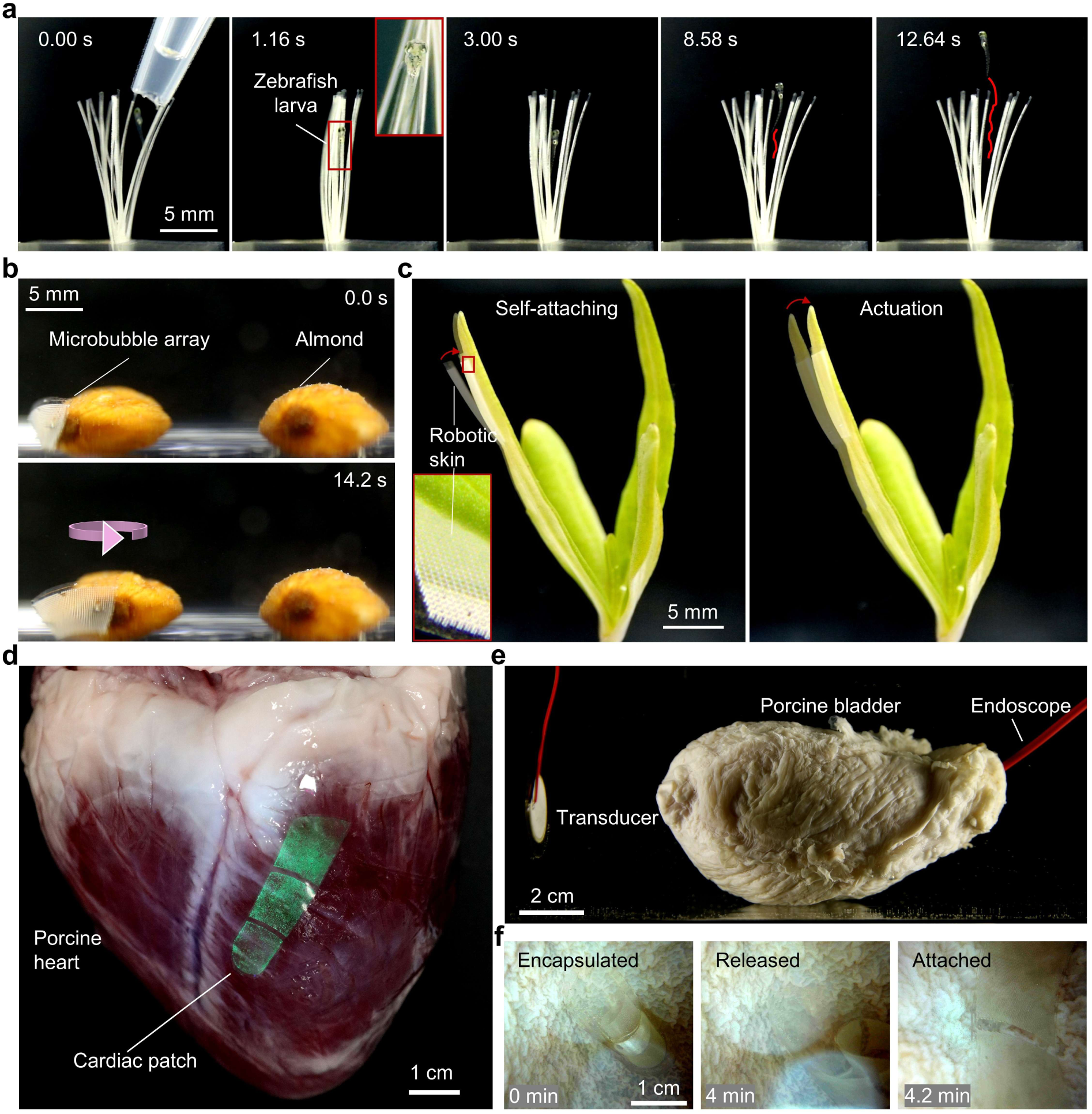
Adaptive gripper and robotic skin of microbubble-array artificial muscles. **a**, A live zebrafish larva trapped by a soft gripper featuring multiple microbubble-array artificial muscles (12 μm × 50 μm) as tentacles (10 mm × 0.7 mm × 80 μm). The inset shows a zoom-in view of the larva. **b**, Rotation of an almond by a conformable microbubble-array robotic skin (12 μm × 50 μm), and **c**, Closing of a piece of grass. The inset shows a zoom-in view of the microbubble array. **d**, Conformal attachment of a green fluorescently labelled cardiac patch (30 mm × 10 mm × 80 μm) to the epicardial surface of an ex vivo porcine heart. **e**, Setup showing an ex vivo porcine bladder with an ultrasound transducer positioned ∼5 cm from its left side and an endoscope inserted for internal visualization. **f**, Timelapse endoscopic images showing the encapsulated artificial muscle inside the bladder, its release at ∼3-5 min, and conformal attachment to the inner wall at 4.2 min under ultrasound activation.

We further demonstrated the artificial muscle as a conformable robotic skin capable of adhering to arbitrary surfaces and imparting motion to stationary objects. For example, we attached the robotic skin containing a uniform-size microbubble array to an arbitrary-shaped almond that exhibited controllable counter-clockwise rotation upon excitation at 95.5 kHz and 60 V_PP_ (**Fig. 3b** and **Supplementary Video 7**). We further show that upon switching on the ultrasound excitation, the robotic skin self-adhered to a blade of grass and enabled it to bend (**Fig. 3c** and **Supplementary Video 8**). The microbubble-array robotic skin offers the inanimate object diverse mobilities without notable size or mass increase.

Similarly, we demonstrated conformal attachment of the robotic skin—an artificial muscle containing a uniform-size microbubble array, to the epicardial surface of an ex vivo porcine heart, where it maintained functional adhesion for over 60 minutes at 96 kHz and 60 V_PP_ (**Fig. 3d**, **Extended Data Fig. 4** and **Supplementary Video 9**). By engineering different microbubble array geometries (e.g., square or circular) and tuning excitation frequency, we can generate multimodal shape transformations (**Extended Data Fig. 5** and **Supplementary Video 10**), selective and programmable localized mechanical forces and drug delivery (**Extended Data Fig. 6**). Localized stimulation enables on-demand mechanical actuation of soft biological tissues and could support a range of future cardiac therapies and clinically relevant interventions, such as targeted anti-fibrotic drug delivery and localized gene or mRNA therapy. These results highlight the potential for the future development of in vivo wireless and wearable devices.

To evaluate the potential for wireless robotic drug delivery and in situ deployment, the artificial muscle was pre-encapsulated in a biodegradable capsule designed for swallowable or minimally invasive delivery (**Fig. 3e**). Upon injection into an excised porcine bladder, the capsule gradually dissolved over ∼3–5 minutes, exposing the actuator to the surrounding environment. Following dissolution, ultrasound (96 kHz, 60 V_PP_) was applied to induce deformation of the actuator, allowing it to attach to the inner surface of the bladder (**Fig. 3f** and **Supplementary Video 11**).

Capitalizing on the dynamic deformation and rapid response capabilities of our artificial muscle, we engineered a bioinspired ultrasound-powered wireless stingraybot. The biomimetic stingraybot features two artificial muscles—designed to mimic the pectoral fins of a natural stingray— integrated on its sides. These pectoral fins incorporate arrays of differently sized microbubbles (12 μm, 16 μm, and 66 μm in diameter; 50 μm in depth) patterned along the head-to-tail axis and paired with a PDMS block for buoyancy adjustment. When exposed to a sweeping-frequency ultrasound stimulation (30 to 90 kHz over 2 s at 60 V_PP_), the stingraybot’s fins exhibit an undulatory motion that mimics the natural motion of a stingray (**Fig. 4a**, **Supplementary Video 12**). Upon release, the stingraybot propels forward at an initial speed of ∼0.8 body lengths per second (**Fig. 4b)**. More control experiments on stingraybot propulsion are provided in the **Methods** section.

**Fig. 4.**
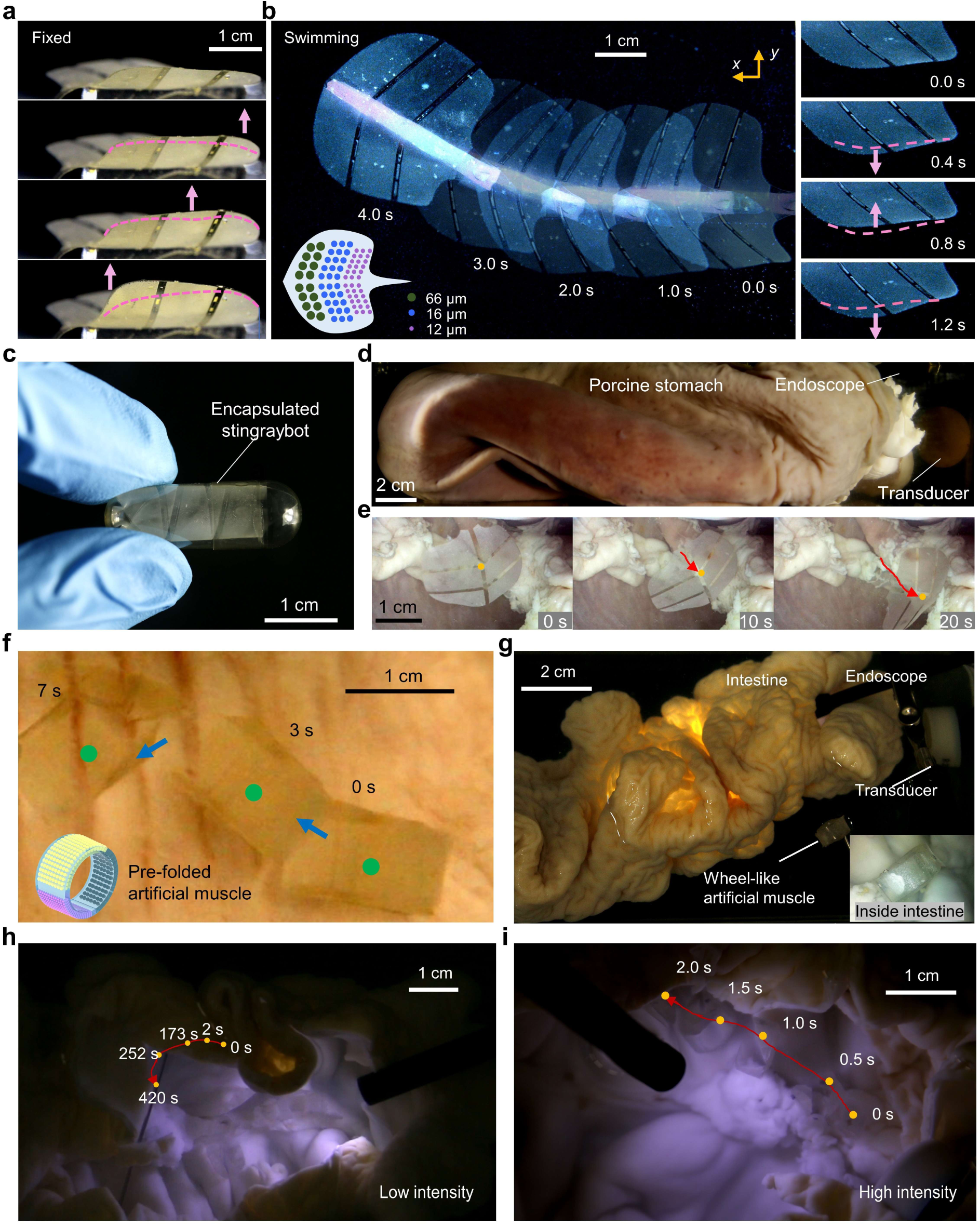
Bioinspired swimming and navigation within ex vivo biomedical environment powered by microbubble-array artificial muscles. a, Undulation motion of the microbubble-array (12 μm, 16 μm, and 66 μm in diameter; 50 μm in depth) fins of the bioinspired stingraybot before releasing. b, Forward swimming of the stringraybot under sweeping-frequency ultrasound excitation (30-90 kHz, 2 s and 60 V_PP_). The right panel displays the undulatory fin in motion during swimming, while the lower inset presents a schematic of the microbubble arrays patterned onto the fin. In **a** and **b**, the **pink** dashed line and arrow respectively denote the shape of the fins in the last time step and the moving direction of the fins in the current time step. **c**, Photograph of the edible capsule (hydroxypropyl methylcellulose/corn starch, in 27 mm length and 12 mm diameter) containing the pre-folded stingraybot. **d**, Setup for the release and navigation of the encapsulated artificial muscle in an excised porcine stomach, with an ultrasound transducer positioned ∼3 cm from the right side of the stomach and an endoscope inserted into the lumen to visualize the motion of the artificial muscle. **e**, Time-lapse images of stingraybot locomotion inside an excised porcine stomach. **f**, Locomotion of a pre-folded, wheel-shaped artificial muscle (30 mm × 5 mm × 80 μm) composed of variable-size microbubble arrays (12 μm, 16 μm, and 66 μm in diameter; 50 μm in depth) inside a porcine stomach. The artificial muscle propels along the stomach surface under sweeping-frequency ultrasound excitation (30-100 kHz, 2 s and 60 V_PP_), with its trajectory indicated by blue arrows. Green dots indicate the center of the artificial muscle. Inset shows the pre-folded configuration. **g,** Setup for ex vivo manipulation of a prefolded artificial muscle inside an excised porcine large intestine, with ultrasound transducers placed externally and an endoscope inserted into the lumen. Inset, endoscopic view of the artificial muscle. **h**, Timelapse images showing the artificial muscle rolling along the curved mucosal wall under ultrasound sweeping-frequency (30–100 kHz over 2 s, 60 V_PP_) delivered by a disk piezo transducer. **i**, Locomotion of the artificial muscle driven by a HIFU transducer (1–3 MHz over 1 s, 60 V_PP_). Red lines trace the trajectory, and yellow dots denote the centre position over time.

To implement practical biomedical applications, we demonstrated ultrasound-guided navigation of a pre-folded artificial muscle through ex vivo porcine gastrointestinal tissues, targeting use cases such as site-specific drug release for gastrointestinal disorders, minimally invasive access to inflamed or fibrotic tissue, and wireless actuation in regions inaccessible to rigid tools. We first pre-folded and encapsulated a stingraybot within an edible capsule (**Fig. 4c**). Once released into the stomach (**Fig. 4d**), the stingraybot propelled on demand within the confined biomedical environment under ultrasound actuation (**Fig. 4e, Supplementary Video 13**). In a separate experiment, we pre-folded a linear artificial muscle—with variable-size microbubble arrays arranged along its outer surface—into a cylindrical, wheel-like structure. Under sweeping-frequency, the actuator exhibited directional rolling propulsion along the complex mucosal surfaces of the stomach and intestine (**Fig. 4f–i, Supplementary Video 14**), illustrating its potential for soft robotic intervention and targeted delivery within the gastrointestinal tract. Future work will focus on parametric studies, dynamic folding strategies, and steering-enabled configurations of the artificial muscle across varied tissue geometries and fluidic environments.

## Discussion

We have introduced a new class of soft artificial muscles that utilize acoustically activated microbubble arrays to achieve programmable actuation. These artificial muscles exhibit dynamic programmability, high force intensity (∼1.21 μN/mm^2^), rapid responsiveness (sub-100 ms), and wireless controllability, all while maintaining exceptionally compact (3,000 microbubbles/mm²) and lightweight (0.047 mg/mm^2^). Through the strategic use of microbubble configurations and voltage and frequency as ultrasound excitation parameters, we engineered a diverse range of preprogrammed movements (e.g., undulatory motion) and demonstrated their applicability across artificial muscle platforms. We showcased the strength and durability of these muscles by integrating variable-size microbubble arrays into functional devices such as a soft gripper, a robotic skin, and a biomimetic stingray robot. We also established a theoretical model that elucidates the actuation mechanism, which serves as a guide for the design of microbubble array patterns with enhanced actuation performance. These artificial muscles offer extensive applications in robotics, flexible electronics, wearable technologies, prosthetics, biomedical instrumentation, and beyond.

To optimize the artificial muscle performance through the geometric design and the density of the microcavities, preliminary experiments revealed that converging trapezoidal cavities generate **∼**3**×** stronger streaming velocities than diverging shapes (**Supplementary Fig. 7**) and a higher density of cavities causes a larger deformation. By incorporating geometric and density optimization with systematic characterization, one can establish a predictive design framework for actuators with tailored deformation profiles—enabling precision control in applications from soft robotics to biomedical devices. Future studies could also explore the application of confocal sound sources, such as high-intensity focused ultrasound (**Supplementary Fig. 8**) to achieve local millimeter deformation—which potentially could lead to new tools for applications such as in vivo mechanotransduction and spatially targeted drug delivery. Additionally, the bubble-based mechanism is material-agnostic of our system and can be extended to biocompatible or biodegradable matrices, such as hydrogels and biodegradable polymers, for more biomedical applications. More robustness evaluation on our ansatz across fluid media are provided in the **Methods** section.

Despite promising results, certain limitations remain. Prolonged actuation triggers microbubble growth within the cavities, destabilizing the muscle operation after ∼30 minutes (**Supplementary Fig. 9**). Resubmersion in water restores the function, and sealing cavities with a thin PDMS membrane will offer a long-term robust solution (**Supplementary Fig. 10)**. Additionally, the stingraybot’s distance-dependent actuation must be taken into account for untethered operation. Our preliminary experiments at varying transducer distances revealed deformation decays with increasing distance (**Supplementary Fig. 11**), dropping by ∼50% at 5 cm compared to the deformation at 1 cm. While this limitation is less critical in vivo (where the robot is intended to operate in confined volumes, e.g., the bladder), optimizing the ultrasound source configurations and the actuation voltage to compensate for ultrasound intensity decay over distance can enhance the performance.

Looking ahead, these artificial muscles hold transformative potential across cutting-edge fields such as soft robotics, haptic medical devices, and minimally invasive surgery. Future research should focus on refining the scalability of these systems across multiple scales (**Extended Data Fig. 7**), enhancing their force generation capabilities, and integrating them into complex devices for biomedical applications.

## Methods

### 1. Fabrication of artificial muscles

The negative patterns of artificial muscles are first designed in the commercial electronic design automation software (as shown in **Supplementary Fig. 1**). The patterns are transferred into a photomask by a direct writing laser (DWL2000) machine in a cleaning room (BRNC, Ruschlikon, Switzerland). Then, we spin coat the negative photoresist SU8-3025 on a 4-inch silicon wafer. By the standard lithographic fabrication, the patterns are transferred to the photoresist via exposing to UV light through the mask. After the developing process, the negative patterns of the microbubble arrays, i.e., micropillars, are additive on the wafer. The height of the micropillar depends on the spinning speed. Next, to enhance surface properties, a silane-based hydrophobic treatment was applied to the 4-inch wafer with micropillars for 1 hour (see fabrication flow at **Supplementary Fig. 12**). The PDMS (polydimethylsiloxane) used in this process was prepared with a 10:1 ratio of base to curing agent. Then, the PDMS mixture was poured onto the wafer. To ensure a high-quality coating, the mixture was degassed under vacuum pressure of less than 1 mBar. After degassing, spin-coating was performed on the wafer. Different spin speeds result in varying PDMS membrane thicknesses (**Supplementary Fig. 2**). After spin-coating, the PDMS was vacuumed again and cured in a sequential heating process: 1 hour at 60°C, followed by 1 hour at 80°C, and finally 1 hour at 100°C. Finally, the PDMS soft membrane was cured and ripped off from the wafer. This process yielded a uniform PDMS layer suitable for use in artificial muscle and soft robotics applications. In all our experiments, each cavity consistently trapped only a single bubble as the artificial muscle submerged into the water (**Supplementary Fig. 13**).

### 2. Acoustic setup

For the microscale characterization of microbubbles, the experimental setup is built on a thin glass substrate with dimensions of 24 mm × 60 mm × 0.18 mm. As shown in **Supplementary Fig. 14**, a circular piezoelectric transducer (27 mm × 0.54 mm, resonance frequency 4.6 kHz ± 4%, Murata 7BB-27-4L0) is affixed to the glass substrate using an epoxy resin (2-K-Epoxidkleber, UHU Schnellfest). A square PDMS acoustic chamber (10 mm × 10 mm × 5 mm) is positioned in the front of the transducer, which is filled with deionized water and covered with a cover glass (22 mm × 22 mm × 0.18 mm). An artificial muscle was suspended in the center of the chamber with one end clamped to the sidewall and the other end left free. The substrate is then mounted on an inverted microscope (Axiovert 200M, ZEISS).

For the macroscale actuation of artificial muscles by sound, the experimental setup is built on a plastic tank with dimensions of 10 cm × 10 cm × 8 cm and a thickness of 2 mm (Specially, for ex vivo porcine experiments, the chamber size is 30 cm × 15 cm × 15 cm and a thickness of 2 mm). As shown in **Supplementary Fig. 15**, the circular piezoelectric transducers are affixed to the inside surfaces and the bottom surface of the tank using the epoxy resin or directly submerged into the liquid because the deionized water filled in the tank is not conductive. An artificial muscle was suspended inside the chamber with one end clamped and three cameras are placed around the tank to capture the actuation of acoustic artificial muscles from multiple view angles. Additionally, a miniaturized endoscopic camera is utilized (8 mm diameter and 1080P resolution, FuanTech) to capture the inside view of porcine specimens. An electronic function generator (AFG-3011C, Tektronix) and an amplifier (0-60 V_PP_, 15x amplification, High Wave 3.2, Digitum-Elektronik) are connected to the transducer to generate sound waves with tunable excitation frequencies and voltages. Square waves effectively drive the artificial muscle, achieving maximum deformation and outperforming other tested waveforms, such as sinusoidal and triangular waveforms under equivalent excitation conditions (**Supplementary Fig. 16**).

### 3. Microstreaming characterization

We evaluated the microstreaming jets generated by ultrasound-driven microbubbles embedded in the muscle, using 6 μm tracer particles in water and particle image velocimetry (PIV) analysis. Three uniform-size microbubble arrays, each comprising a 4×4 grid of microbubbles with diameters of 40 μm, 60 μm, and 80 μm (150 μm in depth), were individually selected and tested in separate miniaturized artificial muscles (500 μm × 500 μm × 200 μm) (**Supplementary Fig. 17a** and **Supplementary Video 15**). When activated at their respective resonance frequencies 76.3 kHz, 57.4 kHz, and 27.6 kHz, we measured the microstreaming velocity 80 μm away from the bubble interface and observed a quadratic relationship between the average velocity and excitation voltage (**Supplementary Fig. 17b**). The streaming velocity near the bubble reached 2.5 mm/s at 60 V_PP_. This voltage-dependent microstreaming directly correlates with the reverse thrust generated by the microbubble array, demonstrating that thrust magnitude can be dynamically tuned by adjusting the ultrasound excitation.

We further investigated the selective actuation of a variable-size microbubble array of 40 μm, 60 μm, and 80 μm diameter, each 150 μm in depth, integrated within a single miniaturized artificial muscle (500 μm × 500 μm × 200 μm) with corresponding frequencies (27.6 kHz, 57.4 kHz, and 76.3 kHz, respectively). The PIV analysis reveals that the microstreaming developed by the 80 μm bubbles generated an average velocity of 0.23 mm/s at 27.6 kHz, which was markedly stronger compared to the velocities (< 0.05 mm/s) produced by the other two microbubble arrays at the same voltage (15 V_PP_). Similarly, adjusting the frequency to 57.4 kHz (76.3 kHz) selectively activates the 60 μm (40 μm) bubble array, resulting in more intense streaming at 0.174 mm/s (0.075 mm/s), in contrast to other arrays (**Extended Data Fig. 2**). Applying a sweeping frequency (10– 90 kHz) over 4 seconds at 30 V_PP_ enabled wave propagation across the artificial muscle (**Supplementary Video 16**).

### 4. Control experiments on artificial muscle deformation

To determine the key factors influencing muscle deformation, a set of control experiments was performed. We first examined the streaming jets of a uniform-size microbubble array artificial muscle (1 cm × 0.3 cm × 80 μm) patterned with over 800 microcavities (each 40 μm in diameter and 50 μm in depth). **Supplementary Video 17** shows that an artificial muscle without microbubbles exhibited minor deformation, with no noticeable microstreaming observable across excitation frequencies sweeps from 1 to 100 kHz at 60 V_PP_. When microbubbles were introduced into the actuator, they generated pronounced microstreaming near the bubbles (∼0.8 mm/s) upon increased ultrasound stimulation frequency to 9.5 kHz, resulting in substantial deformation of the artificial muscle.

### 5. Repeatability and characterization of artificial muscle deformation

We then assessed the repeatability of the artificial muscle’s deformation under identical excitation conditions. With the transducer close to the microbubble-embedded side, as shown in the left panel of **Extended Data Fig. 8a**. When stimulated with ultrasound pulses (80.5 kHz, 60 V_PP_, and 1 s on/off cycle), the muscle exhibited repeatable bending within 150 cycles, with an error of ±0.8 mm, representing 2.7% of the total beam length (**Extended Data Fig. 8b**). With more excitation cycles (500 cycles) of the artificial muscle, the deformation exhibited larger error (∼10%). After 10,000 cycles, there were no observable microbubbles in the artificial muscle, and the artificial muscle exhibited minor deformation. Furthermore, **Extended Data Fig. 8c** reveals a quadratic relationship between the applied voltage and the mean deformation amplitude of artificial muscles patterned with microbubbles of various diameters (40 μm, 60 μm, and 80 μm) when driven at their respective resonance frequencies (80.5 kHz, 62.5 kHz, and 30.3 kHz). Additionally, the PDMS beam, in the absence of microbubbles, exhibited limited bending (∼7% of the 40 μm microbubble-array artificial muscle’s deformation at 60 V_PP_) caused by the weak radiation force from incident sound waves originating from the transducer.

### 6. Control experiments on stingraybot propulsion

In control experiments, a stingraybot without microbubbles exhibited no undulatory motion along its fins under ultrasound excitation and sank without notable lateral displacement (**Supplementary Video 18**). Notably, under continuous single-frequency excitation (60 V_PP_ at 33.2 kHz, 85.2 kHz, and 96.2 kHz) targeting distinct microbubble arrays (with diameters of 66 μm, 16 μm, and 12 μm, respectively), the stingraybot exhibited only limited locomotion (<1 body length). In comparison, sweeping-frequency excitation (10–100 kHz over 2 s) elicited sustained undulatory motion, allowing the stingraybot to swim a significantly greater distance (>3.5 body lengths), as shown in **Supplementary Fig. 18**. These results suggest that the forward motion of the stingraybot is predominant by the propulsion force generated by the sequential undulatory motion, resulting from the reverse thrust generated by microbubble arrays. Moreover, enhancing the design of the stingraybot with additional microbubble sizes could expand its maneuverability. For instance, integrating a navigation tail with microbubble arrays of different sizes on either side enables directional control. When activated at their respective resonance frequencies on one side, these arrays generate an asymmetric torque (**Supplementary Fig. 19**), enabling steer the stingraybot via tail rotation. Since the stingraybot is stealthy and transparent, we further envision that our stingraybot could be employed for environmental data collection or behavioral research on real organisms, for example, detecting water quality within coral reefs and recording swarm interaction by blending into schools of fish.

### 7. Robustness evaluation

To evaluate the robustness of our ansatz across fluid media, we quantified artificial muscle deformation in 100% porcine blood, observing amplitudes of ∼0.4, 1.0, 2.7, and 4.4 mm at 15, 30, 45, and 60 V_PP_, respectively under 96 kHz ultrasound excitation (**Extended Data Fig. 9**). As complementary evidence, we studied the artificial muscle performance in various aqueous solutions (deionized water, tap water, and 25–100% glycerol solutions) as shown in **Supplementary Fig. 20**. The deformation exhibited an inverse relationship with glycerol concentration, with the largest deformation of ∼11.3 mm in a 25% glycerol solution, followed by ∼8.4 mm in 50% glycerol and 3.7 mm in 75% glycerol. The deformation was almost negligible in 100% glycerol. These results clearly demonstrate that the actuator functions effectively in full blood, validating its potential for in vivo applications in fluids with physiological viscosity. We next evaluated artificial muscle actuation in the presence of solid obstructions (**Supplementary Fig. 21**). A frontal obstruction (partially blocking ultrasound) reduced the deformation by 80–90% (0.5–1 mm tip deformation versus 4.8 mm unobstructed). A lateral placement caused moderate attenuation (∼2.5 mm), while posterior positioning retained a better performance (3.8 mm). Furthermore, experimental results demonstrated significant deformation of the artificial muscle behind excised porcine ribs (**Supplementary Fig. 22**). Thus, actuators remained functional near obstacles but required strategic positioning to maximize deformation. Our preliminary results also revealed negligible heating effects near the piezoelectric transducer during artificial muscle and stingraybot operation (**Supplementary Fig. 23**), underscoring the thermally benign nature of our acoustic platform. While frequency-dependent selectivity was achieved, some cross-excitation between microbubble arrays was observed. This effect was mitigated under sweeping-frequency actuation, and temporal control over the sweep dynamics plays a key role in preserving spatial selectivity and ensuring reliable, programmable motion. In vivo biomedical environments present additional challenges such as complex fluid flow, irregular geometry, and variable temperature gradients, all of which may distort ultrasound propagation. Although the actuator demonstrated robust and competitive performance under static conditions with other methodologies (**Extended Data Fig. 10** and **Supplementary Fig. 24)**, future work will explore flow-resilient designs, including optimized microbubble-array geometries, flexible ultrasound configurations, and real-time actuation control strategies to maintain reliable performance in dynamic fluid environments.

### 8. Numerical simulations

Finite element numerical simulations are conducted using the commercial COMSOL Multiphysics software (v6.1, Burlington, MA), including simulations on the acoustic pressure field in the small PDMS chamber, acoustic streaming generated by variable-size microbubbles in the small PDMS chamber, the acoustic pressure field in the big acoustic tank, and the deformations of the artificial muscle. All simulations are performed with dimensions and material properties consistent with the experiments. Physics modules of simulations on acoustic pressure include the solid mechanics, electrostatics, pressure acoustics fields, creeping flow, and heat transfer in solids and fluids. Simulations on the deformations of artificial muscles are performed using the solid mechanics module with corresponding boundary conditions and force conditions. The microstreaming-generated thrust is assumed to be a point force which is loaded on the bottom of each microcavity. Additionally, numerical calculations based on the theoretical model are performed using the commercial Matlab software (R2022b). See **Supplementary Notes** for simulation details.

### 9. Imaging and analysis

The microscale characterization of microbubbles is recorded with a high-speed camera (Chronos 1.4, Kron Technologies) attached to the inverted microscope. Recording frame rates range from 1069 to 32,668 frames per second (fps). The macroscale motion of ultrasound artificial muscles is recorded with a high-sensitive camera (Canon 6D and 24−70 mm camera lens, Canon). The recording frame rate is 50 fps. Recorded footages are analyzed in ImageJ.

#### Acknowledgments

D.A. was funded by the European Research Council (ERC) under the European Union’s Horizon 2020 research and innovation program (grant agreement No. 853309, SONOBOTS); the Swiss National Science Foundation (SNSF) through the SNSF Project funding MINT 2022 (grant agreement No. 213058), Spark Grant (grant agreement No. 221285); and the ETH Research Grant (grant agreement No. ETH-08 20-1). Z.S. acknowledges financial support from the China Scholarship Council (202106320193). Z.Z. acknowledges financial support from the China Scholarship Council (202006210065). N.N. acknowledges the funding from the United States National Science Foundation (OIA-2229636, CBET-2407937). R.W. is funded by the Deutsche Forschungsgemeinschaft (DFG, German Research Foundation) — 535275785. We thank the operational team of the clean room facility in the Binnig and Rohrer Nanotechnology Center (BRNC) for their helpful discussion on the fabrication wafer. The authors thank Dan Schoenenberger for his helpful contribution to the initial fabrication and testing.

## Author contributions

D.A. conceived and supervised the project. R.W. supervised J.S.. Z.Z., Z.S., and D.A. contributed to the designs of the actuator and soft robots. Z.S. and Z.Z. developed the fabrication of prototypes, performed experiments and data analysis. S.C.F.N. provided the zebrafish embryos and provided guidance on the zebrafish experiments. R.W., N.N., J.S., Z.S., Z.Z., and D.A. contributed to the theoretical understanding. J.S. provided the analytical derivations on the acoustofluidic thrust force and the beam deformation. Z.S. contributed to the numerical simulations. Z.Z., Z.S., R.W., J.S., N.N., and D.A. contributed to the scientific presentation, discussion, manuscript revision, and data analysis.

## Competing interests

The authors declare that they have no competing interests.

## Data and materials availability

All data needed to evaluate the conclusions in the paper are present in the paper and/or the Supplementary Materials.

## Extended Data

**Extended Data Fig. 1.**
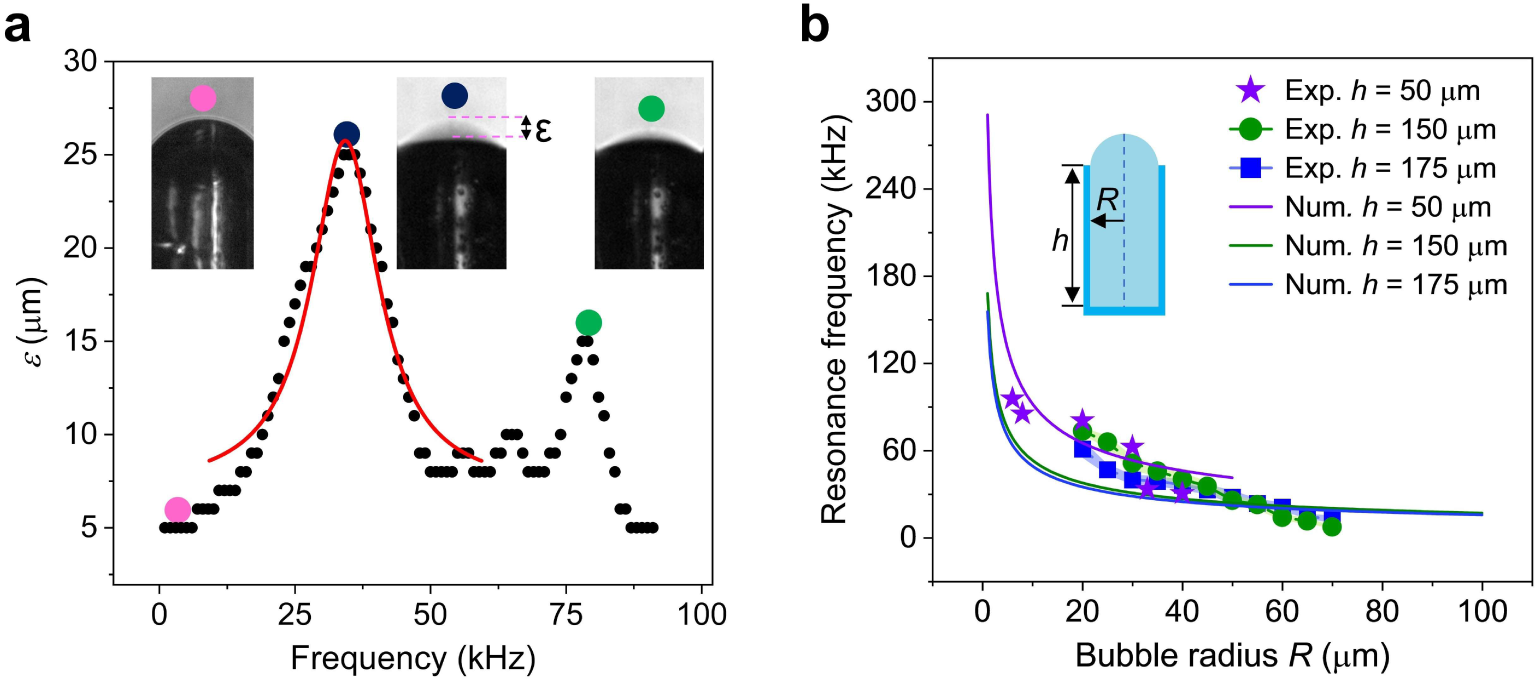
Characterization of the resonance frequency of different sized microbubbles. **a**, Oscillation amplitude of a 60 μm × 150 μm microbubble as a function of the ultrasound excitation frequency. According to the measured oscillation amplitude of the microbubbles, the resonance frequency is identified to be 35.5 kHz. **b**, Experimentally measured resonance frequencies of microbubbles with depths of 150 μm and 175 μm, radii ranging from 20 to 70 μm in 5 μm increments, and a depth of 50 μm with radii of 6 μm, 8 μm, 20 μm, 30 μm, 33 μm and 40 μm. The solid lines represent numerically predicted results. Additionally, notable discrepancies exist between the calculated and measured resonance frequencies, which may originate from transducer coupling with the glass slide, interactions between bubbles, or other factors.

**Extended Data Fig. 2.**
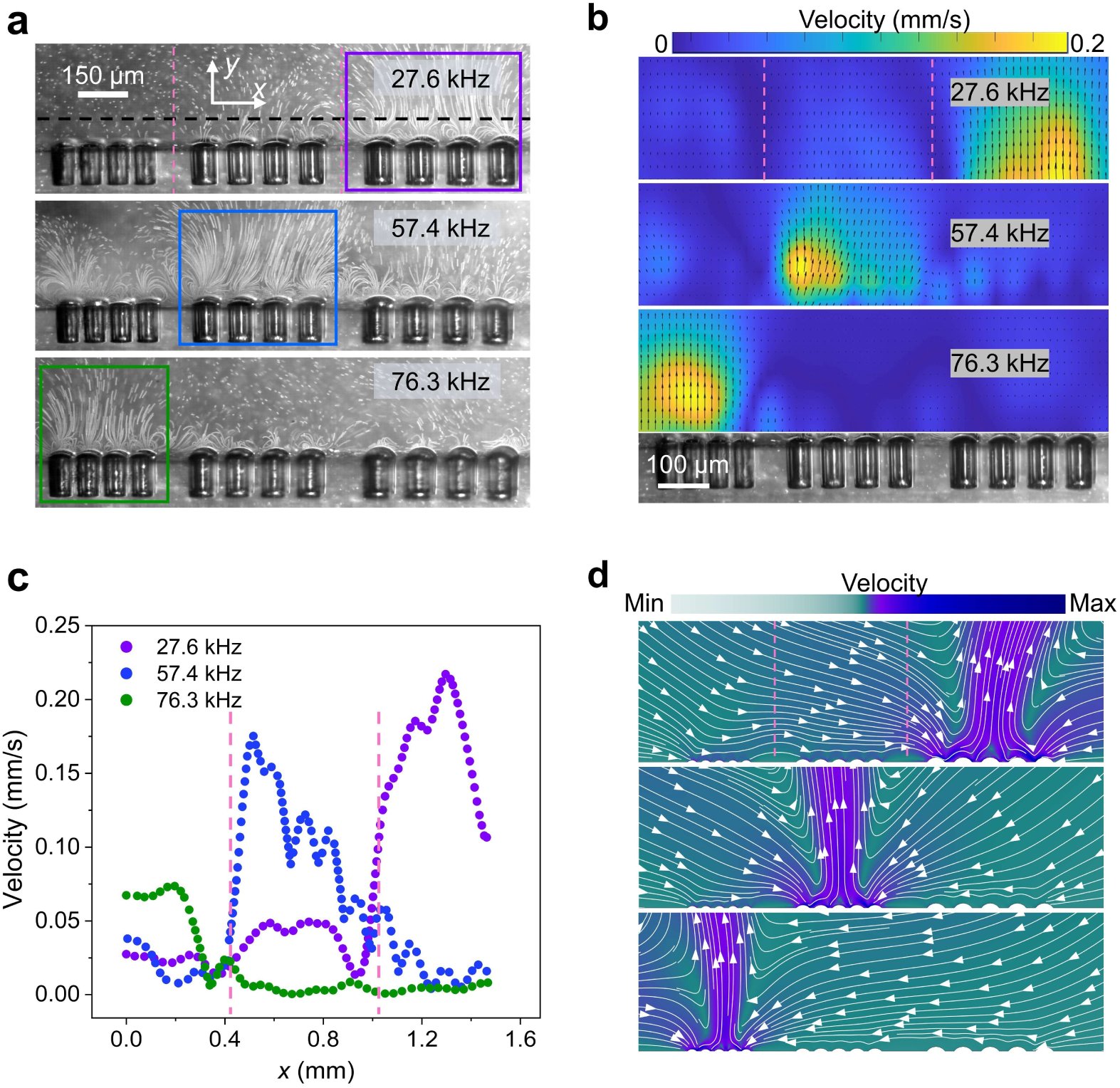
Selective actuation of microbubble arrays. **a**, Microstreaming respectively generated by (40 μm, 60 μm, and 80 μm) microbubble arrays under ultrasound frequencies of 76.3 kHz (green), 57.4 kHz (blue), and 27.6 kHz (purple). Colored boxes indicate the activated microbubbles. The black dashed line denotes the measurement position of the microstreaming velocity, which is 100 μm away from the surface. **b**, Measured streaming flow of **a** with 2 μm trace microparticles. The streaming flow field was analyzed by PIV (Matlab R2022b, PIVlab 2.60). The color bar denotes the particle moving velocity perpendicular to and extending away from the bubble surface. **c**, Plot of the measured microstreaming velocity along the long axis of the variable-size microbubble array under different excitation frequencies. **d**, Simulations of microstreaming around the bubble array with three different excitation frequencies. The pink dotted lines in **a**, **b**, **c**, and **d** delineate the boundaries of three distinct microbubble arrays.

**Extended Data Fig. 3.**
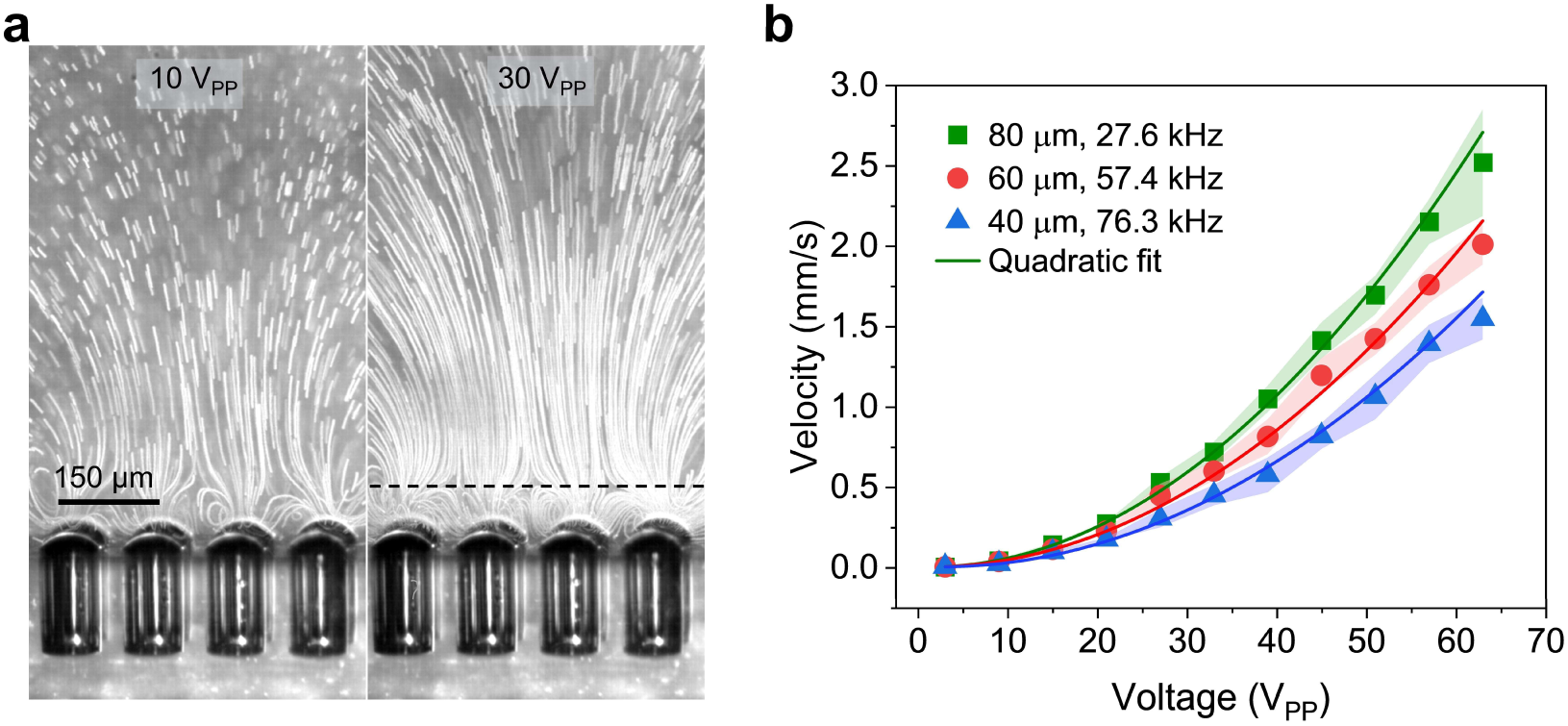
Measurement of microstreaming velocity by PIV. **a**, Microstreaming generated by a 4×4 80 μm × 150 μm microbubble array under an excitation frequency of 27.6 kHz and two different voltages of 10 V_PP_ (left panel) and 30 V_PP_ (right panel). The black dashed line denotes the measurement position of the microstreaming velocity, which is 80 μm away from the surface. **b**, Plot of microstreaming velocity versus ultrasound excitation voltage respectively measured by 4×4 microbubble arrays with three different sizes (40 μm, 60 μm, and 80 μm). The solid lines are the quadratic fitting results.

**Extended Data Fig. 4.**
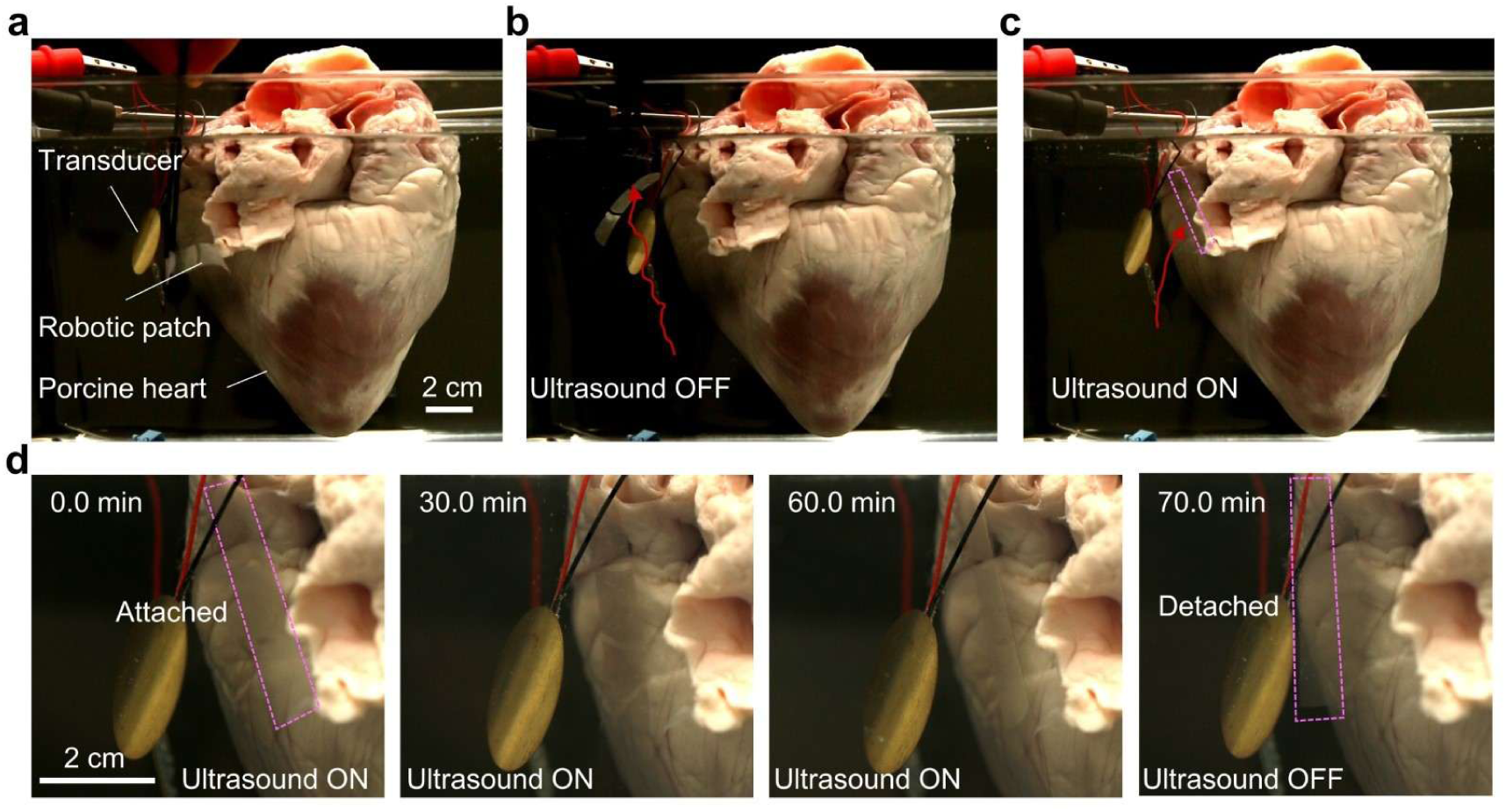
Attachment of a robotic patch to an ex vivo porcine heart. **a**, Experimental setup showing an artificial muscle positioned between an ultrasound transducer (operated at 96 kHz and 60 V_PP_) and the heart, separated by ∼2.5 cm. **b**, The muscle is released from tweezers positioned at the bottom and rises upward due to buoyancy (trajectory indicated by the red arrow). **c**, The artificial muscle conforms to the surface of the heart when stimulated by ultrasound (trajectory indicated by the red arrow). **d**, Time course of attachment robustness, demonstrating firm conformation and adhesion of the artificial muscle to the heart from 0 to 60 minutes under ultrasound actuation, followed by detachment upon ultrasound deactivation at 70 minutes. The pink dashed rectangle indicates the location of the artificial muscle.

**Extended Data Fig. 5.**
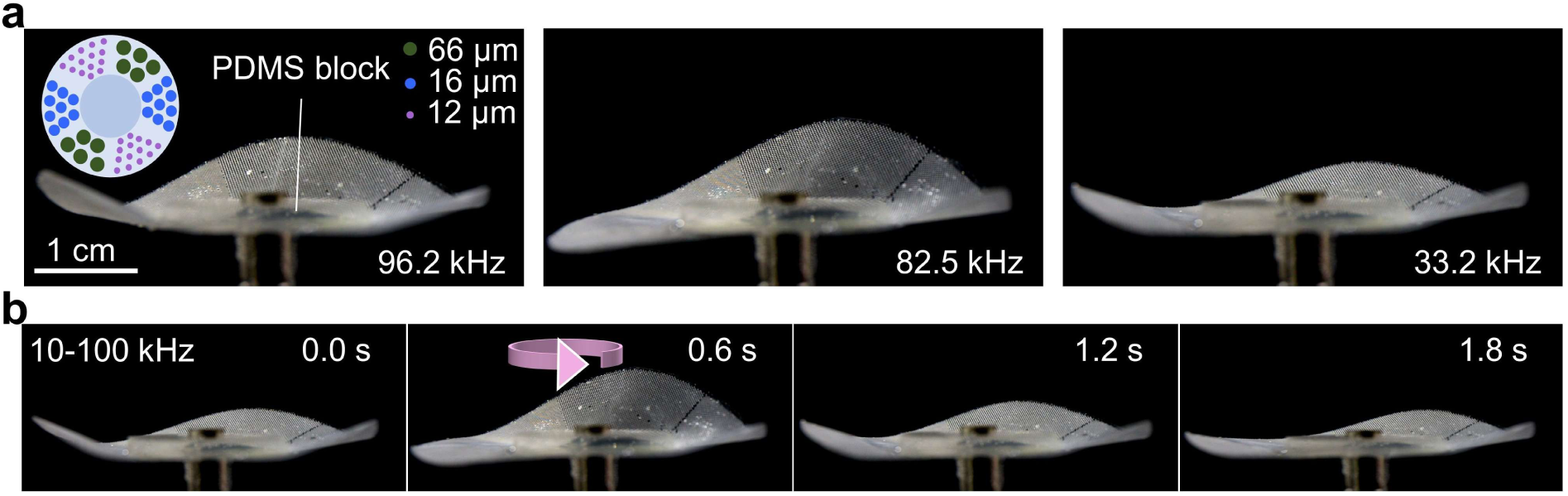
Multimodal shape transformation of a microbubble array-patterned functional surface. **b**, Multimodal shape transformation of a circular surface under continuous ultrasound excitation frequencies of 96.2 kHz, 82.5 kHz, and 33.2 kHz, respectively. The circular surface was topped with a circular PDMS block (2 cm diameter, 0.3 cm thickness) to reduce buoyancy. The inset shows a schematic of the microbubble array patterned on the surface. **c**, Dynamic shape transformation of the circular surface under sweeping-frequency ultrasound excitation spanning from 10 kHz to 100 kHz over 2 s.

**Extended Data Fig. 6.**
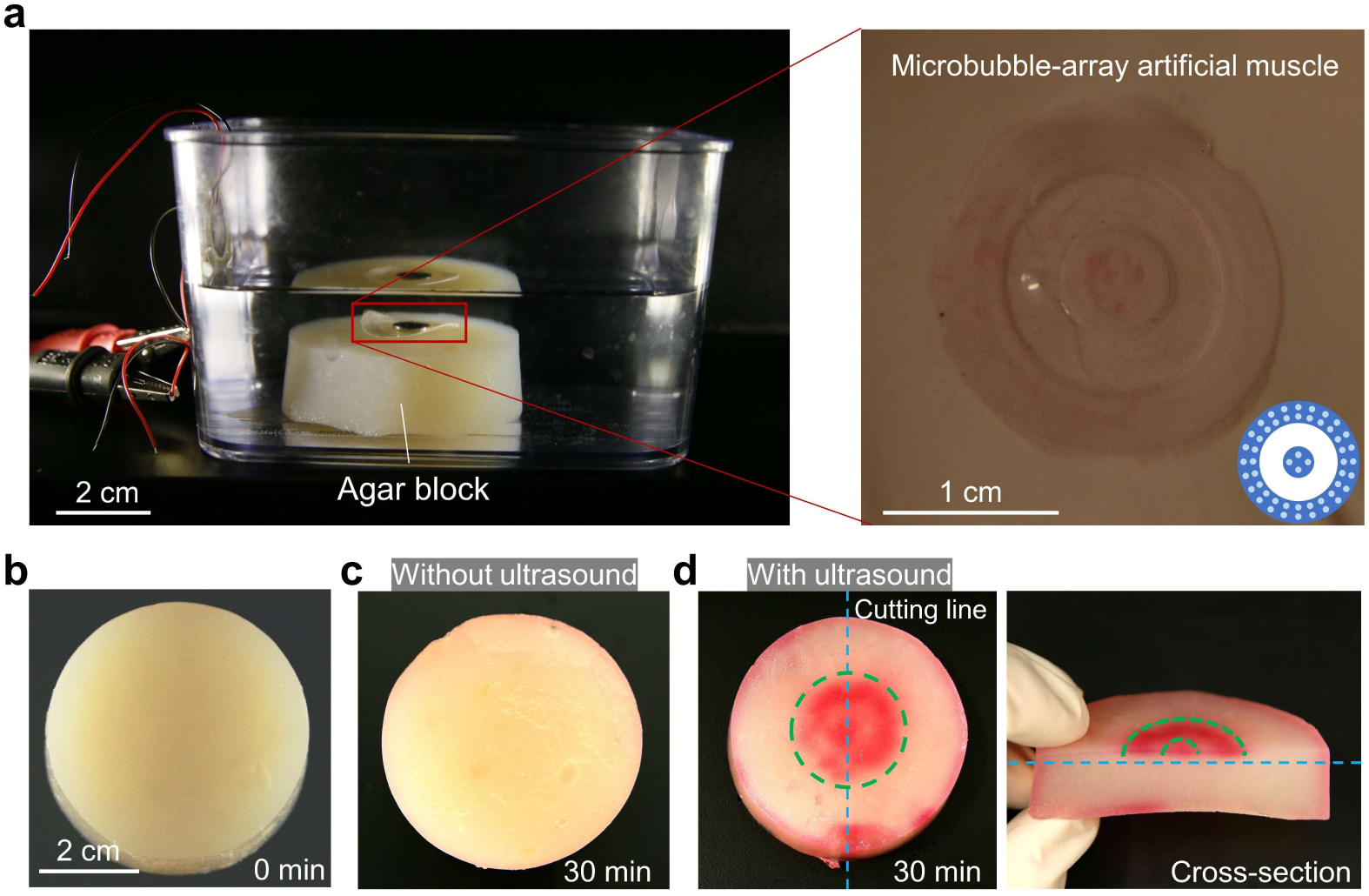
Ultrasound -enhanced dye delivery into an agar phantom by a microbubble -array robotic patch. **a**, Setup illustrates an agar block resting in an acoustic tank, with a piezo transducer positioned 5 cm to the left. A circular robotic patch is placed on the agar block with its microbubble arrays facing downward. The zoom--in highlights the patch surface; inset shows the patterned microbubble array. **b**, Agar block prior to dye exposure. **c**, Control condition showing the agar block after 30 min in a dye-filled tank without ultrasound actuation. **d,** Top and cross-sectional views of the agar block after 30 min of ultrasound actuation (96 kHz, 60 V_PP_), revealing enhanced dye penetration. The blue dashed line indicates the cutting plane; green dashed lines delineate the boundaries of the penetrated region.

**Extended Data Fig. 7.**
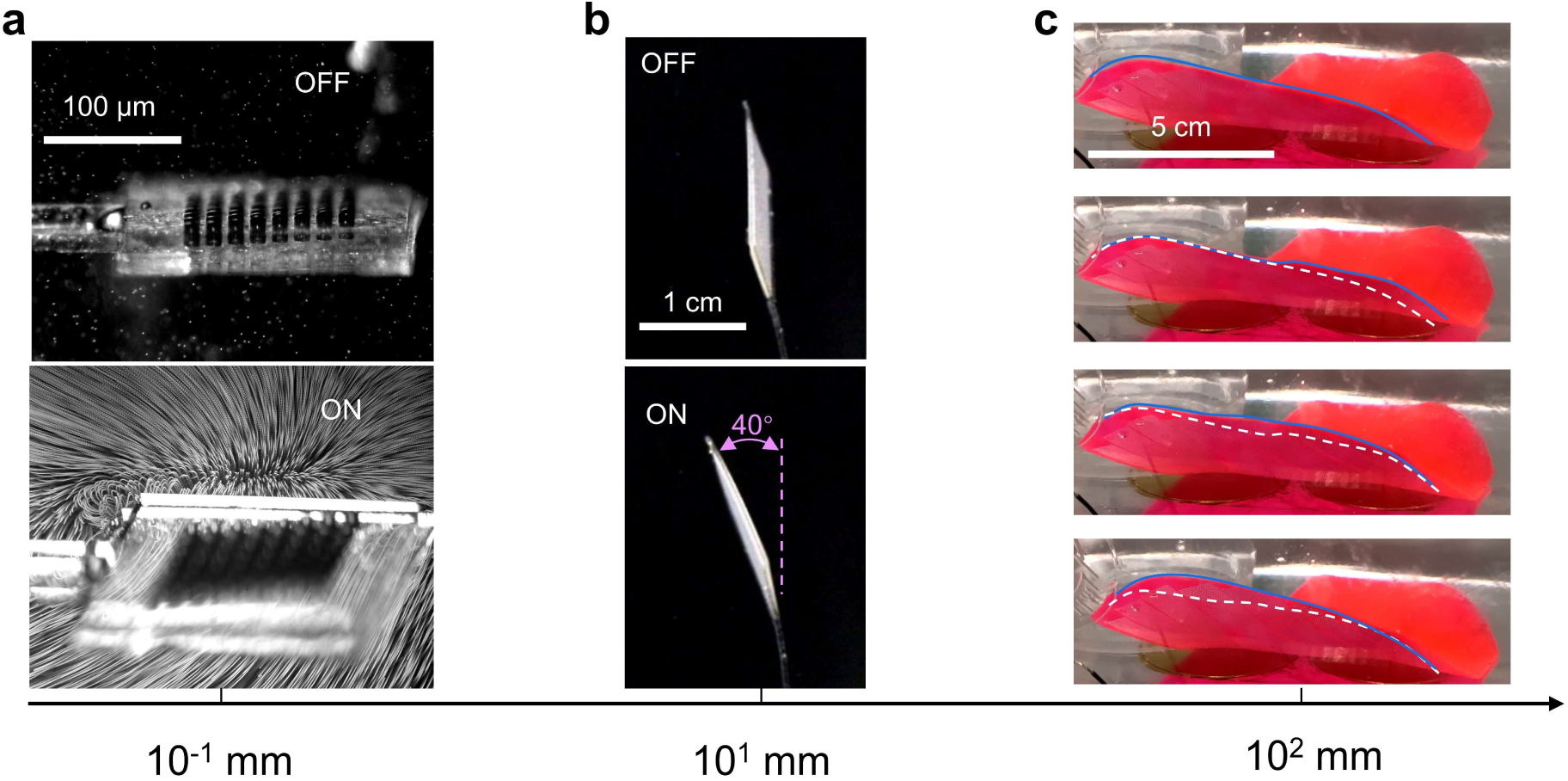
Artificial muscles functioning at scales from 10^-1^ mm to 10^2^ mm. **a,** A microscale rotator featuring an asymmetric 8 × 8 microbubble array (12 μm × 50 μm). The upper and lower panels show the microrotator with ultrasound on and off, respectively, at 95.5 kHz and 60 V_PP_. **b,** A millimeter-scale rotator with an asymmetric 400 × 200 microbubble array (12 μm × 50 μm). The upper and lower panels show the device with ultrasound on and off, respectively, under the same driving conditions. The purple dashed line indicates the original position of the artificial muscle. **c,** A macroscale stingraybot equipped with artificial muscles comprising variable-size microbubble arrays (40 μm × 150 μm, 60 μm × 150 μm, 80 μm × 150 μm, respectively), demonstrating undulatory motion under excitation (10-90 kHz, duty cycles 2 s, 120 V_PP_). The blue line marks the current location of the fin edge, while the white dashed line shows its position in the previous frame of the time-lapse image.

**Extended Data Fig. 8.**
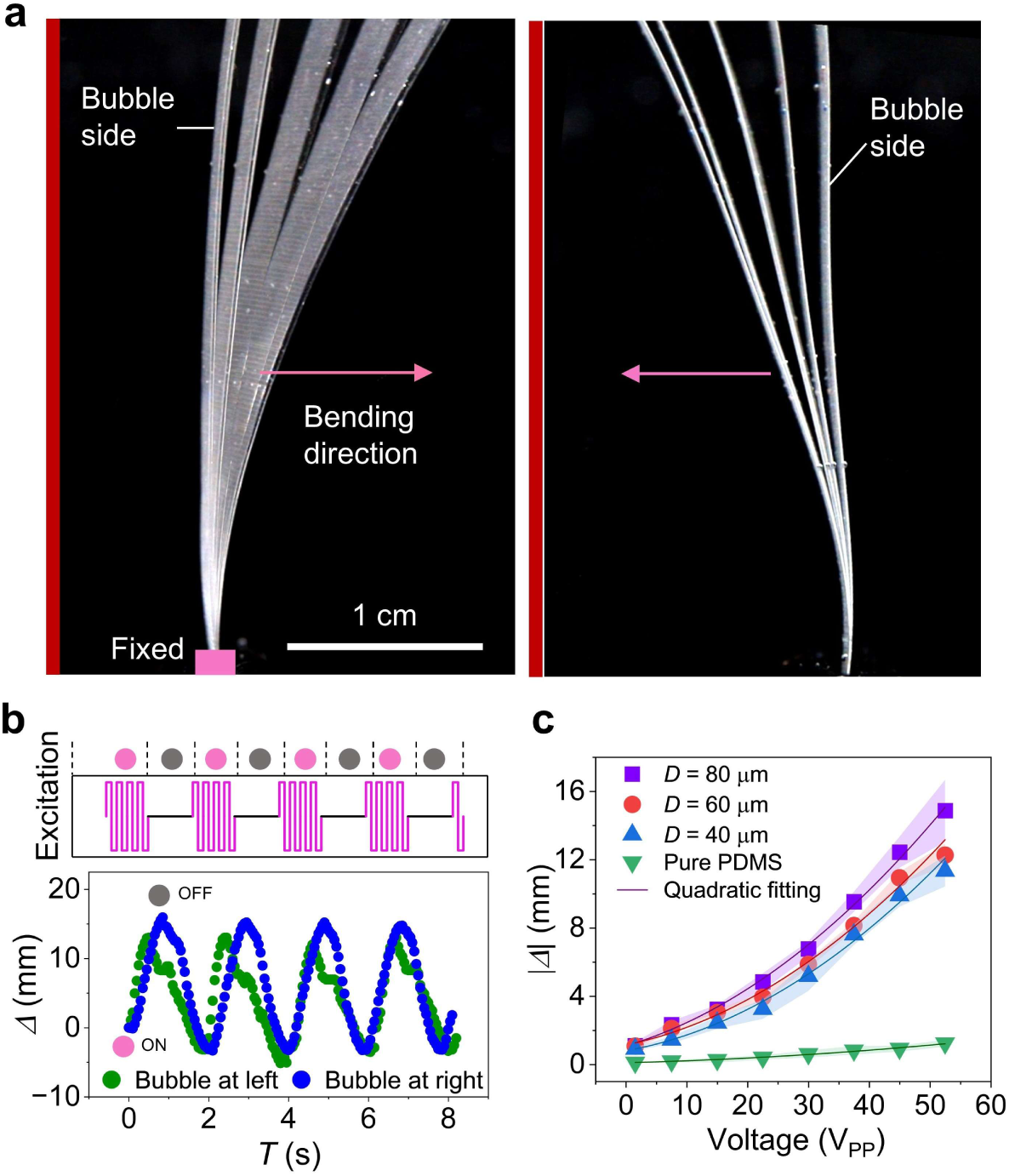
Deformation of uniform-size microbubble array artificial muscle. **a**, Time-lapse images of a uniform-size microbubble array artificial muscle with microbubbles respectively positioned at the left and right side. The pink rectangle and arrow show the fixed end and bending direction of the muscle, respectively. The red line on the left side denotes the location of the transducer. **b**, Plot of the bending amplitude of the muscle tip over multiple excitation cycles with an average repeated-positioning error of ±0.8 mm with the excitation signal shown in the top panel. The pink and gray dots correspond to the excitation ‘on’ (80.5 kHz and 52.5 V_PP_) and ‘off’ stages, respectively. The green and blue dots represent the measured deformation amplitudes when the bubbles are positioned on the left and right sides, respectively, as shown in **a**. **c**, Bending amplitude of uniform-size microbubble array artificial muscles with microbubble diameters (*D*) of 40 μm, 60 μm, and 80 μm and without microbubbles under excitation voltages from 1.5 to 52.5 V_PP_. The microbubbles have a constant depth of 50 μm. The dots and solid lines are the experimental results and quadratic fitting results, respectively. All the muscles bent in the direction as shown in the left panel of **a**.

**Extended Data Fig. 9.**
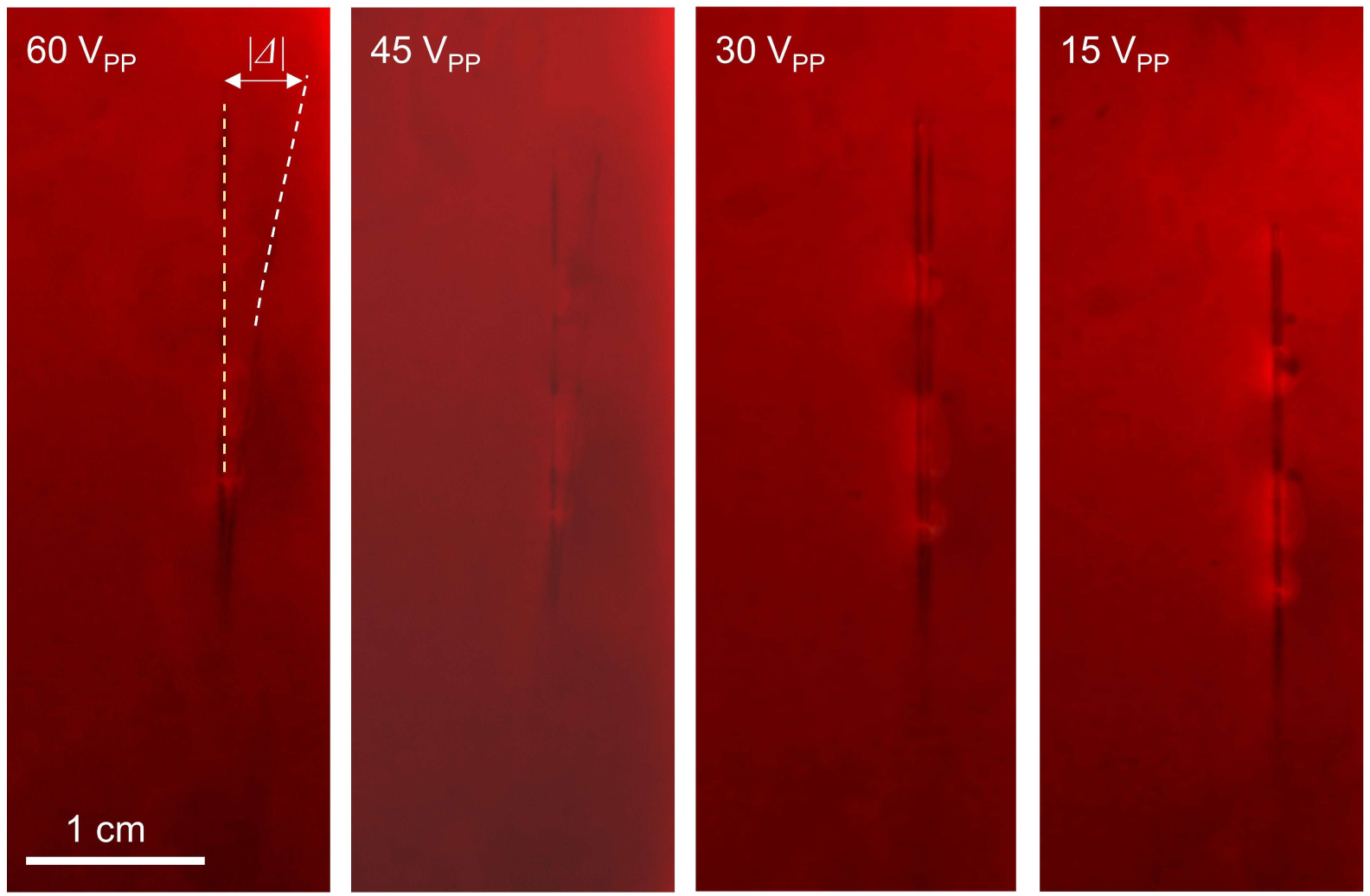
Ultrasound-induced deformation of an artificial muscle in porcine blood. An artificial muscle (12 μm × 50 μm) embedded with uniform microbubbles (diameter ∼12 μm) was immersed in 100% porcine blood, with a piezoelectric transducer bonded to the bottom of the acoustic tank and positioned 3 cm from the muscle. Final deformation (|*Δ*|) of the artificial muscle as a function of excitation amplitude—0.44 cm, 0.27 cm, 0.10 cm and 0.04 cm corresponding to 60 V_PP_, 45 V_PP_, 30 V_PP_ and 15 V_PP_, respectively—under excitation at 96.2 kHz. Yellow dashed lines indicate the initial muscle position; white dashed lines indicate the deformed position.

**Extended Data Fig. 10.**
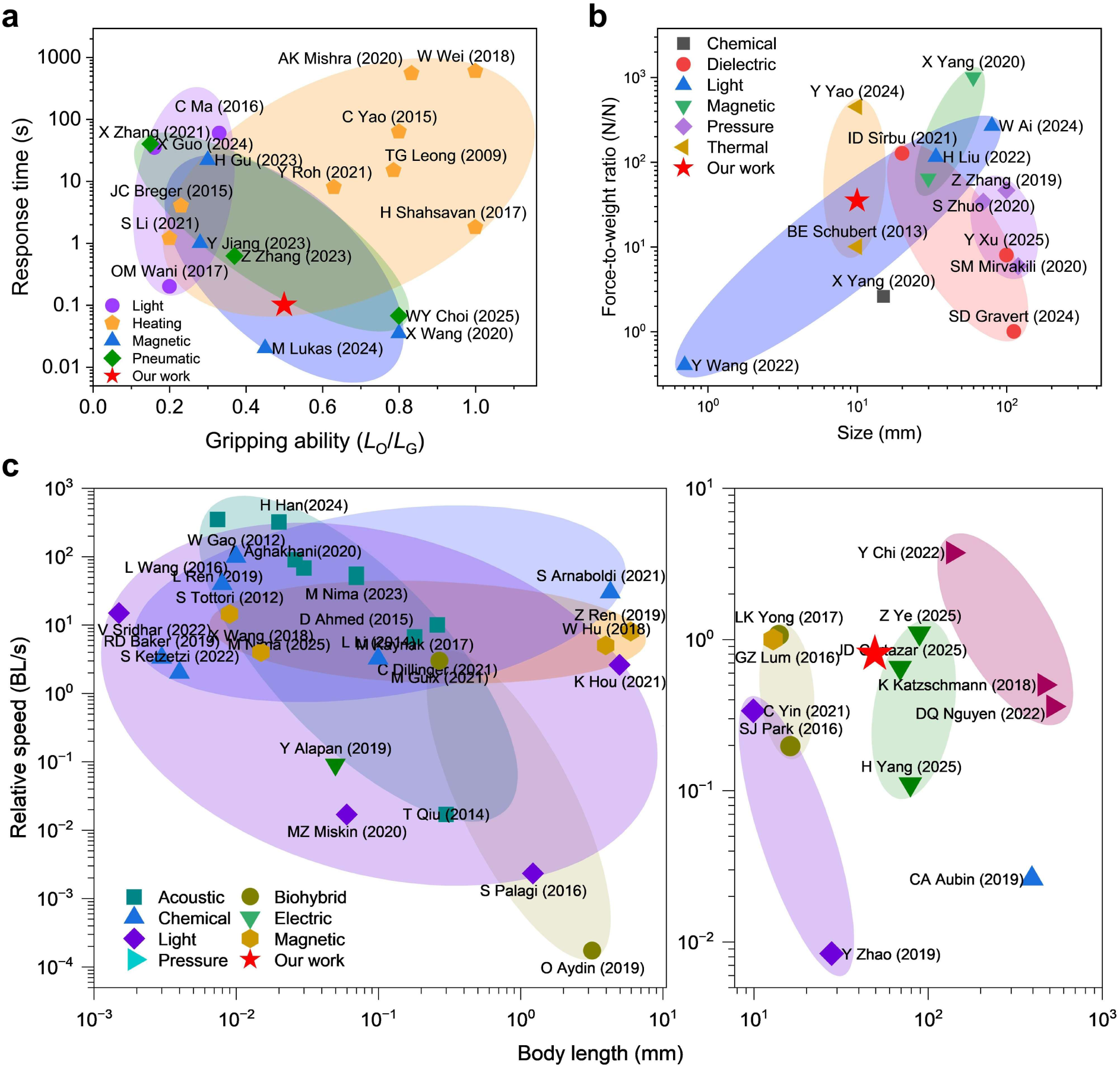
Artificial muscles performance comparison. **a,** Response time versus gripping ability (*L*_O_/*L*_G_) for grippers using different actuation methods, where *L*_O_ and *L*_G_ are the dimensions of the gripped object and gripper, respectively. **b,** Comparison of force-to-weight ratios of artificial muscles versus dimensions for various actuation methods. **c,** Comparison of relative swimming speeds (body lengths per second) of swimmers with different actuation mechanisms across various scales (microscale to macroscale). Shaded regions (convex hulls) indicate typical performance ranges; representative studies are labeled by author and year.

## Notes

### Competing Interest Statement

The authors have declared no competing interest.

### Summary of Updates

We added additional ex vivo experiments.

