## Supplementary Material_2 for "Ultrasound-Driven Programmable Artificial Muscles"

**The PDF file includes:**

Supplementary Notes
Supplementary Figures 1 to 27
Supplementary Tables 1 to 3
Legends for Supplementary Videos 1 to 18
Supplementary References

**Other Supplementary Materials for this manuscript include the following:**

Supplementary Videos 1 to 18

### Table of Contents

Supplementary Notes

Supplementary Fig. 1 | Design of artificial muscles and stingraybot.

Supplementary Fig. 2 | Artificial muscle thickness versus PDMS spin coating speed.

Supplementary Fig. 3 | Deformations of a uniform-size microbubble array artificial muscle with various transducer positions.

Supplementary Fig. 4 | Calculated force intensities.

Supplementary Fig. 5 | Deformation ($|\Delta|$) of a uniform-size microbubble array artificial muscle with varying bubble spacings.

Supplementary Fig. 6 | Deformation of a variable-size microbubble array artificial muscle under continuous and sweeping-frequency ultrasound excitation.

Supplementary Fig. 7 | Measured maximum streaming of three different bubble shapes.

Supplementary Fig. 8 | Deformation of a uniform-size microbubble array artificial muscle with confocal ultrasound excitation.

Supplementary Fig. 9 | Robustness analysis of the microbubbles at different time scales under sweeping-frequency ultrasound excitation.

Supplementary Fig. 10 | Experimental setup and streaming of membrane-sealed bubbles.

Supplementary Fig. 11 | Deformation ($|\Delta|$) of a uniform-size microbubble array artificial muscle under different excitation distances ($D$).

Supplementary Fig. 12 | Fabrication process of the soft artificial muscle.

Supplementary Fig. 13 | Microscopic top and side views of microbubble arrays.

Supplementary Fig. 14 | Experimental setup for microscope characterization of microbubble arrays and corresponding acoustic pressure field simulation.

Supplementary Fig. 15 | Experimental setup for artificial muscle actuation and corresponding acoustic pressure field isosurface simulation.

Supplementary Fig. 16 | Deformation of the artificial muscle under different waveform inputs as a function of applied voltage.

Supplementary Fig. 17 | Measurement of microstreaming velocity by PIV.

Supplementary Fig. 18 | Swimming distances of a stingraybot under different excitation conditions.

Supplementary Fig. 19 | Deformation of a robotic shark tail with the uniform-size microbubble array artificial muscle.

Supplementary Fig. 20 | Deformation of an artificial muscle in liquid within different ratios of glycerol.

Supplementary Fig. 21 | Effect of surrounding solid media on artificial muscle deformation under ultrasound excitation.

Supplementary Fig. 22 | Actuation of the artificial muscle behind porcine ribs.

Supplementary Fig. 23 | Temperature variation for different ultrasound frequencies versus time.

Supplementary Fig. 24 | A microbubble-array soft gripper captured a 35 times heavier PDMS block.

Supplementary Fig. 25 | Modeling of the uniform-size microbubble array artificial muscle.

Supplementary Fig. 26 | Deformation of an artificial muscle under ultrasound excitation.

Supplementary Fig. 27 | Comparison of artificial muscle deformation between experimental results and analytical calculations.

Supplementary Table 1 | Comparison data of the response time and gripping ability of grippers for various actuation mechanisms.

Supplementary Table 2 | Comparison data of force-to-weight ratios of artificial muscles versus dimensions for various actuation mechanisms.

Supplementary Table 3 | Comparison data of relative swimming speeds of swimmers relative to their body length for various actuation mechanisms.

Legends for Supplementary Videos

Supplementary References

### Supplementary Notes

#### Supplementary Note 1. Thrust force developed by a single oscillating microbubble

In this work, our primary goal is focusing on the thrust force generated by oscillating microbubbles, omitting other complex interactions[S1-S4]. Experimental results confirm that thrust is the dominant actuation force driving the artificial muscle. Considering an axisymmetric microbubble trapped in a PDMS cavity (cylindrical cavity with radius $R_c$ and depth $h$, as shown in the inset of **Supplementary Fig. 25**) and surrounded by fluid with the mass density $\rho$, the resonance frequency of the microbubble is estimated as $f_0 = \frac{1}{2\pi}\sqrt{\frac{3\pi\gamma P_0}{8R_c h\rho}}$[63], where $\gamma$ is the adiabatic index (1.4 for air) and $P_0$ is the atmospheric pressure. Estimating the resonance frequency of the microbubble aids in the design and measurement of its resonance characteristics.

For an analytical description of the force generated by a single bubble, we present the thrust force's scaling and show how to estimate the force given the surface streaming velocity. For this purpose, we apply the theoretical framework developed by Spelman and Lauga[68,69]. We assume incompressibility of the water and $\epsilon \ll \delta \ll 1$. Here, $\epsilon$ is the relative bubble oscillation amplitude $\epsilon = a/R$, for the bubble's (center) displacement $a$ and curvature radius $R$, and $\delta = (2\eta/\omega)^{1/2}/R$ is the relative viscous penetration length with the kinematic viscosity $\eta$ of the liquid and the ultrasound's (angular) frequency $\omega$.

The incompressibility assumption implies an infinite speed of sound, which leads to neglection of the radiation force that results from the momentum transfer of the sound wave to the bubble when the sound wave is scattered at the bubble. This assumption is necessary to obtain an analytical result for the thrust force and valid because the force corresponding to the acoustic streaming is dominant compared to this radiation force – although they can be of the same order of magnitude as can be seen in **Supplementary Fig. 26**. While the assumption $\epsilon \ll \delta \ll 1$ usually holds, it is violated if $\epsilon$ is too large, which can be the case when the ultrasound frequency approaches the resonance frequency. For describing an oscillating bubble on a beam, Spelman and Lauga's work[65,66] gives the force on the beam as

$$F = \alpha\rho R^4\omega^2\epsilon^2, \tag{S1}$$

where $\alpha$ is a purely microbubble oscillation shape-dependent prefactor and $\epsilon$ depends on the incoming ultrasound frequency and intensity as well as on the microbubble oscillation shape. Since the exact value of $\alpha$ is unknown in our case, we calculate $F$ using some approximations.

First of all, we deviate from our model sketched in the inset of **Supplementary Fig. 25** and bend the PDMS-water surface so that the total water surface is a sphere of the same curvature radius as the (resting) bubble. At the end of our calculation, we will further assume the bubble radius to be large, thereby absorbing the error of the assumption made here.

A non-dimensionalization is carried out canonically using $1/\omega$ as the time scale and $R$ as the length scale. We describe the oscillating surface of the sphere by a radial coordinate $r$ and an angular coordinate $\Theta$ (see **Supplementary Fig. 25**). As the (resting) sphere's azimuthal angle $\theta$ is used as a bubble-surface coordinate, the deformation of the bubble in radial and angular direction is time-locally given as

$$r(\theta) = 1 + \epsilon\, r^{(1)}(\theta)\, \mathrm{e}^{\mathrm{i}(t+\pi/2)} + \epsilon^2 r^{(2)}(\theta)\, t + O(\epsilon^3), \tag{S2a}$$
$$\Theta(\theta) = \theta + \epsilon\Theta^{(1)}(\theta)\, \mathrm{e}^{\mathrm{i}(t+\pi/2)} + \epsilon^2\Theta^{(2)}(\theta)\, t + O(\epsilon^3). \tag{S2b}$$

An expansion of $\Theta^{(2)}$ is given by

$$\Theta^{(2)}(\mu) = \sum_{n=1}^{\infty} \alpha_n \frac{\int_\mu^1 P_n(x)\mathrm{d}x}{\sqrt{1-\mu^2}}, \tag{S3}$$

where $P_n$ is the $n$-th Legendre polynomial and $\mu$=cos($\theta$) is the height relative to the center of curvature. Due to the orthogonality of the Legendre polynomials, the expansion coefficients are given by

$$\alpha_n = -\frac{2n+1}{2}\int_{-1}^{1} P_n(\mu)\frac{\mathrm{d}}{\mathrm{d}\mu}\left(\sqrt{1-\mu^2}\,\Theta^{(2)}(\mu)\right)\mathrm{d}\mu. \tag{S4}$$

Spelman and Lauga now give the non-dimensional thrust force as a sum of two terms with

$$\alpha = 2\pi\alpha_1 + \Gamma. \tag{S5}$$

The first term contains the deformation coefficient $\alpha_1$, whereas the second term $\Gamma$ is lengthy and neglected here to obtain a simple approximative result. This approximation is in line with the fact that a precise measurement of the surface oscillations in experiments is challenging due to limitations in temporal and spatial resolution.

Applying $P_1(\mu)=\mu$, partial integration, and a coordinate transformation with $\mu=\cos\theta$, $\alpha_1$ can be written as

$$\alpha_1 = \frac{3}{2}\int_0^\pi \sin^2(\theta)\,\Theta^{(2)}(\theta)\mathrm{d}\theta. \tag{S6}$$

If we neglect Stokes drift, we obtain $\Theta^{(2)}(\theta)$ via the tangential component of the surface streaming velocity. Next, we approximate the tangential surface streaming velocity $v$ to be constant along the bubble surface:

$$\epsilon^2\,\Theta^{(2)}(\theta) = \frac{v}{R\omega}\chi_{0\leq\theta\leq\theta_c}. \tag{S7}$$

Here, $\chi$ is the indicator function and $\theta_c$ is the angle corresponding to the bubble boundary. The prefactor $1/(R\omega)$ non-dimensionalizes $v$ which is measured using tracer particles. If $R$ is large, then $\theta_c$ is small, so the coefficient $\alpha_1$ simplifies to

$$\alpha_1 \approx \frac{3}{2}\int_0^{\theta_c}\theta^2 d\theta\frac{v}{R\omega\epsilon^2} = \frac{{\theta_c}^3}{2}\frac{v}{R\omega\epsilon^2}. \tag{S8}$$

Thus, the (dimensional) thrust force exerted by one bubble on the artificial muscle can be estimated as

$$F \approx \pi\,\rho\omega{\theta_c}^3R^3v \approx \pi\,\rho\omega{R_c}^3v, \tag{S9}$$

where we introduced the cavity radius $R_c = R\sin(\theta_c)$ and again applied the large-$R$ assumption. Since we expect $v$ to have a negative sign (see the streaming inset in **Supplementary Fig. 25**), we expect a downwards oriented force leading to a bending away from the microbubble array.

**Supplementary Note 2. Deformation of the artificial muscle**

We model the artificial muscle as a slender, inextensible beam divided into discrete segments corresponding to patterned microbubble arrays (**Fig. 2e**, **Supplementary Figs. 6** and **25**). The beam of length $L$ is parameterized by the coordinate $s \in [0,L]$. Due to symmetry and the nature of the applied thrust, deformation is confined to a plane and fully described by the local bending angle $\theta(s)$.

The strain energy of the slender beam due to bending is expressed as

$$V = \frac{1}{2}\int_0^L EI\theta'^2 \mathrm{d}s. \tag{S10}$$

with $E$ as Young's modulus and $I$ as the second moment of area. The clamped-free boundary conditions are,

$$\theta(0) = 0,\ \theta'(L) = 0,\ \delta\theta(0) = 0, \tag{S11}$$

The variation in strain energy is given by:

$$\delta V = \int_0^L EI\theta'\delta\theta'\mathrm{d}s = -\int_0^L (EI\theta')'\delta\theta\mathrm{d}s. \tag{S12}$$

To evaluate the virtual work done by the external force, we compute the beam's deformation vector $(\delta x, \delta y)$:

$$\begin{pmatrix} x \\ y \end{pmatrix}(s) = \int_0^s \begin{pmatrix} \cos(\theta(s')) \\ \sin(\theta(s')) \end{pmatrix} \mathrm{d}s' \tag{S13}$$

$$\delta \begin{pmatrix} x \\ y \end{pmatrix}(s) = \int_0^s \begin{pmatrix} -\sin(\theta(s')) \\ \cos(\theta(s')) \end{pmatrix} \delta\theta(s')\mathrm{d}s'\ . \tag{S14}$$

Assuming a perpendicular force density $P_{\mathrm{u}i} = n_i F_i / S_i$, where $n_i$, $F_i$ and $S_i$ denote the number of bubbles, the thrust force of a single bubble, and the coverage area at the $i$-th segment, respectively, the external virtual work simplifies as:

$$\begin{aligned} \delta W &= \int_0^L P_u(s) \begin{pmatrix} -\sin(\theta(s)) \\ \cos(\theta(s)) \end{pmatrix} \int_0^s \begin{pmatrix} -\sin(\theta(s')) \\ \cos(\theta(s')) \end{pmatrix} \delta\theta(s')\mathrm{d}s'\mathrm{d}s \\ &= \int_0^L \delta\theta(s) \int_s^L P_u(s')\cos(\theta(s) - \theta(s'))\mathrm{d}s'\mathrm{d}s. \end{aligned} \tag{S15}$$

From the principle of minimum potential energy:

$$\delta V - \delta W = 0, \tag{S16}$$

and arbitrariness of $\delta\theta$ it follows that

$$EI\theta'' + (EI)'\theta' + \int_s^L P_u(s')\cos(\theta(s) - \theta(s'))\mathrm{d}s' = 0 \qquad \text{(S17)}$$

which is the second order integro-differential equation governing $\theta$. Equation (S17) can be solved numerically for $s$ going from $L$ to 0, where the boundary condition at 0 is obeyed afterwards by adding some constant to $\theta$. But we want to give an analytical expression. As a first (linear) approximation of small differences in $\theta$ along the beam, one can write

$$EI\theta'' + (EI)'\theta' + \int_s^L P_u(s')\mathrm{d}s' = 0. \qquad \text{(S18)}$$

To coordinate with the variable-size microbubble array artificial muscle, we subdivide the beam into $N$ segments with endpoints $s_j$ ($j$=1…$N$) and lengths $L_j$ ($j$=1…$N$), and replace the arguments of both $\theta$ in the integrand by their nearest higher segment endpoints. For $N \to \infty$, with segment length vanishing, the approximation becomes exact. We now assume $E$, $I$, and $P_u$ to be constant along each segment. From Eq. (S17), we get the interface condition: $EI\theta'$ must be continuous. In each segment $s_{j-1} \le s \le s_j$ on the other hand, $\theta$ is governed by

$$E_jI_j\theta'' + (s_j - s)P_{u,j} + \sum_{i=j+1}^{N} P_{u,i}L_i\cos\left(\theta(s_i) - \theta(s_j)\right) = 0, \qquad \text{(S19)}$$

where the cosine term reflects our improved approximation. This system of equations is solved for $j$=$N$ up to $j$=1. The initial conditions for the first step are $\theta'(L)$=0 and $\theta$ arbitrary. Later, continuity of $EI\theta'$ is enforced and in the end, $\theta(0)$=0 makes us add a constant on $\theta$ everywhere. Obviously, $\theta$ is a third order polynomial in s in every segment.

Applying the aforementioned algorithm for $N$=3, we successively get the solution for $\theta_i$ using the auxiliary variables $C_i, D_i, \varphi_i, \vartheta_i$ as follows:

$$\begin{aligned}
D_2 &= \frac{P_{u,3}}{2}{L_3}^2, \\
\varphi_2 &= \frac{P_{u,3}}{6E_3I_3}{L_3}^3, \\
C_2 &= P_{u,3}L_3\cos(\varphi_2), \\
D_1 &= \frac{P_{u,2}}{2}{L_2}^2 + C_2L_2 + D_2, \\
\varphi_1 &= \frac{1}{E_2I_2}\left(\frac{P_{u,2}}{6}{L_2}^3 + \frac{C_2}{2}{L_2}^2 + D_2L_2\right), \\
C_1 &= P_{u,2}L_2\cos(\varphi_1) + P_{u,3}L_3\cos(\varphi_1 + \varphi_2), \\
\vartheta_1(s + s_1) &= \frac{1}{E_1I_1}\left(\frac{P_{u,1}s^3}{6} - \frac{C_1s^2}{2} + D_1s\right)\theta_1(s) = \vartheta_1(s) - \vartheta_1(0), \\
\vartheta_2(s + s_2) &= \frac{1}{E_2I_2}\left(\frac{P_{u,2}s^3}{6} - \frac{C_2s^2}{2} + D_2s\right), \\
\theta_2(s) &= \vartheta_2(s) - \vartheta_2(s_1) + \theta_1(s_1), \\
\vartheta_3(s + s_3) &= \frac{P_{u,3}s^3}{6E_3I_3}, \\
\theta_3(s) &= \vartheta_3(s) - \vartheta_3(s_2) + \theta_1(s_2).
\end{aligned} \qquad \text{(S20)}$$

The resulting $Y$-direction deformation as a function of $s$ in $i$-th segment is then given by

$$\Delta(s) = \int_0^s \sin(\theta(s))\,\mathrm{d}s\,. \tag{S21}$$

Our model is also applicable of artificial muscles featuring uniform-size microbubble array. For $N$=1, the rotation angle at location $s$ is:

$$\theta(s) = \frac{P_u}{6EI}\left((s-L)^3 + L^3\right), \tag{S22}$$

And the deformation at end point is

$$\Delta_y = \int_0^L \sin(\theta(s))\,\mathrm{d}s. \tag{S23}$$

**Supplementary Note 3. Quantitative comparison**

To enable quantitative comparison with experiments, we consider the measured results of the uniform-size microbubble array artificial muscle, approximately subjected to a uniformly distributed orthogonal thrust $P_u$. As an example, we calculate the thrust force for a microbubble with a radius of 40 μm using **Eq. (S9)**. Based on the experimentally measured velocity of 2.5 mm/s at an excitation frequency of 27.6 kHz and input voltage of 60 $V_{PP}$ (**Extended Data Fig. 3b**), and assuming a water density of 1000 kg/m$^3$, the resulting thrust force is 13.9 nN per bubble. Scaling this to an array of approximately 10,000 uniformly sized microbubbles on a 30 mm × 5 mm artificial muscle yields a total force of ~139 μN, corresponding to a force intensity of 0.93 μN/mm$^2$ under uniform distribution. Additional calculations for microbubbles with radii of 20, 30, and 40 μm across a voltage range of 1–60 $V_{PP}$ are shown in **Supplementary Fig. 26**, with maximum force intensities ranging from 0.83 to 1.21 μN/mm$^2$ at 60 $V_{PP}$.

Using Eqs. (S9), (S22), and (S23), we calculated the deformation of the artificial muscle by incorporating the experimentally measured streaming velocities of 40 μm-radius microbubbles at different voltages, and compared the results with experimental data (**Supplementary Fig. 27**). The relative error is approximately 70%, which is reasonable given that the model neglects the cavities within the artificial muscle, leading to an overestimation of stiffness and thus an underestimation of deformation.

**Supplementary Note 4. Simulation setups**

**1. Acoustic pressure distribution in smaller chamber**

We employed a three-dimensional geometry modeled with the same dimensions as the experimental setup. A circular piezoelectric transducer (27 mm × 0.54 mm) is affixed to the glass substrate (24 mm×60 mm×0.18 mm). A square PDMS acoustic chamber (10 mm×10 mm×5 mm) is positioned in the front of the transducer, which is filled with deionized water and covered with a cover glass (22 mm×22 mm×0.18 mm). The simulation domain included the following materials: water (built-in material), cover glass and glass substrates (CompoGlass, built-in material), a piezoelectric transducer (PZT-5H, built-in material) (with parameters extracted from experiments). The physics modules comprised solid mechanics, electrostatics, frequency-domain pressure acoustics, heat transfer in solids and fluids, and creeping flow. Multiphysics couplings accounted for acoustic–structure boundary interactions, acoustic streaming domain couplings, acoustic streaming boundary couplings, and the piezoelectric effect. A user-controlled (fine) mesh was applied, and the excitation frequency was set to 33 kHz and the voltage as 60 $V_{PP}$.

**2. Acoustic pressure distribution in bigger chamber**

We employed a three-dimensional geometry modeled with the same dimensions as the experimental setup. A thin plastic tank with dimensions of 10 cm ×10 cm × 8 cm and a thickness of 2 mm. The circular piezoelectric transducers (27 mm × 0.54 mm) are affixed to the inside surfaces and the center of the bottom surface of the tank, which is filled with deionized water. The simulation domain included the following materials: water (built-in material), a piezoelectric transducer (PZT-5H, built-in material), plastic chamber (Polyethylene Terephthalate). The physics modules comprised solid mechanics, electrostatics, frequency-domain pressure acoustics, heat transfer in solids and fluids, and creeping flow. Multiphysics couplings accounted for acoustic–structure boundary interactions, acoustic streaming domain couplings, acoustic streaming boundary couplings, and the piezoelectric effect. A user-controlled (fine) mesh was applied, and the excitation frequency was set to 33 kHz and the voltage as 60 $V_{PP}$.

**3. Acoustic streaming**

To address the computational complexity and time constraints of 3D streaming flow simulations, we employed a simplified 2D geometric model. This model features three air bubbles (12 μm, 16 μm, and 66 μm in diameter) arranged sequentially, each size comprising five microbubbles. The displaced region shown in **Extended Data Fig. 4d** represents a zoomed-in view of an acoustic chamber embedded within a 10 mm × 10 mm water domain. The air bubbles and surrounding fluid were modeled using Thermoviscous Acoustics, coupled to a Pressure Acoustics domain with a Spherical Wave Radiation boundary condition. Surface tension effects at the bubble interface were incorporated via the Surface Tension boundary condition. A rectangular piezoelectric transducer (27 mm × 0.54 mm, PZT-5H material) was affixed to the chamber base. The simulation domain included water (built-in material) and the PZT-5H transducer. For flow tracking, 1 μm-radius particles were uniformly distributed in the chamber. Key multiphysics couplings included: Acoustic–structure boundary interactions, acoustic streaming (domain and boundary couplings), piezoelectric effects. A user-controlled fine mesh was applied, and excitation frequencies of 30 kHz, 80 kHz, and 100 kHz were tested (**Extended Data Fig. 4d**).

**4. Oscillation shape of the variable-size beam**

We employed a three-dimensional model of an artificial muscle, representing a PDMS structure with dimensions of 3 cm × 0.5 cm × 0.1 cm, simulated using the Solid Mechanics module. The micro-cavities within the muscle were simplified into three distinct arrays with a reduced number of cavities to streamline the computational model. In this simulation, the central region of the muscle was excited by applying a uniform stress (force per unit area) to each cavity, and the time-dependent solution was recorded at 0.5 s. The boundary conditions were set to fixed-free, with the fixed edges constrained, while the free edges were allowed to deform. The PDMS material properties were defined with a Young's modulus of 1 MPa and a Poisson's ratio of 0.1.

### Supplementary Figures

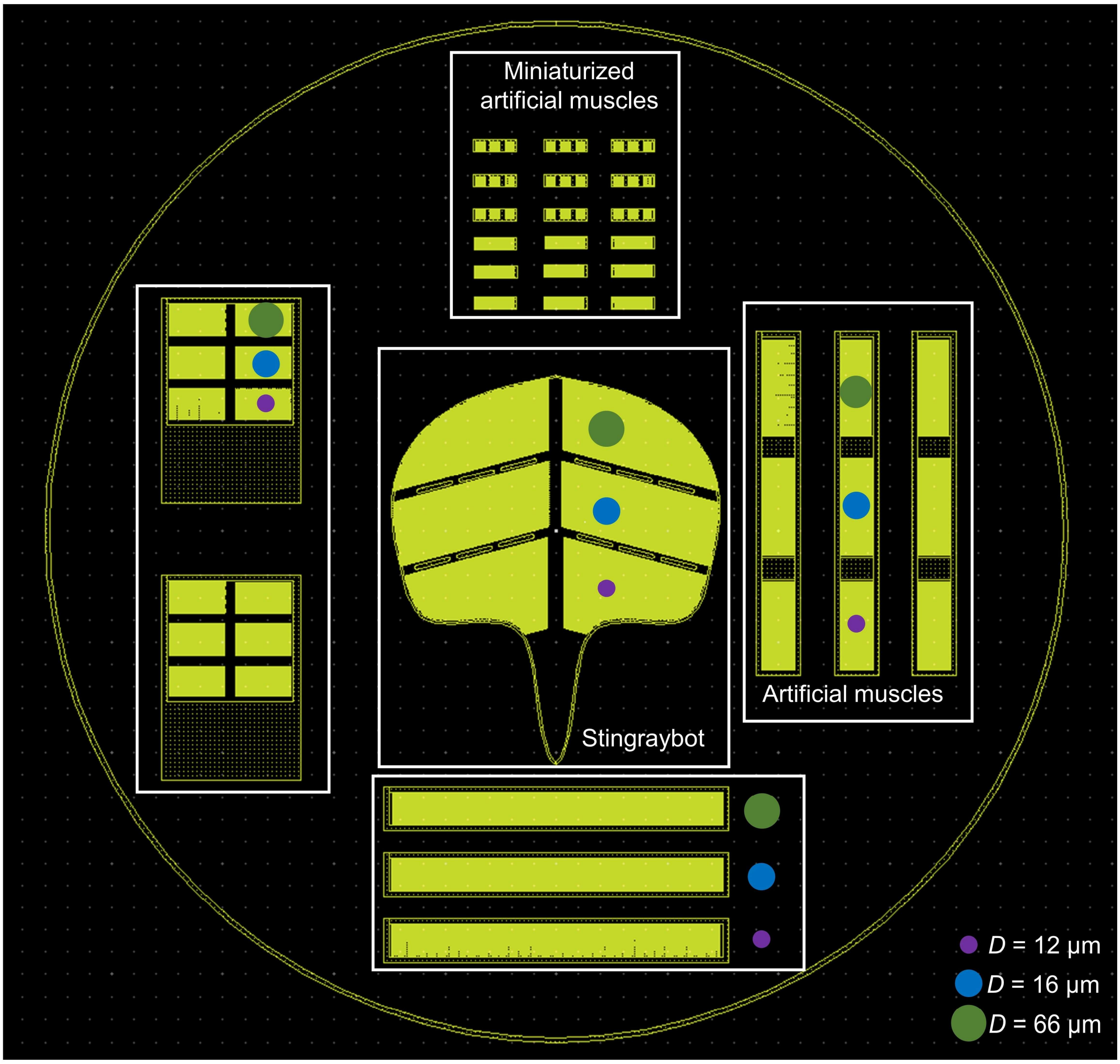


**Supplementary Fig. 1 | Design of artificial muscles and stingraybot.** At the center of the 4-inch wafer profile, a bioinspired stingraybot is depicted, comprising three distinct microbubble arrays with varying radii arranged from the head to the tail. At the bottom are three artificial muscles with uniform-size microbubble arrays. On the right are the artificial muscles with variable-size microbubble arrays. Both the artificial muscles at the bottom and the right side are 30 mm in length and 5 mm in width. The top presents the miniaturized artificial muscles, which are scaled down by a factor 0.12 compared to the muscles at the bottom and the right. However, the microbubbles of these miniaturized artificial muscles have the same radii.

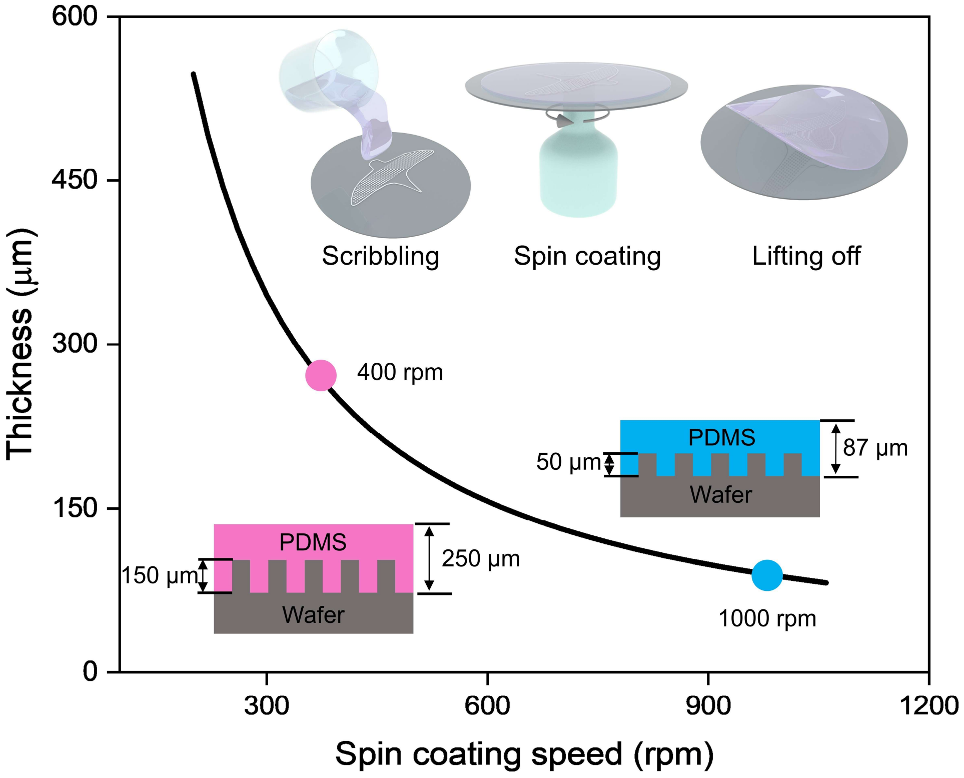


**Supplementary Fig. 2 | Artificial muscle thickness versus PDMS spin coating speed.** The black curve illustrates that the resulting thickness of the artificial muscle decreases with a higher spinning speed. The spinning speeds utilized in experiments are highlighted by two specific data points: The pink dot corresponds to a spinning speed of 400 rpm, which was employed on a wafer with 150 μm pillars, resulting in a thicker thickness of 250 μm for the artificial muscle. The blue dot represents a spinning speed of 1000 rpm, which was applied to a wafer with 50 μm pillars, yielding a thinner artificial muscle with a thickness of 87 μm. Insets demonstrate the fabrication process.

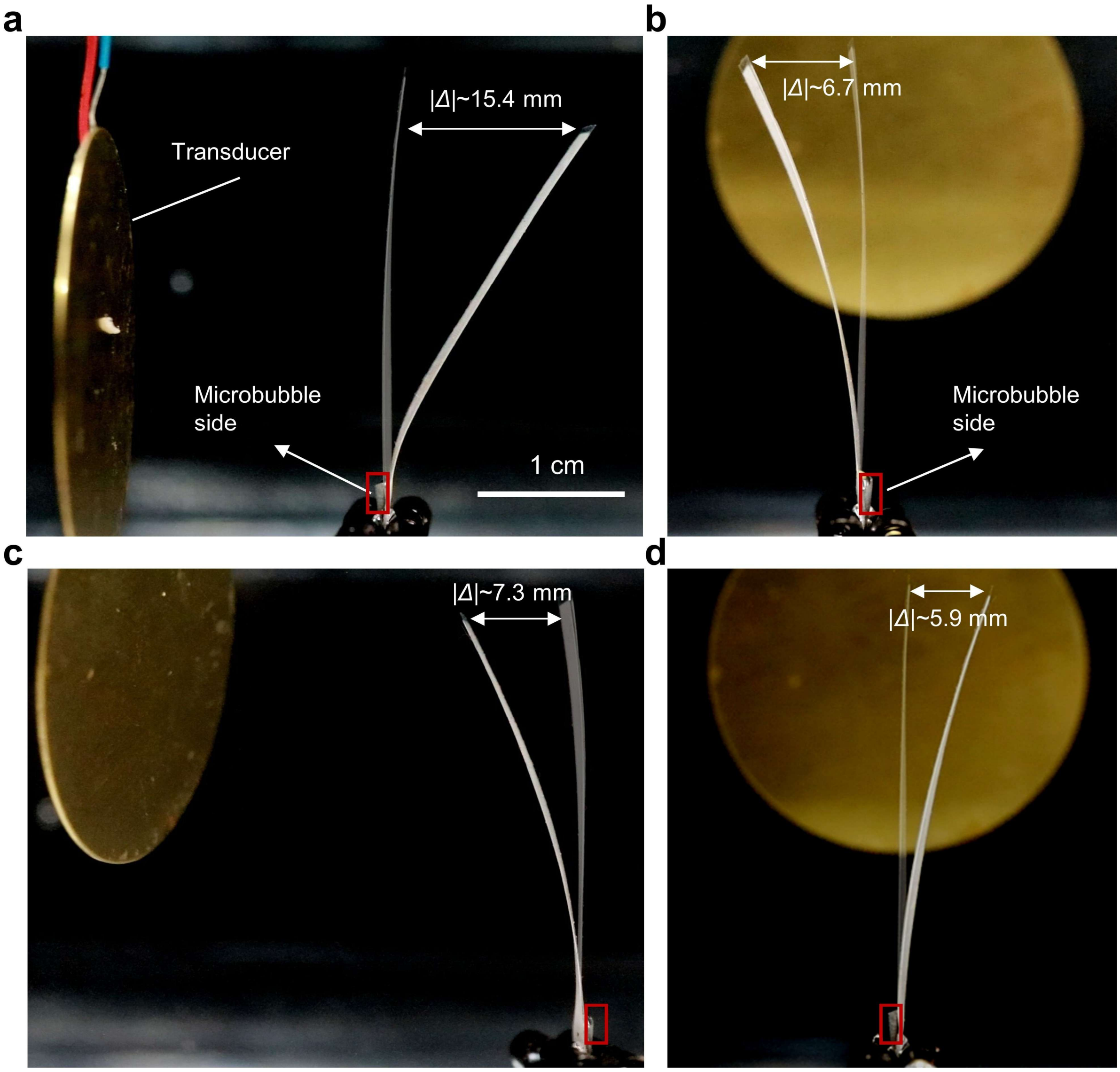


**Supplementary Fig. 3 | Deformations of a uniform-size microbubble array artificial muscle with various transducer positions. a** to **d,** The transducer was positioned with four distinct orientations: **a**, directly facing the microbubble-embedded side, **b**, opposite to it, and perpendicular to the array's left (**c**) and right (**d**) sides of the artificial muscle. The side with the microbubble array is marked by a red rectangle. The muscle demonstrated a consistent bending direction regardless of these positions, i.e., all samples bent in the direction opposite to the microbubble-array side, despite the variations in bending amplitudes. The ultrasound excitation was 80.5 kHz and 60 $V_{PP}$. In **b** and **d** the bending is due to the acoustic streaming thrust. One can expect the sound wave in **a** and **c** to push the artificial muscle to the right, but the unaltered deflection in **c** indicates that a force due to sound reflection plays only a minor role compared to the force due to acoustic streaming.

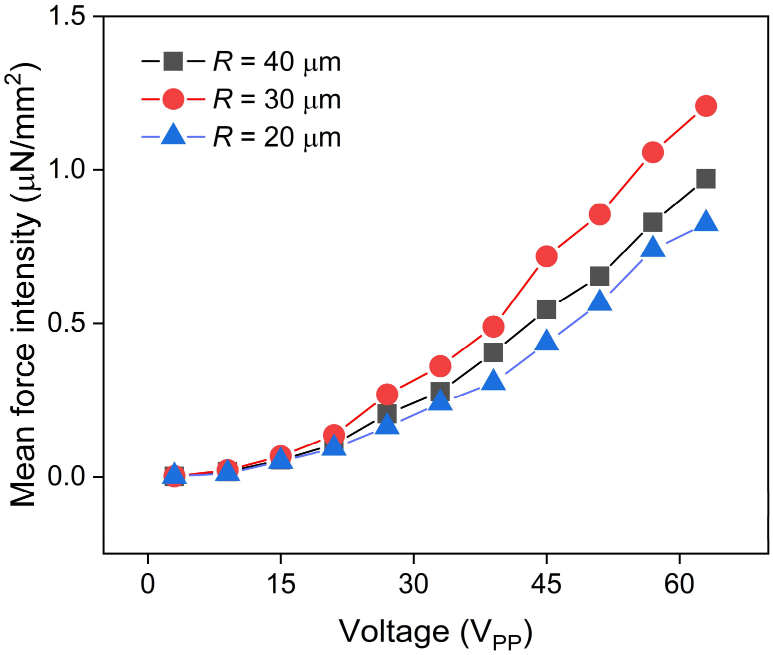


**Supplementary Fig. 4 | Calculated force intensities.** Calculated mean force intensities as a function of applied voltage. The values were obtained using Eq. (S9), incorporating experimental data from **Extended Data Fig. 6b.**

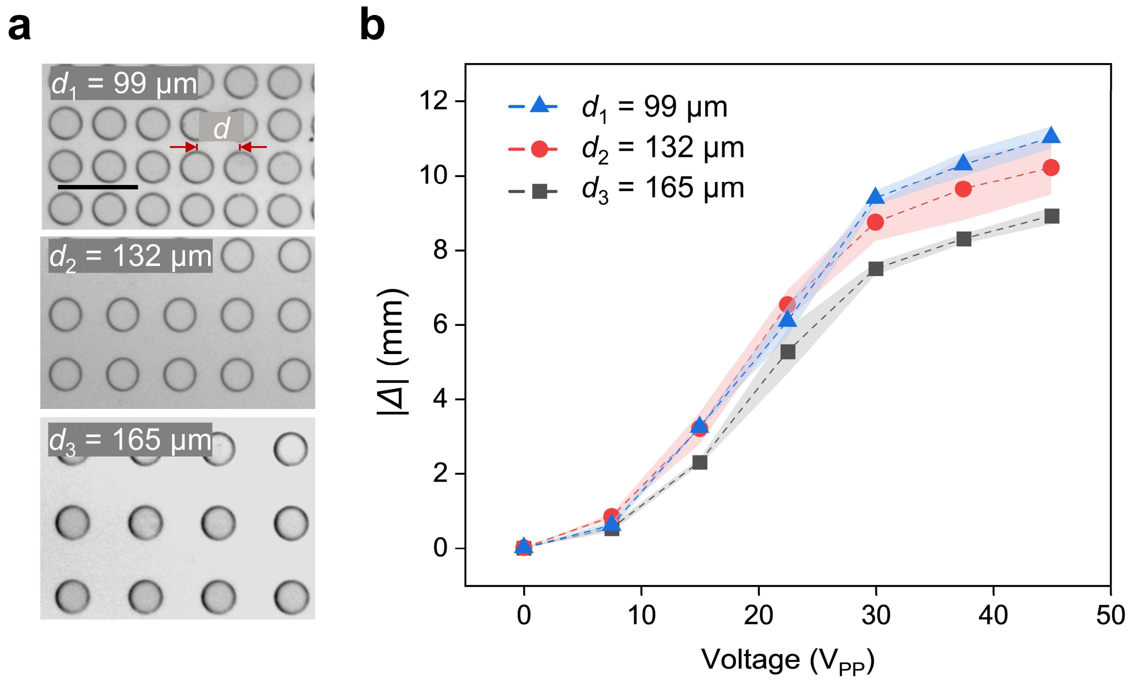


**Supplementary Fig. 5 | Deformation (|*Δ*|) of a uniform-size microbubble array artificial muscle with varying bubble spacings**. **a**, Microscopic image of the microbubble arrays (66 μm × 50 μm) utilized with different spacing ($d$ = 99 μm, 132 μm, 165 μm), where the scale bar is 200 μm. **b**, Deformation of the artificial muscles with different spacings between microbubbles as function of the applied voltage, where the input frequency is 33.2 kHz.

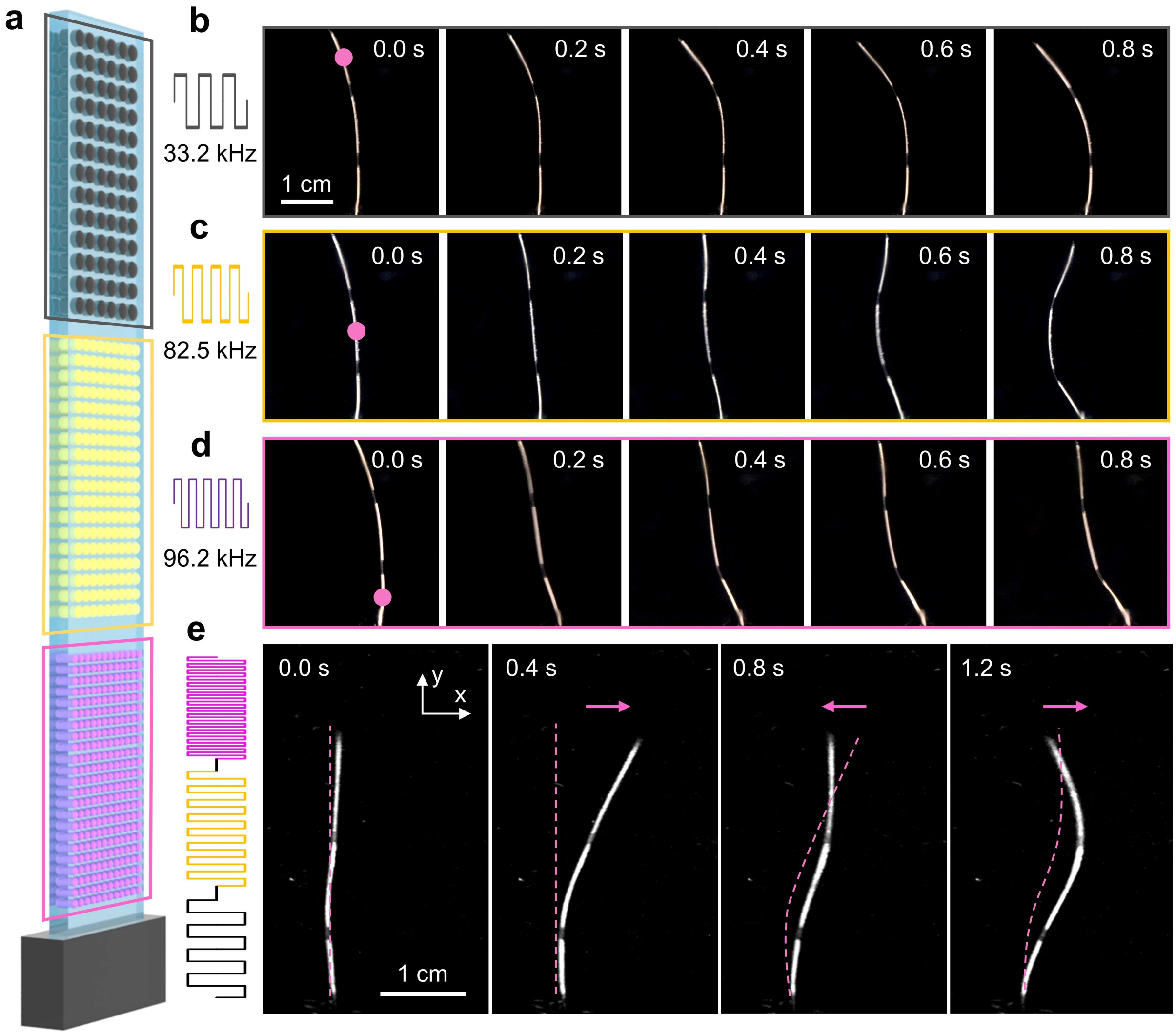


**Supplementary Fig. 6 | Deformation of a variable-size microbubble array artificial muscle under continuous and sweeping-frequency ultrasound excitation. a,** Schematic of the artificial muscle (thickness: 200 µm), comprising three different microbubble arrays, denoted by black (66 µm × 50 µm), yellow (16 µm × 50 µm), and pink (12 µm × 50 µm) colors. **b** to **d,** Time-lapse sequences of the artificial muscle's deformation when we respectively excited the top, middle, and bottom microbubble arrays, each corresponding to a specific frequency indicated by a color-coded square signal. The magenta dot denotes the excited microbubble array. **e**, A time-lapse sequence image of the artificial muscle's response to sweeping frequency ultrasound excitation, ranging from 20 kHz to 90 kHz, over a sweeping time of 2 seconds. The magenta arrow denotes the deformation direction. The magenta dotted line denotes the posture of the artificial muscle at the previous time step. The ultrasound excitation voltage for all panels is 60 $V_{PP}$.

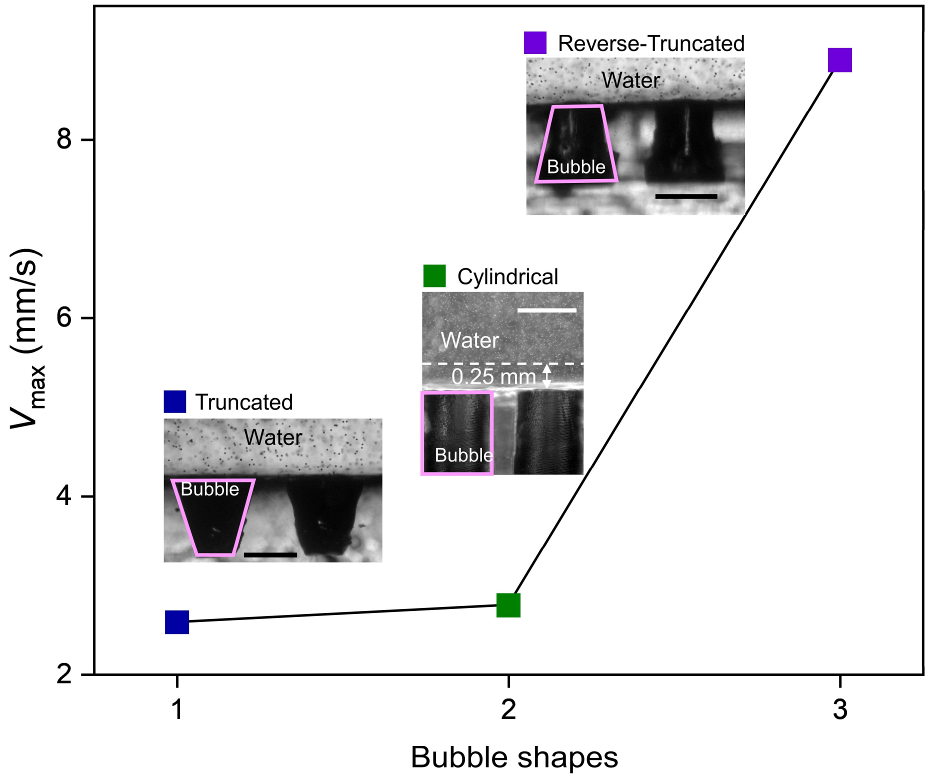


**Supplementary Fig. 7 | Measured maximum streaming of three different bubble shapes.** The blue, green, and purple points denote truncated, cylindrical, and reverse-truncated bubble shapes, respectively, with velocity measured along a line 0.25 mm from the bubble surface and 2 µm trace microparticles at 2.1 kHz and 30 $V_{PP}$ in all experiments. Scale bar: 1 mm.

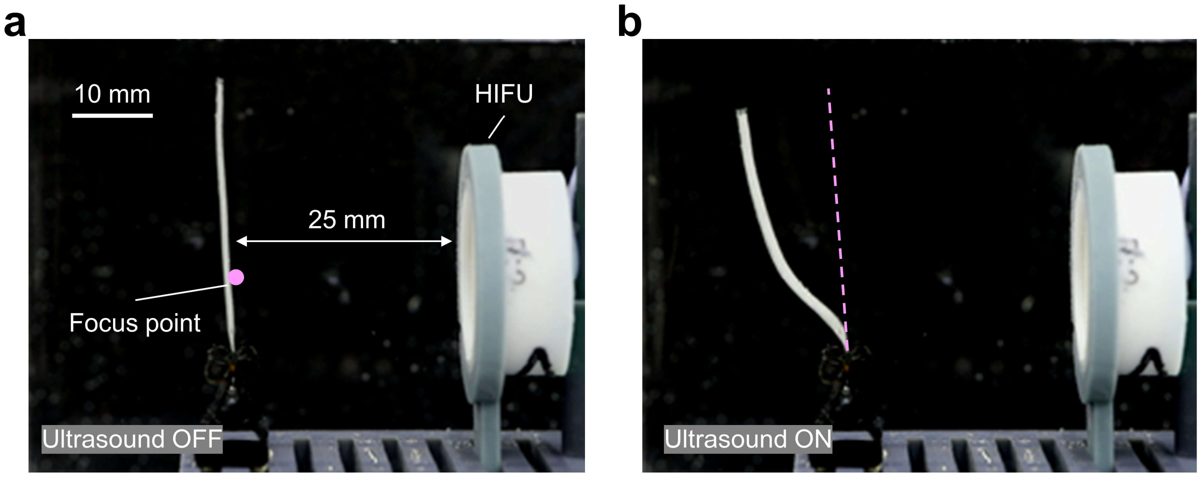


**Supplementary Fig. 8 | Deformation of a uniform-size microbubble array artificial muscle with confocal ultrasound excitation**. **a** and **b**, Artificial muscle before and after ultrasound excitation, respectively. The pink dashed line in **b** indicates the original position of the muscle prior to ultrasound excitation. The confocal ultrasound excitation was generated using a HIFU transducer with a focal distance of 25 mm, driven at 70 kHz and 45 $V_{PP}$.

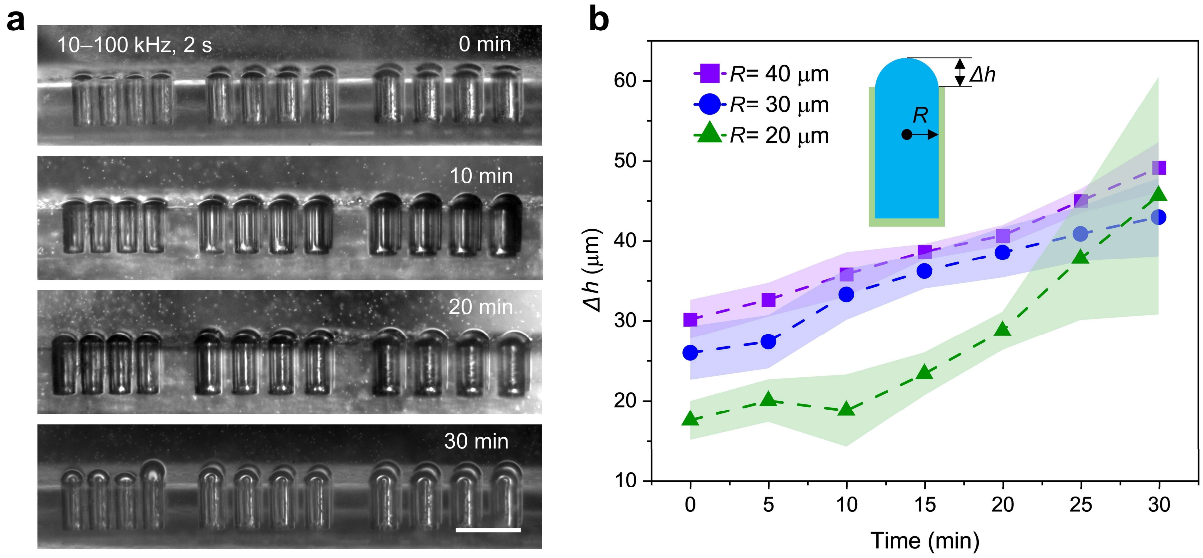


**Supplementary Fig. 9 | Robustness analysis of the microbubbles at different time scales under sweeping-frequency ultrasound excitation. a,** Microscopic images of the variable-size microbubble arrays at different excitation times, with an excitation frequency ranging from 10 to 100 kHz, a sweeping duty cycle of 2 s, and voltage at 60 $V_{PP}$. Scale bar: 200 μm. **b**, Measurement of the mean outer height change ($\Delta h$) outside the cavity for bubbles of different sizes as a function of time.

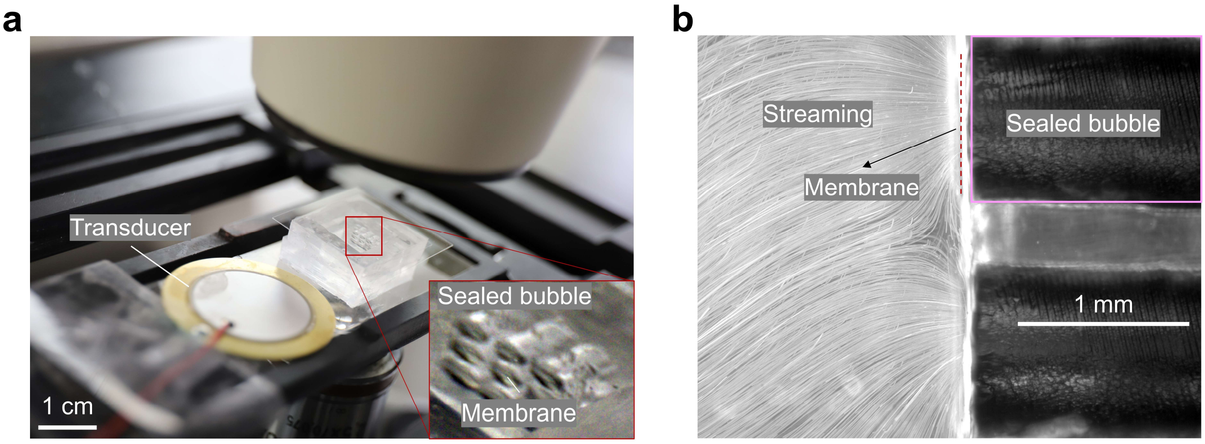


**Supplementary Fig. 10 | Experimental setup and streaming of membrane-sealed bubbles. a**, Experimental setup showing the transducer and a membrane-sealed 3 × 3 bubble array (1 mm diameter, 1.5 mm depth) mounted on a glass slide. The inset highlights the sealed bubbles. The membrane is 10 µm thick. **b**, Leftward microstreaming jetting generated from a microbubble array oscillating under ultrasound excitation demonstrated by 6 µm tracer microparticles. Excitation was applied at 2.1 kHz with a voltage of 60 $V_{PP}$. Pink rectangular represent the contour of the cross section of the sealed bubble, red dashed line indicates the location of the membrane.

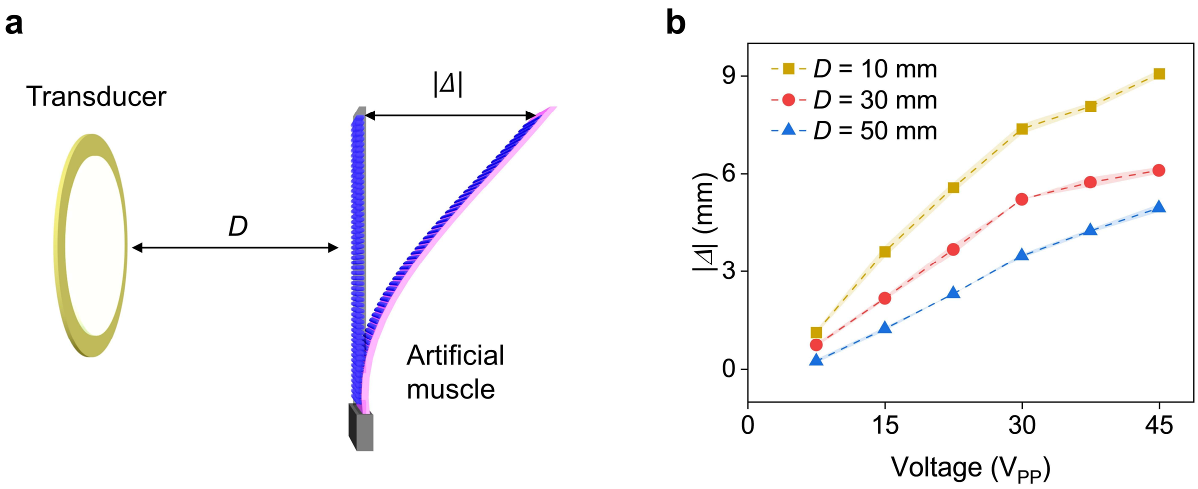


**Supplementary Fig. 11 | Deformation (|*Δ*|) of a uniform-size microbubble array artificial muscle under different excitation distances (*D*). a,** Schematic shows a transducer exciting a uniform-size microbubble array artificial muscle (12 µm × 50 µm) at varying distances, under a fixed excitation frequency of 96.2 kHz. **b,** Measured deformation as a function of voltages at varying excitation distances.

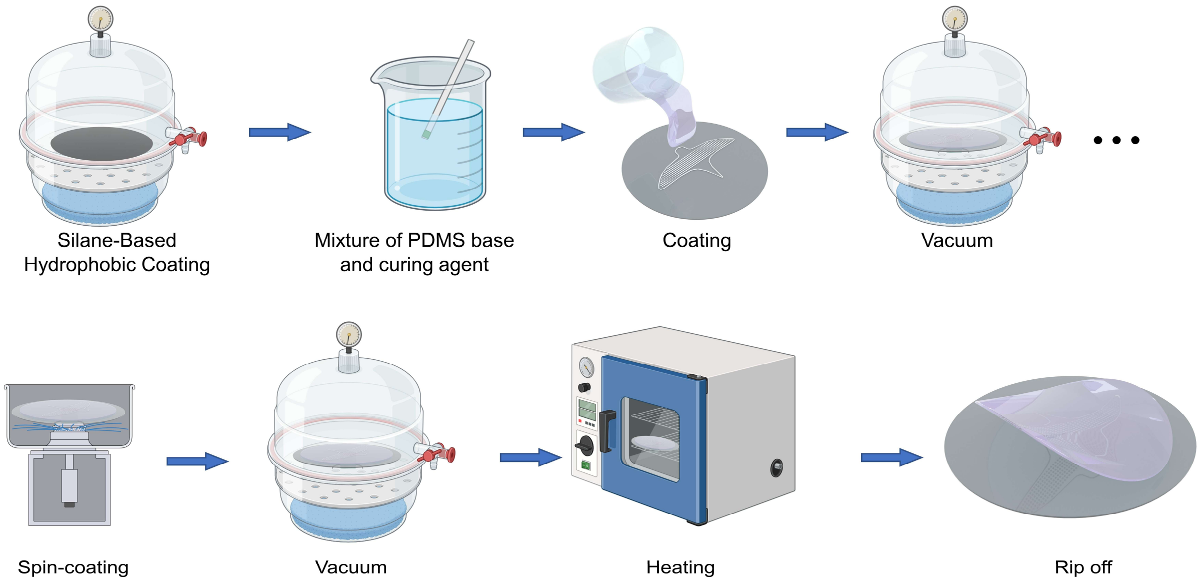


**Supplementary Fig. 12 | Fabrication process of the soft artificial muscle.** A 4-inch micropillar-patterned wafer was treated with a silane-based hydrophobic coating to enhance surface properties. A PDMS mixture, prepared with a 10:1 base-to-curing-agent ratio, was poured onto the wafer and degassed under vacuum (<1 mBar). Spin-coating was performed at various rotation speed to get various thickness, respectively. After a second vacuum treatment, the PDMS was cured in a sequential heating process (1 hour at 60°C, 1 hour at 80°C, and 1 hour at 100°C). The cured PDMS membrane was then peeled off, yielding a uniform soft layer suitable for artificial muscle and soft robotic applications.

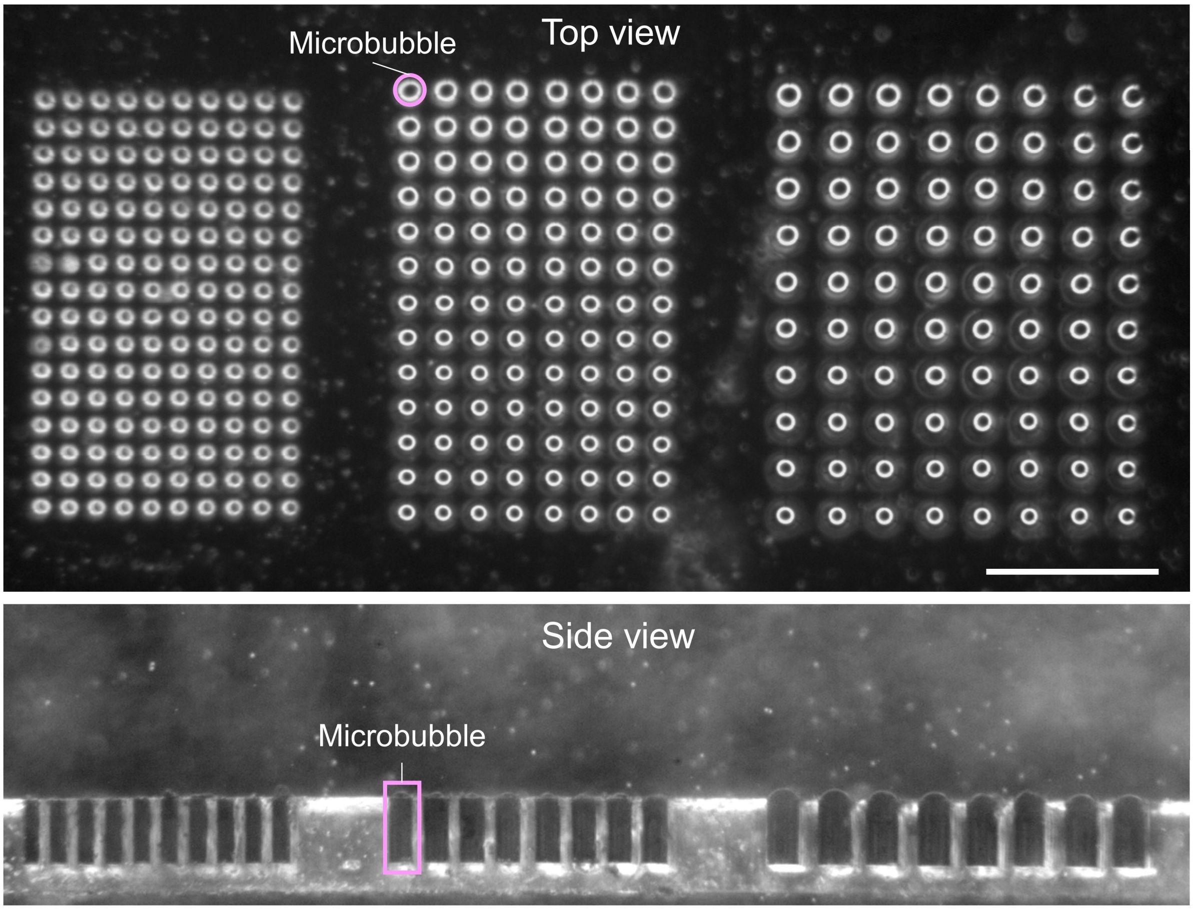


**Supplementary Fig. 13 | Microscopic top and side views of microbubble arrays.** The images illustrate the structure and arrangement of microbubbles. It is notable that each cavity trapped only a single bubble. From left to right, the diameters of the microbubble arrays are 20 µm, 30 µm, and 40 µm, respectively. The trapped microbubbles within cavities are marked with a pink circle (top) and a rectangle (bottom). Scale bar: 200 µm.

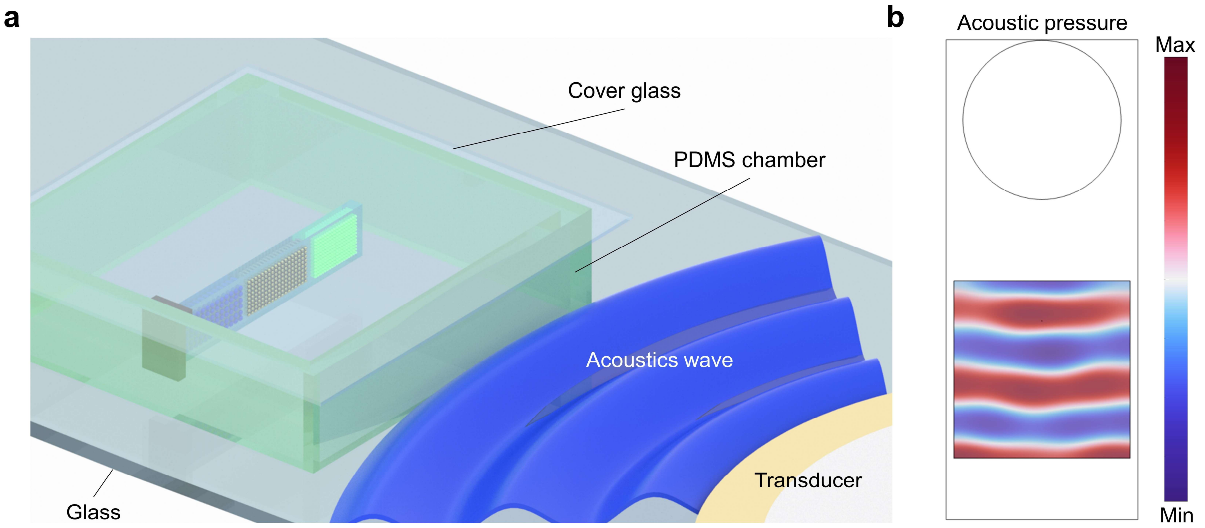


**Supplementary Fig. 14 | Experimental setup for microscope characterization of microbubble arrays and corresponding acoustic pressure field simulation. a**, Experimental setup. A transducer is affixed to the glass slide to generate sound waves and a PDMS chamber is placed in front of the transducer. The artificial muscle was suspended in the center of the chamber with one end clamped to the sidewall and the other end left free. The chamber is filled with deionized water and sealed with a cover glass. **b**, Simulation of the acoustic pressure distribution in the chamber.

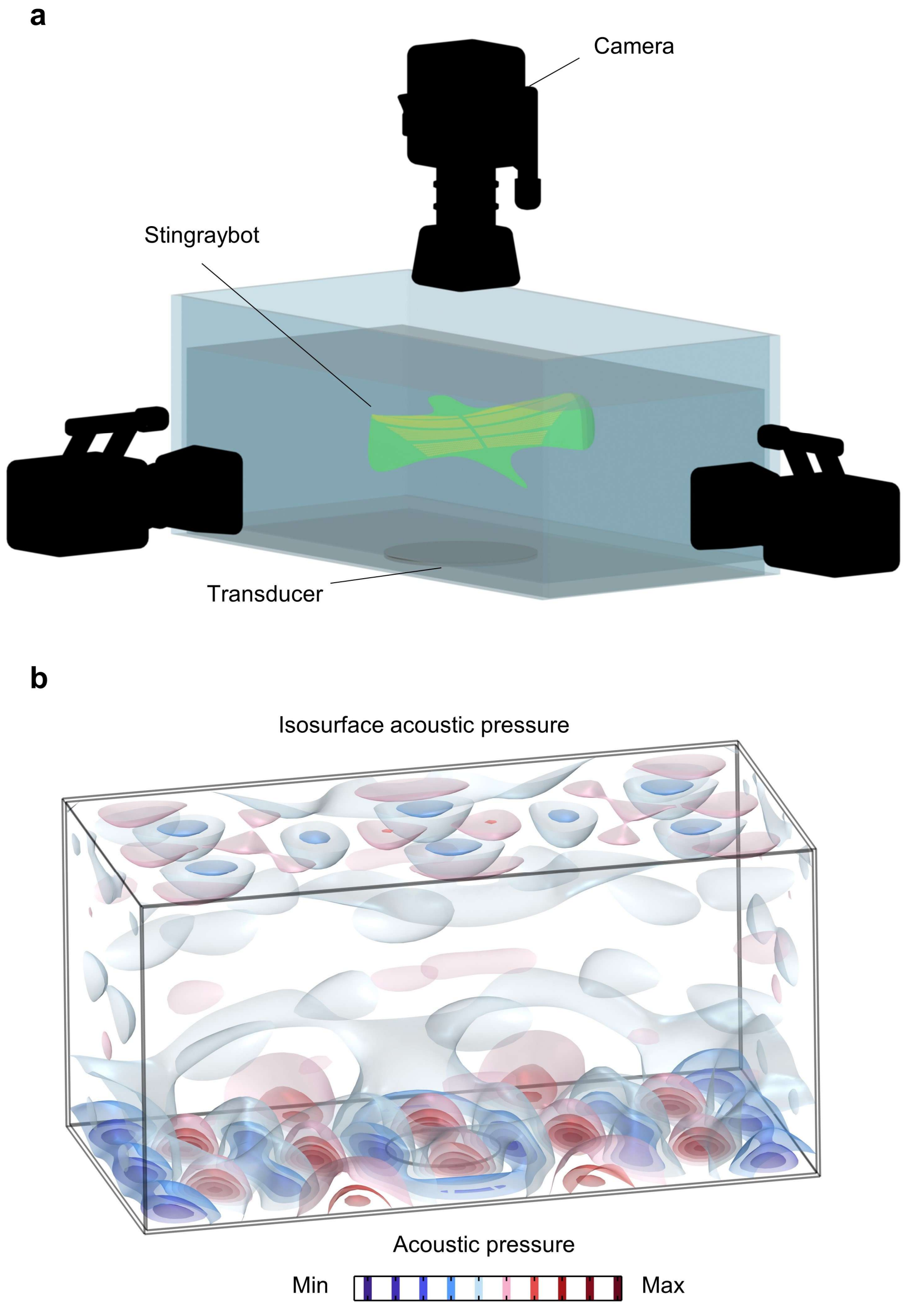


**Supplementary Fig. 15 | Experimental setup for artificial muscle actuation and corresponding acoustic pressure field isosurface simulation**. **a**, Experimental setup. The acoustic tank is filled with deionized water. Three cameras are positioned at the top, left, and front to capture the motion of the bioinspired stingraybot. Transducers are affixed to the bottom of the tank's wall. **b**, Simulation of acoustic pressure isosurface within the tank, where the pressure demonstrates no obvious pressure nodes inside the tank.

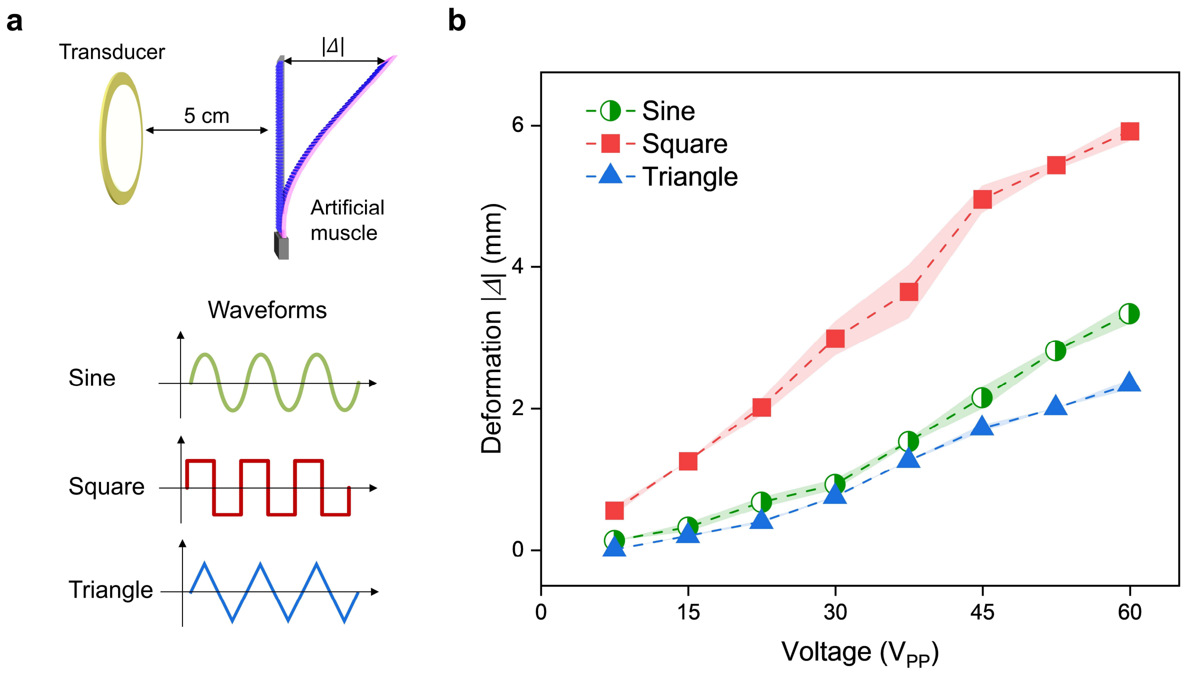


**Supplementary Fig. 16 | Deformation of the artificial muscle under different waveform inputs as a function of applied voltage**. **a**, Experimental setup and the waveforms of the excitation signal. The distances between the transducer and the uniform-size microbubble array (12 µm × 50 µm) artificial muscle is 5 cm, the excitation frequency is set to 95.5 kHz. **b**, The deformation response for three waveform types. Shaded areas represent the standard deviation of deformation for each waveform type, indicating variability across measurements.

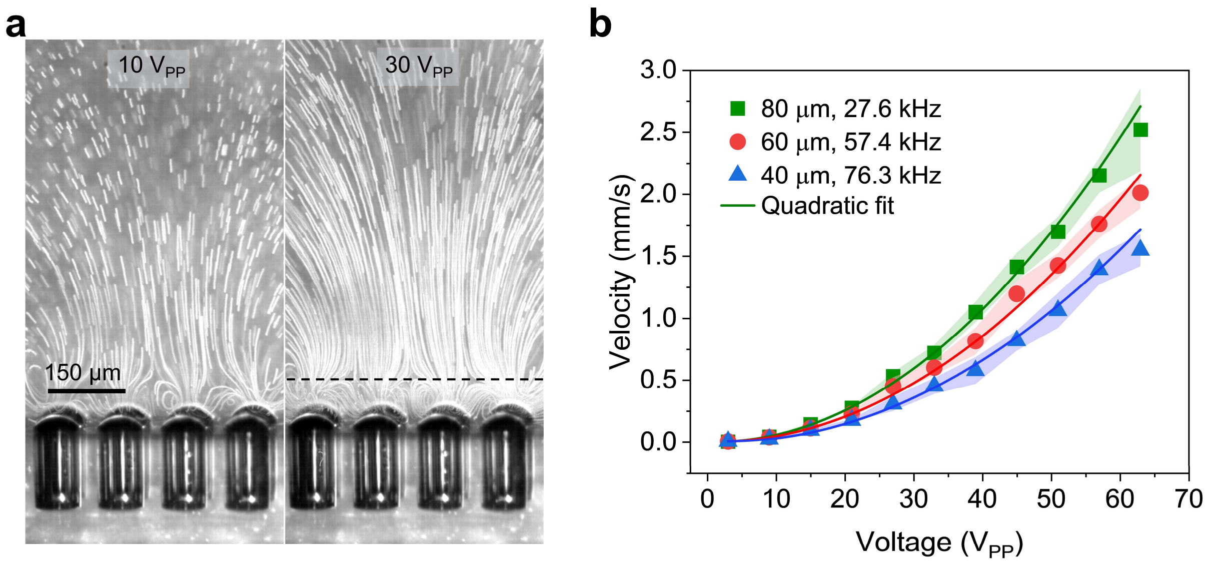


**Supplementary Fig. 17 | Measurement of microstreaming velocity by PIV. a**, Microstreaming generated by a 4×4 80 µm × 150 µm microbubble array under an excitation frequency of 27.6 kHz and two different voltages of 10 $V_{PP}$ (left panel) and 30 $V_{PP}$ (right panel). The black dotted line denotes the measurement position of the microstreaming velocity, which is 80 µm away from the surface. **b**, Plot of microstreaming velocity versus ultrasound excitation voltage respectively measured by 4×4 microbubble arrays with three different sizes (40 µm, 60 µm, and 80 µm). The solid lines are the quadratic fitting results.

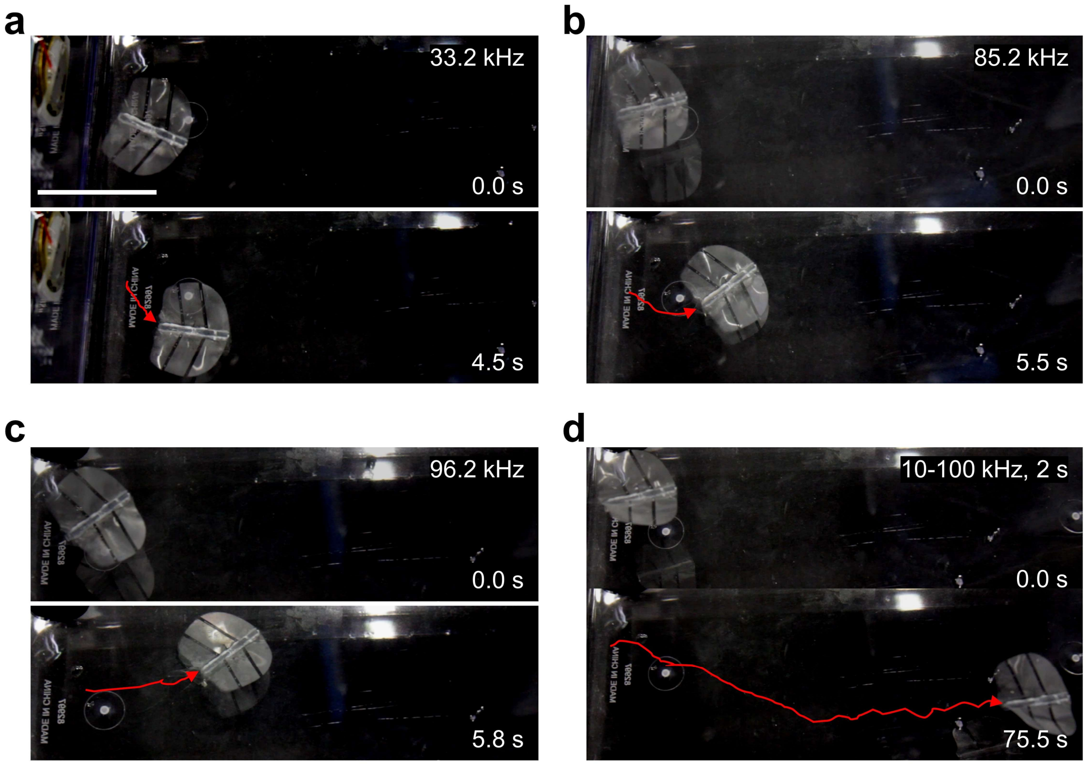


**Supplementary Fig. 18 | Swimming distances of a stingraybot under different excitation conditions**. **a–c**, Stingraybots with microbubble diameters of 12 µm, 16 µm, and 66 µm swim under continuous-frequency excitation at 96.2 kHz, 85.2 kHz, and 33.2 kHz, respectively. **d**, Swimming of the stingraybot under frequency-swept excitation (10–100 kHz over 2 s). All excitations are driven at 60 $V_{PP}$. Scale bar: 5 cm.

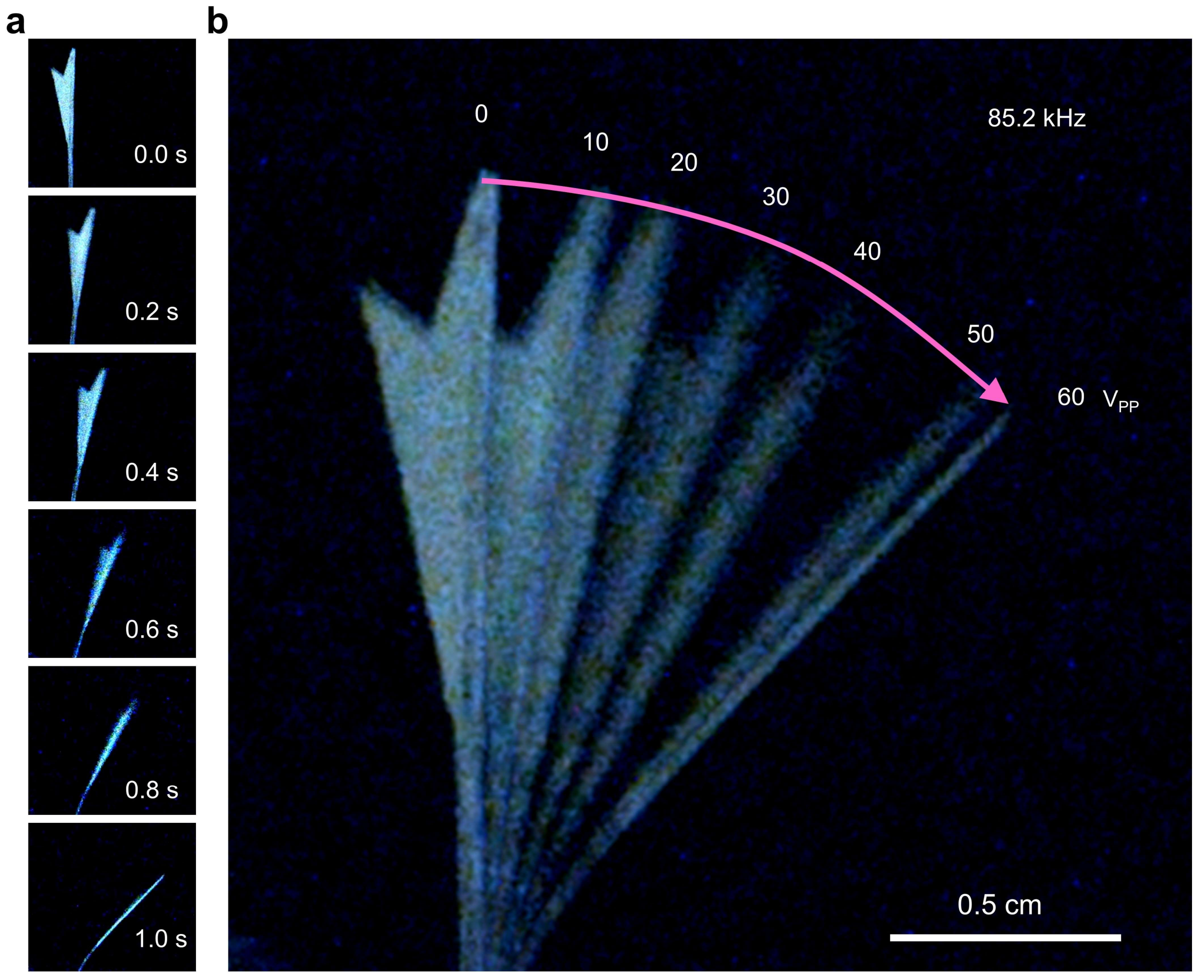


**Supplementary Fig. 19 | Deformation of a robotic shark tail with the uniform-size microbubble array artificial muscle. a,** Time-lapse deformation of the shark tail over a 1 s time interval at ultrasound excitation of 85.2 kHz and 60 $V_{PP}$ with uniform-size microbubble array (16 µm × 50 µm). **b,** Superpositioned deformation of the shark tail with voltage increased from 0 to 60 $V_{PP}$, while maintaining a constant frequency of 85.2 kHz. The magenta arrow denotes the deformation direction with the microbubble array on the left side of the shark tail.

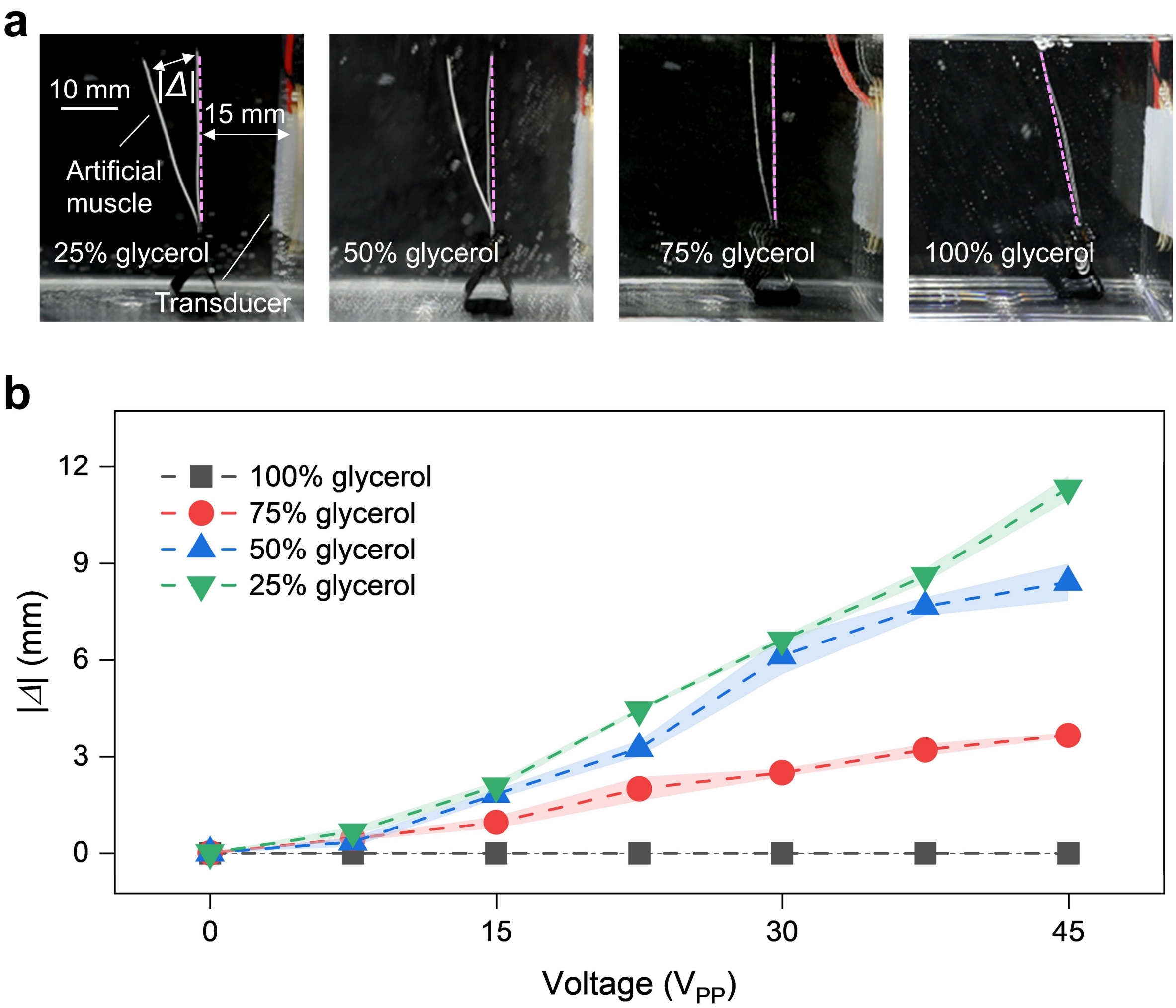


**Supplementary Fig. 20 | Deformation of an artificial muscle in liquid within different ratios of glycerol. a**, Experimental results for a uniform-size microbubble array artificial muscle (12 μm × 50 μm) under varying glycerol-water ratios. The distance between the transducer and the artificial muscle is 15 mm. Pink dashed lines indicate the initial position of the artificial muscle. Excitation was applied at 96.2 kHz with a voltage of 45 $V_{PP}$. **b**, Deformation ($|\Delta|$) of the artificial muscle as a function of the excitation amplitude for different glycerol-water ratios.

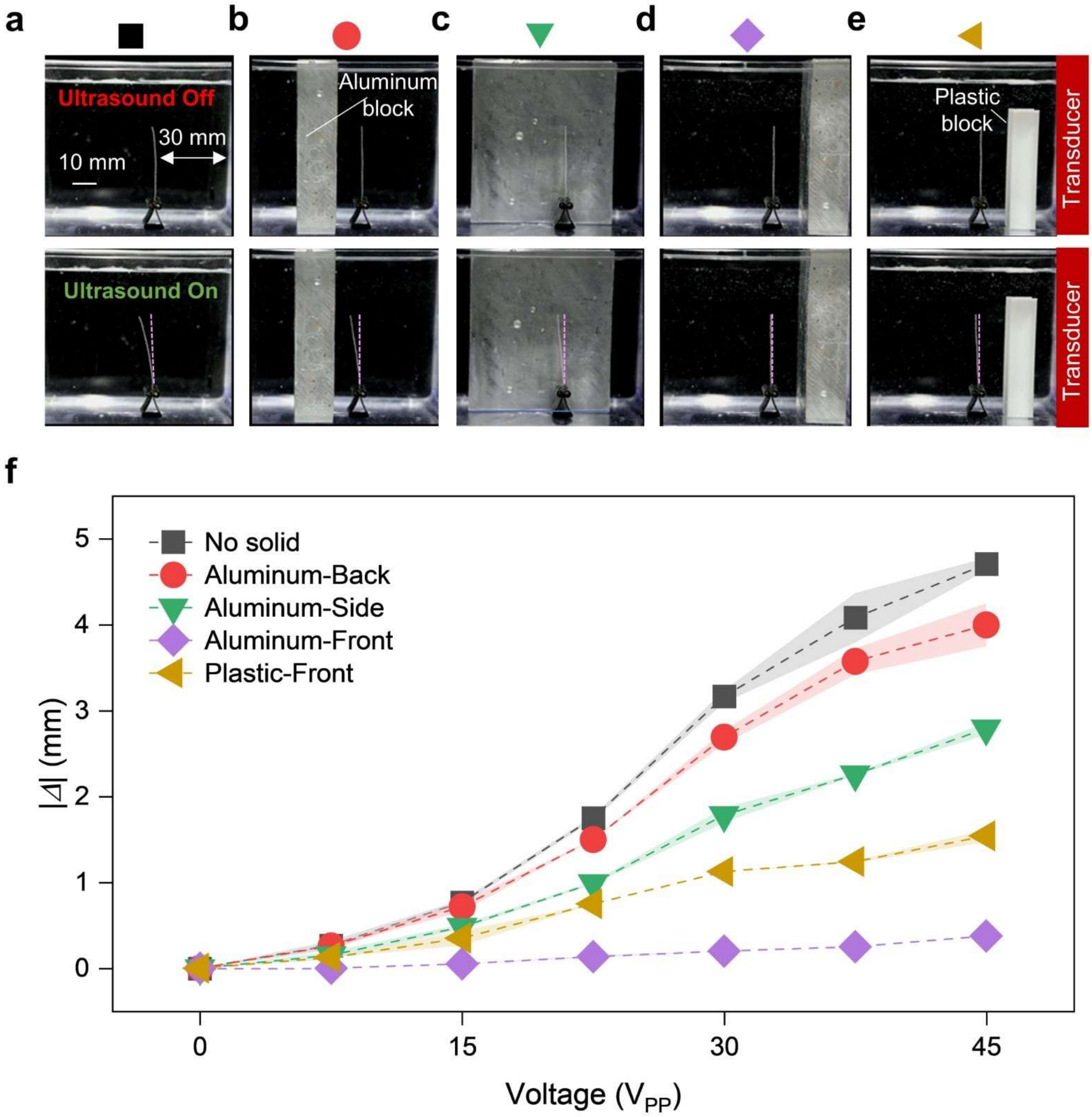


**Supplementary Fig. 21 | Effect of surrounding solid media on artificial muscle deformation under ultrasound excitation. a-e**, Experimental setup showing initial (pink dashed lines) and deformed positions of a uniform-size microbubble array artificial muscle (12 μm × 50 μm) with various solid-media configurations: no solid (black square), aluminum at the back (red circle), side (green triangle), and front (purple diamond), and plastic at the front (yellow triangle). All solids are placed 10 mm from the muscle, with the transducer 30 mm away. Excitation was applied at 95.5 kHz with a voltage of 45 $V_{PP}$. **f**, Deformation ($|\varDelta|$) of the artificial muscle as a function of the excitation voltage for different solid media types and positions.

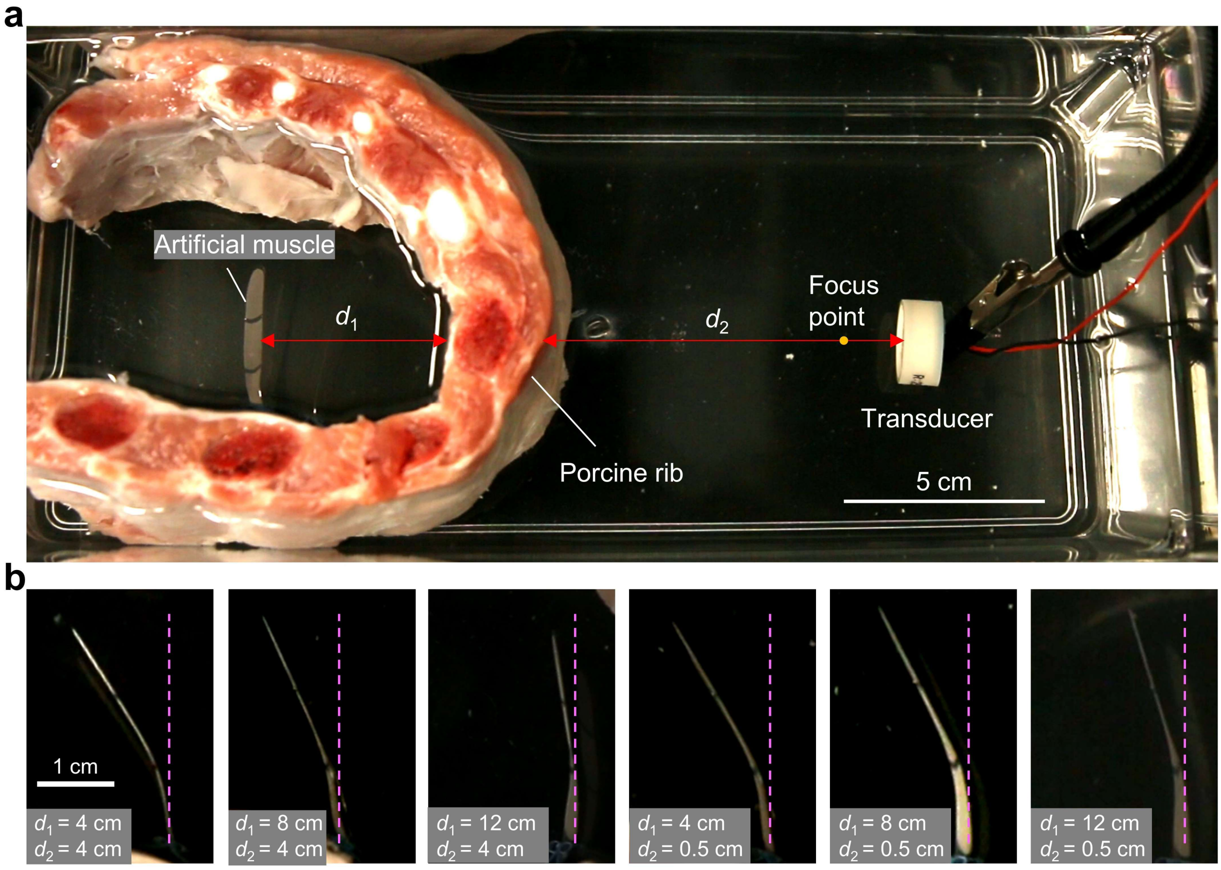


**Supplementary Fig. 22 | Actuation of the artificial muscle behind porcine ribs. a**, Experimental setup showing the artificial muscle positioned at distance $d_1$ behind a porcine rib segment, with a HIFU transducer placed at distance $d_2$ on the opposite side. The transducer's focal point is indicated (2 cm), but actuation was performed in the far field. **b**, Deformation of the artificial muscle under ultrasound excitation (2 MHz, 60 $V_{PP}$) at different positions. Left to right: $d_1$ = 4, 8, 12 cm (top row: $d_2$ = 4 cm); $d_1$ = 4, 8, 12 cm (bottom row: $d_2$ = 0.5 cm). The actuator exhibits clear deformation even through bone, demonstrating reliable performance across clinically relevant distances. Pink dashed lines indicate the initial position of the artificial muscle.

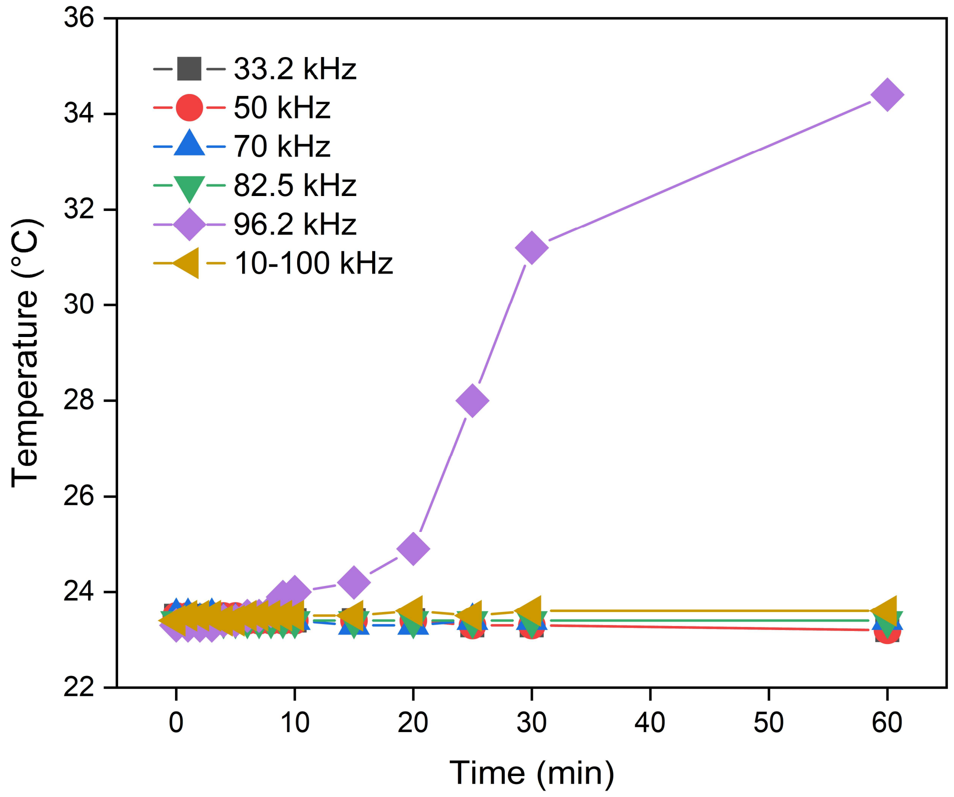


**Supplementary Fig. 23 | Temperature variation for different ultrasound frequencies versus time.** The temperature remains relatively stable for 33.2 kHz, 50 kHz, 70 kHz, 82.5 kHz and 10-100 kHz with 2 seconds duration time, while 96.2 kHz shows a significant increase in temperature as time progresses. The temperature changes were measured using a temperature probe positioned ~1 mm away from the piezo transducer.

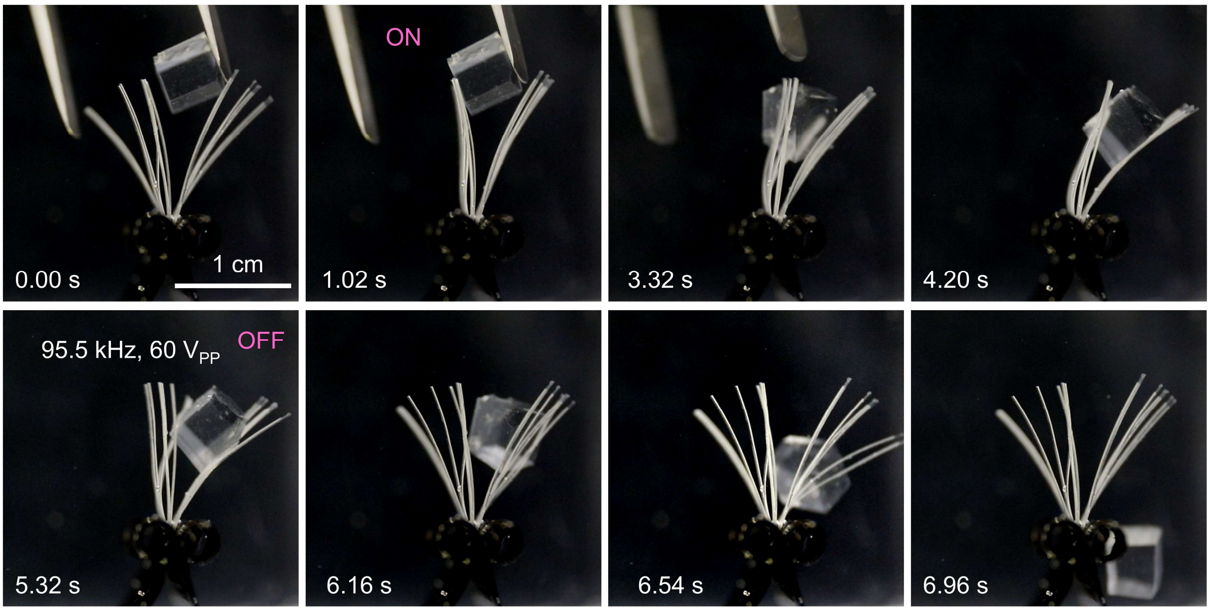


**Supplementary Fig. 24 | A microbubble-array soft gripper captured a 35 times heavier PDMS block.** The gripper is 7 mg and the PDMS block is 250 mg. The soft gripper comprises of 10 uniform-sized microbubble-array artificial muscle, each of them housed >20000 microbubbles with the diameter of 12 μm. The ultrasound excitation frequency and voltage are 95.5 kHz and 60 $V_{PP}$.

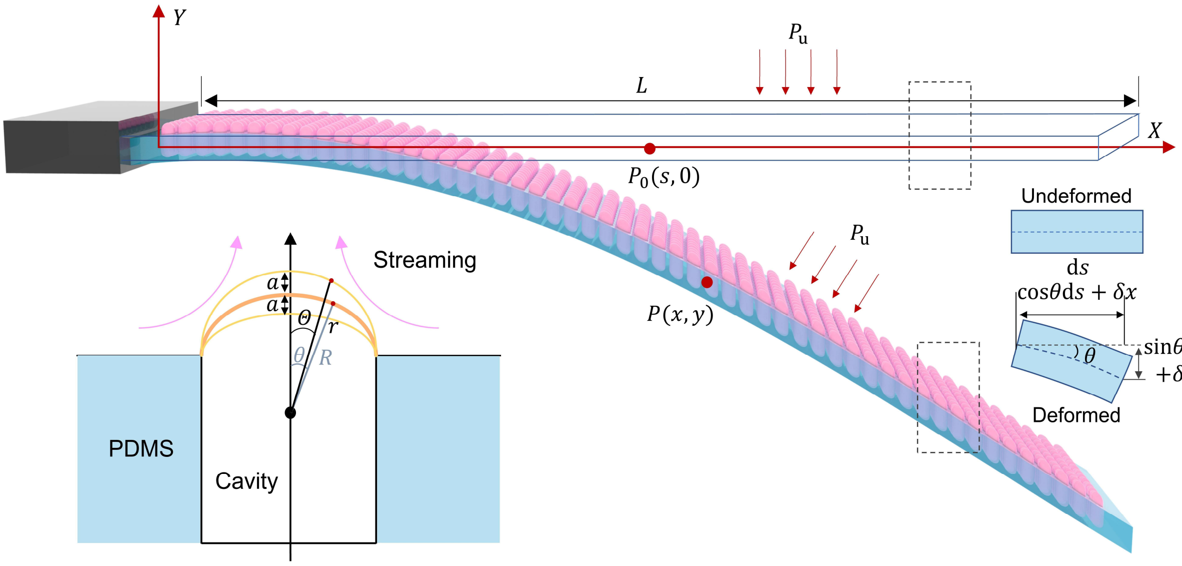


**Supplementary Fig. 25 | Modeling of the uniform-size microbubble array artificial muscle**. Schematic of a cantilever-like artificial muscle composed of a uniform-size microbubble array, transitioning from the undeformed to the deformed state under microbubble streaming actuation. The inset illustrates the parametrization of an oscillating microbubble. For detailed bubble and muscle parameters, see **Supplementary Notes**.

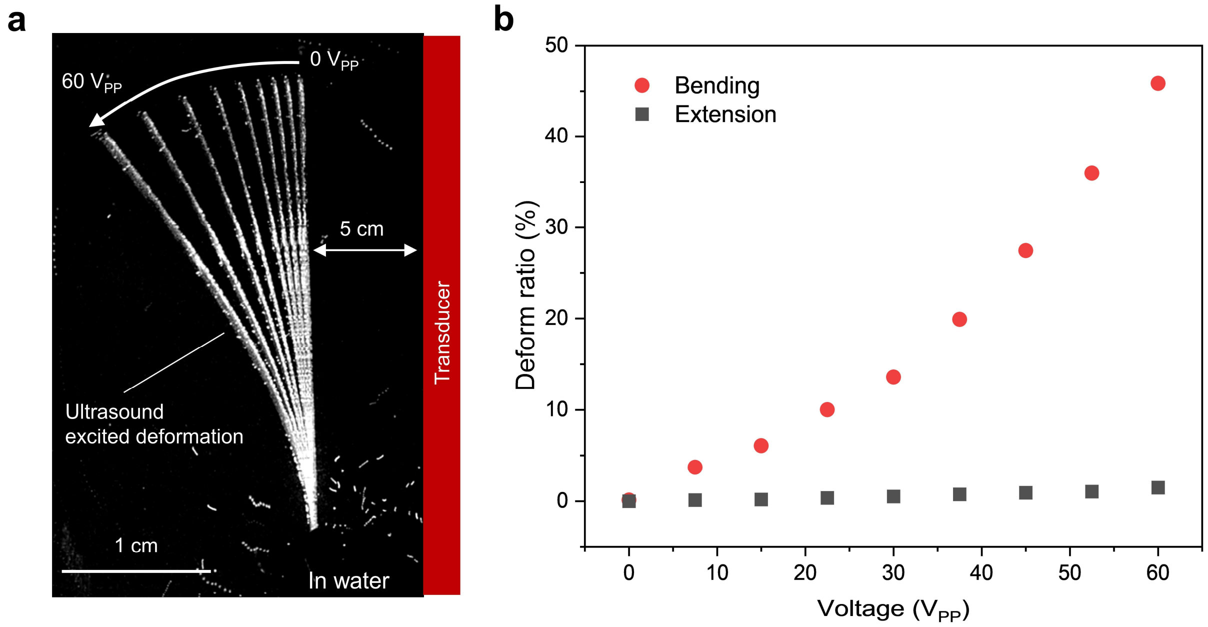


**Supplementary Fig. 26 | Deformation of an artificial muscle under ultrasound excitation. a,** Superpositioned deformation of the uniform-size microbubble array artificial muscle with voltage increased from 0 to 60 $V_{PP}$, where the excitation frequency was 27.6 kHz. **b**, The bending ratio (red dots) and extension ratio (black dots) along the long axis of the artificial muscle at different excitation voltages, where the measured extension ratio remains below 1.5%.

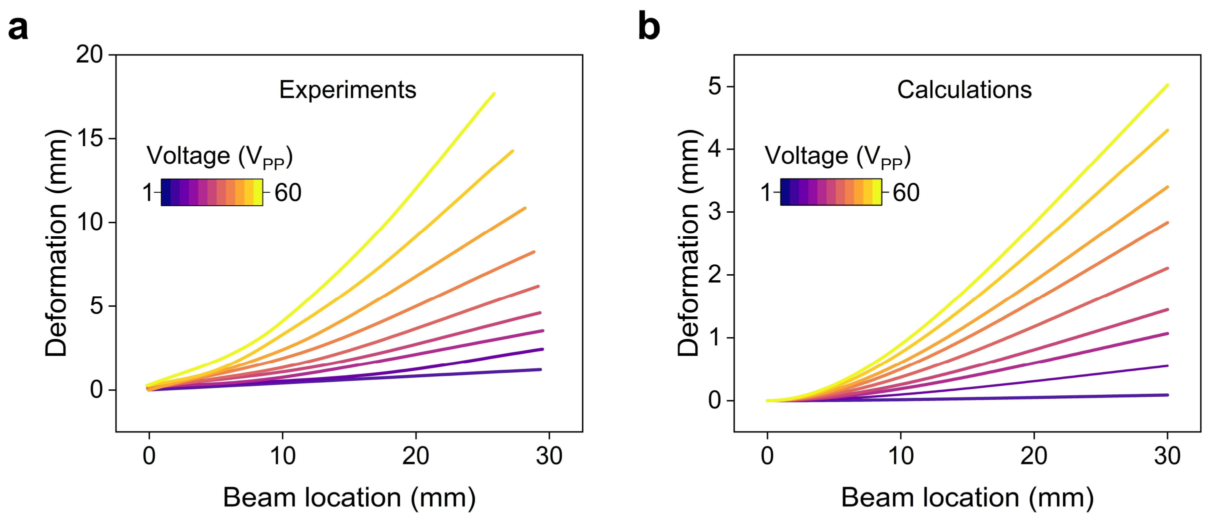


**Supplementary Fig. 27 | Comparison of artificial muscle deformation between experimental results and analytical calculations. a**, Experimentally observed deformed muscle shapes under voltages ranging from 1 to 60 $V_{PP}$, extracted from **Supplementary Fig. 6a**. **b**, Corresponding analytical predictions obtained from Eqs. (S9), (S21), and (S22), using force intensities obtained from experimentally measured velocities (**Extended Data Fig. 3b**). Both experimental and analytical results correspond to a beam measuring 30 mm × 5 mm × 0.2 mm, with a Young's modulus of 1.2 MPa for the PDMS material.

**Supplementary Table 1**. Gripper performance data across actuation mechanisms: response time and grip ability

| Category | Response time (s) | Grip ability (Object/Gripper) | Ref. | Ref. Resources |
|---|---|---|---|---|
| **Acoustic** | 0.1 | 0.5 | Our work | Fig. 3a |
| **Light** | 0.2 | 0.2 | [S5] OM Wani (2017) | Fig. 4d |
| | 60 | 0.33 | [S6] C Ma (2016) | Fig. 4 |
| | 35 | 0.16 | [S7] X Zhang (2021) | Fig. 3 |
| **Thermal** | 600 | 1 | [S8] W Wei (2018) | Fig. 5 |
| | 552 | 0.833 | [S9] AK Mishra (2020) | Fig. 5 |
| | 15 | 0.786 | [S10] TG Leong (2009) | Fig. 4 |
| | 8 | 0.63 | [S11] Y Roh (2021) | Fig. 3 |
| | 1.8 | 1 | [S12] H Shahsavan (2017) | Fig. 4 |
| | 1.2 | 0.2 | [S13] S Li (2021) | Fig. 4 |
| | 4 | 0.23 | [S14] JC Breger (2015) | Fig. 5 |
| | 63 | 0.8 | [S15] C Yao (2015) | Fig. 10 |
| **Magnetic** | 0.035 | 0.8 | [S16] X Wang (2020) | Fig. 1 |
| | 0.02 | 0.45 | [S17] M Lukas (2024) | Fig. 4 |
| | 22 | 0.3 | [S18] H Gu (2023) | Fig. 5 |
| | 1 | 0.28 | [S19] Y Jiang (2023) | Fig. 3 |
| **Pneumatic** | 0.62 | 0.37 | [S20] Z Zhang (2023) | Fig. 3 |
| | 40 | 0.15 | [S21] X Guo (2024) | Fig. 3 |
| | 0.065 | 0.6 | [S22] WY Choi (2025) | Fig. 3 |

**Supplementary Table 2**. Comparison data of force-to-weight ratios of artificial muscles versus dimensions for various actuation mechanisms.

| Category | Size (mm) | Force-to-weight ratio (N/N) | Ref. | Ref. Resources |
|---|---|---|---|---|
| **Acoustic** | 5 | 35 | Our work | Supplementary Fig. S24 |
| **Chemical** | 15 | 2.61 | [S23] X Yang (2020) | Movie. S7 |
| **Dielectric** | 112 | 1 | [S24] SD Gravert (2024) | Fig. 1C |
| | 100 | 8 | [S25] Y Xu (2025) | Fig. 5D |
| | 20 | 127 | [S26] ID Sîrbu (2021) | Fig. S10 |
| **Light** | 0.7 | 0.4 | [S27] Y Wang (2022) | Fig. S11 |
| | 80 | 268 | [S28] W Ai (2024) | Fig. S4 |
| | 33.7 | 114.9 | [S29] H Liu (2022) | Fig. 4F |
| **Magnetic** | 60 | 1004 | [S30] M Seong (2024) | Fig. S11b |
| | 30 | 64 | [S31] Q Ze (2022) | Fig. 3D |
| **Pneumatic** | 100 | 46.4 | [S32] Z Zhang (2019) | Fig. 19 |
| | 70 | 34 | [S33] S Zhuo (2020) | Fig. 4 |
| | 120 | 5.9 | [S34] SM Mirvakili (2020) | Fig. 6 |
| **Thermal** | 10 | 10 | [S35] BE Schubert (2013) | Fig. 1B |
| | 10 | 450 | [S36] Y Yao (2024) | Fig. S25 |

**Supplementary Table 3**. Comparison data of relative swimming speeds of swimmers relative to their body length for various actuation mechanisms.

| Category | Body length (mm) | Relative speed (BL/s) | Ref. | Ref. Resources |
|---|---|---|---|---|
| **Acoustic** | 50 | 0.8 | Our work | Fig. 4b |
| | 0.07 | 50 | [42] D Ahmed (2015) | Fig. 3 |
| | 0.0074 | 350 | [43] L Ren (2019) | Fig. 2D |
| | 0.026 | 90 | [44] A Aghakhani(2020) | Fig. 3D |
| | 0.02 | 320 | [S37] M Nima (2025) | Fig. 2D |
| | 0.03 | 68 | [39] H Han(2024) | Fig. 2N |
| | 0.07 | 55 | [S38] M Nima (2023) | Fig. 4B |
| | 0.18 | 6.7 | [S39] M Kaynak (2017) | Fig. 4 |
| | 0.3 | 0.017 | [S40] T Qiu (2014) | Fig. 3a |
| | 0.26 | 10 | [37] C Dillinger (2021) | Fig. 4b |
| **Biohybrid** | 14 | 1.07 | [S41] LK Yong (2017) | Movie. S11 |
| | 16.3 | 0.196 | [S42] SJ Park (2016) | Fig. 4 |
| | 0.27 | 3 | [S43] M Guix (2021) | Abstract |
| | 3.2 | 0.00017 | [S44] O Aydin (2019) | Fig. 4D |
| | 0.0025 | 0.001 | [S45] W. C. Drennan (2025) | Abstract |
| **Chemical** | 400 | 0.026 | [S46] CA Aubin (2019) | Fig. 5D |
| | 0.003 | 3.3 | [S47] RD Baker (2019) | Fig. 6B |
| | 0.004 | 2 | [S48] S Ketzetzi (2022) | Fig. 2F |
| | 4.3 | 30 | [S49] S Arnaboldi (2021) | Video 6-8 |
| | 0.01 | 100 | [S50] W Gao (2012) | Fig. 3 |
| | 0.008 | 39.75 | [S51] L Wang (2016) | Fig. 4 |
| | 0.1 | 3.2 | [S52] L Li (2014) | Fig. 5 |
| **Electric** | 0.05 | 0.09 | [S53] Y Alapan (2019) | Fig. 3d |
| | 90 | 1.1 | [S54] Z Ye (2025) | Movie. S2 |
| | 70 | 0.65 | [S55] JD Cortazar (2025) | Abstract |
| | 80 | 0.11 | [S56] H Yang (2025) | Fig. 8G |
| **Light** | 0.0015 | 14.9 | [S57] V Sridhar (2022) | Abstract |
| | 1.23 | 0.0023 | [S58] S Palagi (2016) | Fig. 3c |
| | 0.06 | 0.0167 | [S59] MZ Miskin (2020) | Fig. 4b, c |
| | 10 | 0.337 | [S60] C Yin (2021) | Fig. 3 |
| | 5 | 2.6 | [S61] K Hou (2021) | Fig. 3C |
| | 28.2 | 0.0083 | [S62] Y Zhao (2019) | Fig. 5C |
| **Magnetic** | 4 | 5 | [S63] W Hu (2018) | Fig. 2 |
| | 6 | 1.2 | [S64] Z Ren (2019) | Fig. 1d |
| | 13 | 1 | [S65] GZ Lum (2016) | Movie. S3 |
| | 0.009 | 14.4 | [S66] S Tottori (2012) | Video S3 |
| | 0.015 | 4 | [S67] X Wang (2018) | Fig. 2 |
| **Pressure** | 528 | 0.36 | [S68] DQ Nguyen (2022) | Abstract |
| | 470 | 0.5 | [S69] K Katzschmann (2018) | Fig. 3 |
| | 140 | 3.74 | [S70] Y Chi (2022) | Fig. 3 |

**Legends for Supplementary Videos**

**Video 1. Selectively excitation of variable-size microbubble arrays.**

Upward microstreaming generated by (40 μm, 60 μm, and 80 μm) × 150 μm microbubble arrays under ultrasound frequencies of 27.6 kHz, 57.4 kHz, and 76.3 kHz, respectively, at an excitation voltage of 24 $V_{PP}$. Rectangles indicate the excited regions.

**Video 2. Deformations of a uniform-size microbubble array artificial muscle with varying transducer positions.**

The transducer was respectively positioned with four distinct orientations: top left panel: directly facing the microbubble-embedded side, top right panel: opposite to it, and perpendicular to the array's left (low left panel) and right (low right panel) sides of the artificial muscle. The side with the microbubble array is marked by a red rectangle. The ultrasound excitation was 80.5 kHz and 60 $V_{PP}$.

**Video 3. Static deformation of a variable-size microbubble array artificial muscle under continuous-frequency ultrasound actuation.**

The diameters of the microbubbles in the three different arrays, from top to bottom, are 12 μm, 16 μm, and 66 μm, with corresponding excitation frequencies of 96.5 kHz, 82.3 kHz, and 33.2 kHz. The excitation voltage was 60 $V_{PP}$.

**Video 4. Undulatory deformation of a variable-size microbubble array artificial muscle under sweeping-frequency ultrasound actuation.**

The artificial muscle undergoes undulatory motion as the excitation frequency was swept from 20 to 90 kHz, with a sweep time of 1.2 seconds each cycle at 60 $V_{PP}$.

**Video 5. Operation of the soft gripper at arbitrary locations.**

The gripper consisting of six to ten uniform-size microbubble array artificial muscles (12 μm × 50 μm), each measuring 10 mm × 0.7 mm × 0.08 mm, was held by a tweezer and functions at arbitrary locations within the acoustic tank at 95.5 kHz and 60 $V_{PP}$.

**Video 6. Soft gripper handling delicate zebrafish larvae.**

The gripper was composed of six to ten uniform-size microbubble array artificial muscle tentacles (12 μm × 50 μm), each measuring 10 mm × 0.7 mm × 0.08 mm. When submerged in water, each tentacle contained ~20,000 microbubbles. The ultrasound excitation was 95.5 kHz and 60 $V_{PP}$. Upon ultrasound stimulation, the tentacles grasped the motile larva; once the acoustic excitation was deactivated, the larva readily swarm away.

**Video 7. Actuation of an almond.**

Rotating of an almond using a conformable robotic skin composed of a uniform-size microbubble array (12 μm × 50 μm), operated at an excitation voltage of 60 $V_{PP}$ and a frequency of 95.5 kHz.

**Video 8. Actuation of a piece of grass.**

Closing grass blades using a conformable robotic skin composed of a uniform-size microbubble array (12 μm × 50 μm), operated at an excitation voltage of 60 $V_{PP}$ and a frequency of 95.5 kHz.

**Video 9. Attachment of the robotic patch to a porcine heart.**

The artificial muscle was positioned between an ultrasound transducer (operating at 96 kHz and 60 $V_{PP}$) and the heart, with a separation of approximately 2.5 cm. It was released from below using tweezers.

**Video 10. Multi-modal deformation of the shape transformer.**

Multi-modal shape transformation of a circular functional surface as the frequency was swept from 10 to 100 kHz. Each cycle had a sweeping time of 2 seconds, and the input voltage was 60 $V_{PP}$.

**Video 11. Deployment of the encapsulated microbubble-array artificial muscle in a porcine stomach and bladder.**

Within the porcine stomach and bladder, the encapsulated artificial muscle released and attaches to the inner wall.

**Video 12. Swimming of the stingraybot.**

Fins of the stingraybot were constructed using variable-size microbubble array artificial muscles (12 μm × 50 μm, 16 μm × 50 μm, and 66 μm × 50 μm), arranged vertically from head to tail. The actuation was achieved by sweeping-frequency ultrasound excitation (30–90 kHz, 2 s duration, 60 $V_{PP}$).

**Video 13. Undulatory motion and locomotion of the stingraybot in a porcine stomach.**

The stingraybot exhibits undulatory motion with a fixed tail and swims untethered.

**Video 14. Locomotion of a pre-folded, wheel-like artificial muscle driven by ultrasound inside a porcine stomach and intestine.**

The 30 mm × 5 mm × 80 μm muscle, composed of variable-size microbubble arrays (12, 16, and 66 μm in diameter; 50 μm in depth), was pre-folded into wheel-like, propels along the stomach surface under sweeping-frequency ultrasound excitation (30–100 kHz, 2 s, 60 $V_{PP}$).

**Video 15. Acoustic streaming under different ultrasound excitation voltages.**

The microbubble array contains 4×4 microbubbles with a diameter of 80 μm. The left and right videos correspond to 10 $V_{PP}$ and 30 $V_{PP}$, respectively, at a frequency of 27.6 kHz. 6 μm microparticles were used as tracer particles.

**Video 16. Response of variable-size microbubble arrays under sweeping-frequency ultrasound excitation.**

The actuation was achieved by sweeping the frequency within the range of 10-90 kHz, with a sweep time of 4 seconds each cycle at 30 $V_{PP}$.

**Video 17. Comparison of the deformation of a uniform-size microbubble array artificial muscle without and with microbubbles.**

Without microbubbles, the excitation frequency was in the range of 1—100 kHz, with a sweeping time of 3 seconds per cycle at 60 $V_{PP}$. With 40 μm microbubbles, the excitation frequency and voltage were 9.5 kHz and 60 $V_{PP}$.

**Video 18. Comparison between the stingraybot fin deformations with/without microbubbles.**

The left panel shows the fin without embedded microbubbles, while the right panel includes the microbubble arrays. Actuation was achieved using a sweeping-frequency ultrasound excitation (60–100 kHz, 1.5 s duration, 60 $V_{PP}$). In both experiments, the stingraybot was fixed to the bottom of the acoustic tank.
